# Glycometabolic, inflammatory and exocrine plasma proteins predict cancer in individuals with cardiovascular disease

**DOI:** 10.64898/2026.09.09.750392

**Authors:** Marine Fidelle, David Boulate, Thao-Nguyen Pham, Antoine Giboudot, Louis Picq, Pamela Abdayem, Aline Renneville, Pauline Pradère, Caroline Caramella, Sylvère Durand, Fanny Aprahamian, Thibault Dayris, Marc Deloger, Nathalie Droin, Océane Hache, Magalie Martineau, Thierry Walzer, Olaf Mercier, Tatiana Kuznetsova, Laurence Zitvogel, Nikos Paragios, Guido Kroemer

**Author notes:** Shared first authors (Marine Fidelle, David Boulate).

## Abstract

Cancer development is preceded by systemic immune, metabolic and tissue perturbation. We asked whether a minimal circulating-protein signature anticipates incident cancer in patients with atheromatous cardiovascular disease (ACVD) exposed to tobacco. Discovery used a nested case–control set drawn from the longitudinal FLEMENGHO cohort (n=156; 38 incident lung cancers). Multi-omic profiling returned a two-step classifier: four proteins, smoking status, personal cancer history. Preprocessing constants, classifiers and both thresholds were fixed before transfer to PREVALUNG (n=397; 58 incident cancers, 26 of them lung). There, low O-GlcNAcase (OGA) with low interleukin-6 (IL-6) defined a very-low-risk stratum holding 1 of 58 cancers (sensitivity 98.3%, negative predictive value 98.8%); low chymotrypsin C (CTRC) with high β-microseminoprotein (MSMB) defined a high-risk stratum in which 15 of 28 participants developed cancer. No additional omic feature — metabolomic, immunophenotypic, clonal-haematopoietic or metagenomic — was retained as a reproducible improvement to the classifier under our selection framework. Discrimination was greater for cancers diagnosed ≥12 months after sampling than for near-term cancers (AUC 0.766, 95% CI 0.671–0.857, versus 0.589, 0.485–0.692), arguing against performance being driven principally by occult disease. These estimates are conditional on the cancer frequency of this population. They require prospective confirmation before any imaging schedule is changed.

## Introduction

The practical objective of cancer screening campaigns is to identify people in whom an actionable premalignant or malignant process is incipient and hence curable. For example, low-dose computed tomography (LDCT) shifts diagnosis towards stage I and reduces lung-cancer mortality: the National Lung Screening Trial (NLST) reported a 20% relative reduction versus chest radiography, and the Dutch-Belgian NELSON trial found a reduction of 24% mortality risk among screened men at 10 years versus unscreened controls.^1,2^ When disease is confined, surgery or stereotactic radiotherapy can be curative; at an even earlier biological phase, risk-adapted surveillance, smoking cessation and prevention or interception strategies may suppress progression before an invasive, metastatic clone is established. Nonetheless, LDCT is a resource-intensive population intervention. Eligibility based mainly on age and cigarette exposure misses some people who will develop cancer, including former smokers outside a quit-time window, lighter smokers and never-smokers, while exposing many people who will not develop cancer to repeated imaging. In the NLST, 96.4% of positive screening results in the first three rounds were false positives.^1^ Most are resolved non-invasively, but indeterminate nodules generate anxiety, follow-up scans and occasional invasive procedures. A blood test should therefore be conceived primarily as a calibrated risk-stratification or triage tool that directs LDCT to those most likely to benefit, rather than as a stand-alone diagnostic replacement for imaging.

Clinical prediction models outperform simple eligibility criteria based on age and cumulative tobacco exposure (pack-years). PLCO_m2012_^3^, LLP^4^ and related models combine smoking intensity, duration and cessation with variables such as chronic obstructive pulmonary disease (COPD) or emphysema, personal history of cancer, family history of lung cancer, body-mass index (BMI), education or deprivation, ethnicity, occupational exposures and, in some models, pneumonia, asbestos exposure or respiratory symptoms.^3–7^ Risk-model selection can identify more future cancers per scan than categorical smoking criteria and is already used to guide some LDCT programs. Clinical models remain unsatisfactory as biological predictors of cancer. Several inputs are correlated, self-reported or imprecisely measured.^7^ Consequently, clinical scores enrich rather than cleanly separate future cases from non-cases, leaving both preventable false-negative exclusions and a large imaging burden.

Prediagnostic biobanks have revealed a blood-based signal that is much broader than clinical scores. Replicated associations include systemic inflammation (C-reactive protein [CRP], interleukin-6 [IL-6], and interleukin-8 [IL-8)), tumor- or airway-associated proteins (including pro-SFTPB, CA125, CEA, CYFRA21-1, CDCP1 and ACBP/DBI), smoking-responsive DNA methylation at AHRR and F2RL3, multi-CpG scores and serum small-RNA profiles.^8–17^ Routine leukocyte counts are inexpensive correlates; directly measured or methylation-derived immune composition suggests higher regulatory T-cell and lower CD8 T-cell proportions before diagnosis.^18–20^ Metabolic studies implicate tryptophan–kynurenine metabolism, vitamin-B6 catabolism, polyamines, acylcarnitines, bilirubin, branched-chain amino acids and histidine.^21–28^ Clonal hematopoiesis of indetermined prognosis (CHIP), particularly larger clones, and expanded mosaic chromosomal alterations add a somatic-genetic record of ageing and exposure.^29,30^ Cotinine, polygenic scores, autoantibodies, cell-free DNA, extracellular vesicles and rare circulating cells have shown varying promise, but their incremental utility for lung cancer risk prediction remains incompletely validated.^31^

The predictive horizon is crucial. A marker that rises one year before diagnosis may be detecting an occult cancer and be useful for near-term LDCT triage, but it is not necessarily a causal marker of long-term susceptibility. Conversely, smoking-related methylation,^32^ CHIP^29,30^ or immune dysregulation^18–20^ detected a decade earlier may report a permissive host state yet offer less immediate localization of the cancer. Another limitation is that many recent successes result from an accumulation of markers. A 14-protein signature predicted incident lung cancer across cohorts, while the 13-protein INTEGRAL-Risk model added age and smoking to improve one- to three-year prediction over PLCO_m2012_.^16,17^ Other approaches combine a four-protein panel with the PLCO_m2012_ clinical score, use a 100-feature cfDNA-methylation classifier, or exploit high-dimensional serum-RNA and metabolomic profiles.^33–36^ Complementary biology can raise discrimination, but every analyte adds cost, pre-analytical sensitivity, missingness, batch drift and recalibration. Complexity is not undesirable in itself; it raises the bar for screening millions of asymptomatic people.

The most useful next-generation strategy would combine a parsimonious clinical baseline with a small, mechanistically diverse biomarker set. Cardiovascular phenotype defines an ascertainable risk stratum among smokers. Prospective studies show that cardiovascular disease (CVD) is associated with increased lung-cancer incidence, including among current smokers; severe coronary artery calcification and progression to dyslipidemia are associated with incident lung cancer.^37–42^ These observations indicate that smokers with ACVD, i.e. with previous myocardial infarction, stroke, atheromatous aortic aneurysm, peripheral arterial stenosis or cardiovascular disease requiring revascularization or surgery, are at higher risk than otherwise comparable smokers without such cardiovascular characteristics. We therefore focused on this high-risk population, seeking to establish a convenient and affordable predictive score based on no more than four circulating proteins. Across the exploration and validation cohorts, we identified two circulating proteins that defined individuals with no or very low risk of future lung-cancer development, as well as two complementary proteins that identified individuals at very high risk. This four-analyte, bidirectional rule-out/rule-in architecture may provide an inexpensive blood-first triage for risk-adapted LDCT.

## Results

### Four circulating proteins stratify individuals at risk of cancer

We employed a prospective discovery–validation strategy designed to identify a minimal, four analyte plasma protein-based signature of asymptomatic or future lung cancer among individuals with atheromatous cardiovascular disease (ACVD) and tobacco exposure in the context of a model that additionally included smoking status and personal cancer history as prespecified clinical covariates (**Fig. 1**). Model development was initially based on 156 participants from the Flemish Study on Environment, Genes and Health Outcomes (FLEMENGHO), a population-based Belgian cohort, including 38 individuals who developed lung cancer after blood collection (**Supplementary Table 1 and Supplementary Fig. 1**).^15,43^ Transfer of the locked cascade used 397 participants from PREVALUNG, a prospective French cohort of current or former smokers with smoking-associated ACVD. In the PREVALUNG study, 58 participants developed cancer, among which 40 were tobacco-related (26 lung cancers, 10 bladder cancers, 4 head and neck cancers) and 18 unrelated (prostate, pancreas, colon, breast…) (**Supplementary Table 2 and Supplementary Fig. 2**).^44^ Samples (serum, plasma, peripheral blood mononuclear cells [PBMC], feces) from these patients collected at the date of inclusion and subjected to serum proteomics (using Olink), metabolomics (by mass spectrometry), immunonomics (by high-dimensional cytometry), CHIP detection and shotgun metagenomics for microbiome profiling (**Fig. 1**).

**Figure 1.**
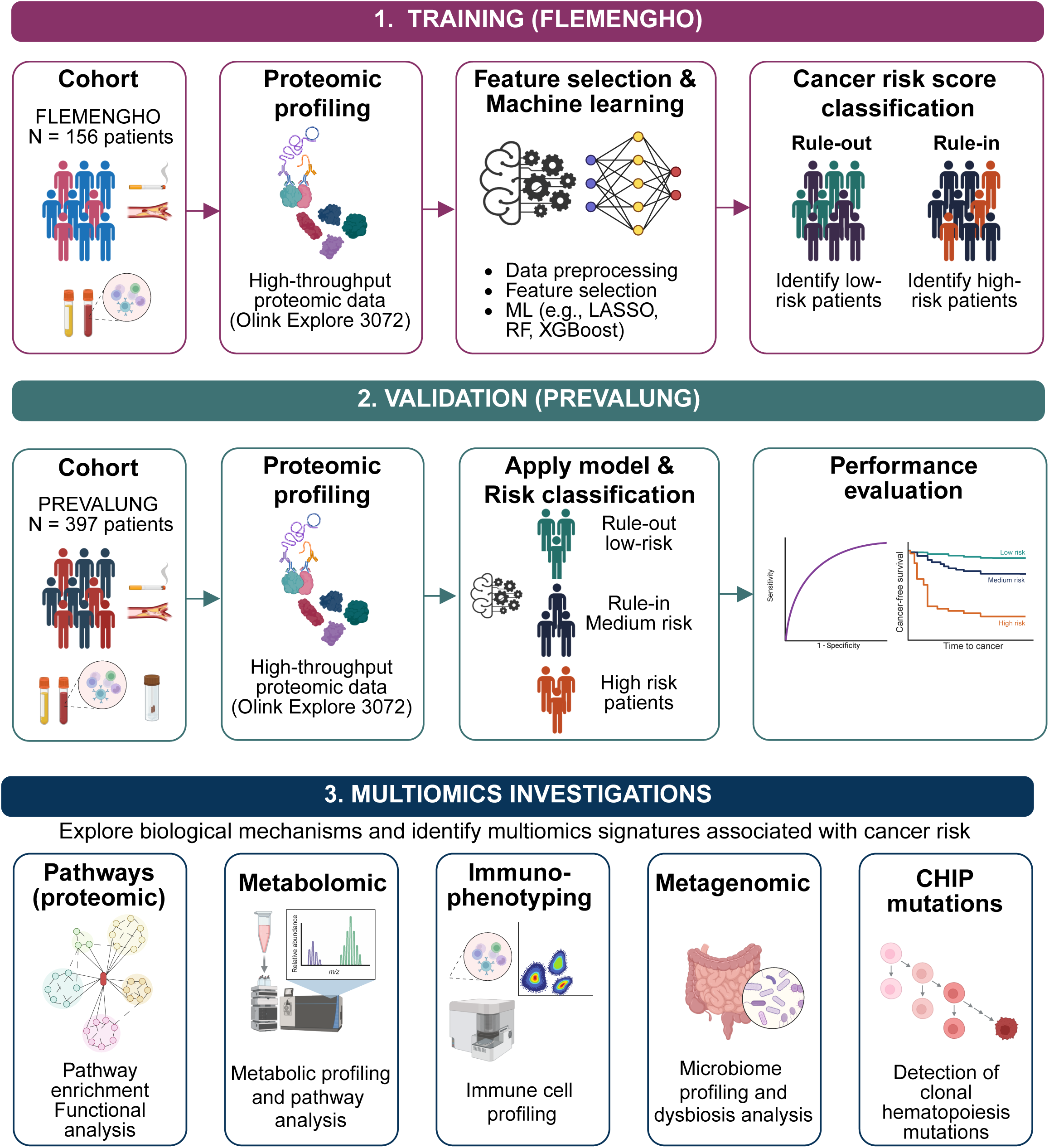
Cohort summary and machine learning model. **a-c**, Schematic overview of the study workflow. **1^st^ step**: Risk-score development in the FLEMENGHO training cohort (N = 156 participants with cardiovascular disease and tobacco exposure), high-throughput serum proteomic profiling was integrated with data pre-processing, feature selection, and machine-learning approaches, including logistic regression, LASSO, random forest, and XGBoost. This framework was used to develop a rule-out score for identification of individuals at low risk of cancer, a rule-in score for identification of individuals at higher risk, and a combined continuous risk score integrating the two stages (**a**). **2^nd^ step**: The resulting model was applied to the PREVALUNG transfer cohort (N = 397 participants), for whom proteomic risk scores were calculated. Model performance was evaluated using discrimination analyses (ROC curves and area under the curve [AUC]) and time-to-event analyses of cancer-free survival using Kaplan-Meier methods. Participants were subsequently stratified according to their predicted cancer risk (**b**). **3^rd^ step**: To investigate the biological mechanisms underlying the proteomic risk score and its association with cancer risk, complementary multiomics analyses were performed, including proteomic pathway analysis, plasma metabolomics, detection of clonal hematopoiesis of indeterminate potential (CHIP) mutations, immune-cell profiling by immunophenotyping, and stool metagenomic profiling of the gut microbiota. Integration of these datasets provided a multidimensional characterization of biological pathways and molecular signatures associated with cancer risk, supporting biomarker discovery and improved risk stratification and enabling investigation of mechanisms involved in cancer development and early detection (**c**). *Figure Created with BioRender.com*

After training-set median imputation and standardization, 2,910 plasma proteins were screened by complementary feature-selection procedures: mutual-information clustering, least absolute shrinkage and selection operator (LASSO) stability selection,^45^ and minimum-redundancy maximum-relevance (mRMR) ranking,^46^ followed by backward greedy ablation within inner cross-validation. Candidate learners included random forests and XGBoost,^47^ the locked pipeline ultimately used a soft-voting ensemble of Gaussian naive Bayes and class-balanced, L2-regularized logistic regression, grounded in classifier-combination theory.^48^ A four-analyte, bidirectional rule-out/rule-in architecture was used (**Fig. 2**). Both operating thresholds were determined in the FLEMENGHO discovery cohort. For Stage 1 (rule-out), the threshold maximizing specificity among operating points achieving sensitivity ≥0.925 was selected. Stage 2 (rule-in) was subsequently trained among participants not cleared by Stage 1, and its threshold was selected by maximizing specificity subject to sensitivity ≥0.40. Both thresholds were then locked and transferred unchanged to the external validation cohort. Imputation values, scaling parameters, classifiers and thresholds were then frozen and transferred to PREVALUNG without refitting. Feature ranking, signature selection and both thresholds were derived on FLEMENGHO. FLEMENGHO estimates are therefore apparent, not optimism-corrected. Signature selection additionally favoured configurations that transferred across cohorts, PREVALUNG among them; PREVALUNG is consequently a development cohort rather than an untouched test set. We state this rather than claim an independence the procedure does not support. Score distributions were visualized using kernel-density estimates and compared by non-parametric rank tests (**Fig. 2a-d**) and categorical assignments were evaluated using contingency matrices (**Fig. 2e-h**). “Rule-out” and “rule-in” name the two arms of the classifier and the strata they define. They do not name clinical decisions. No imaging decision follows from either label.

**Figure 2.**
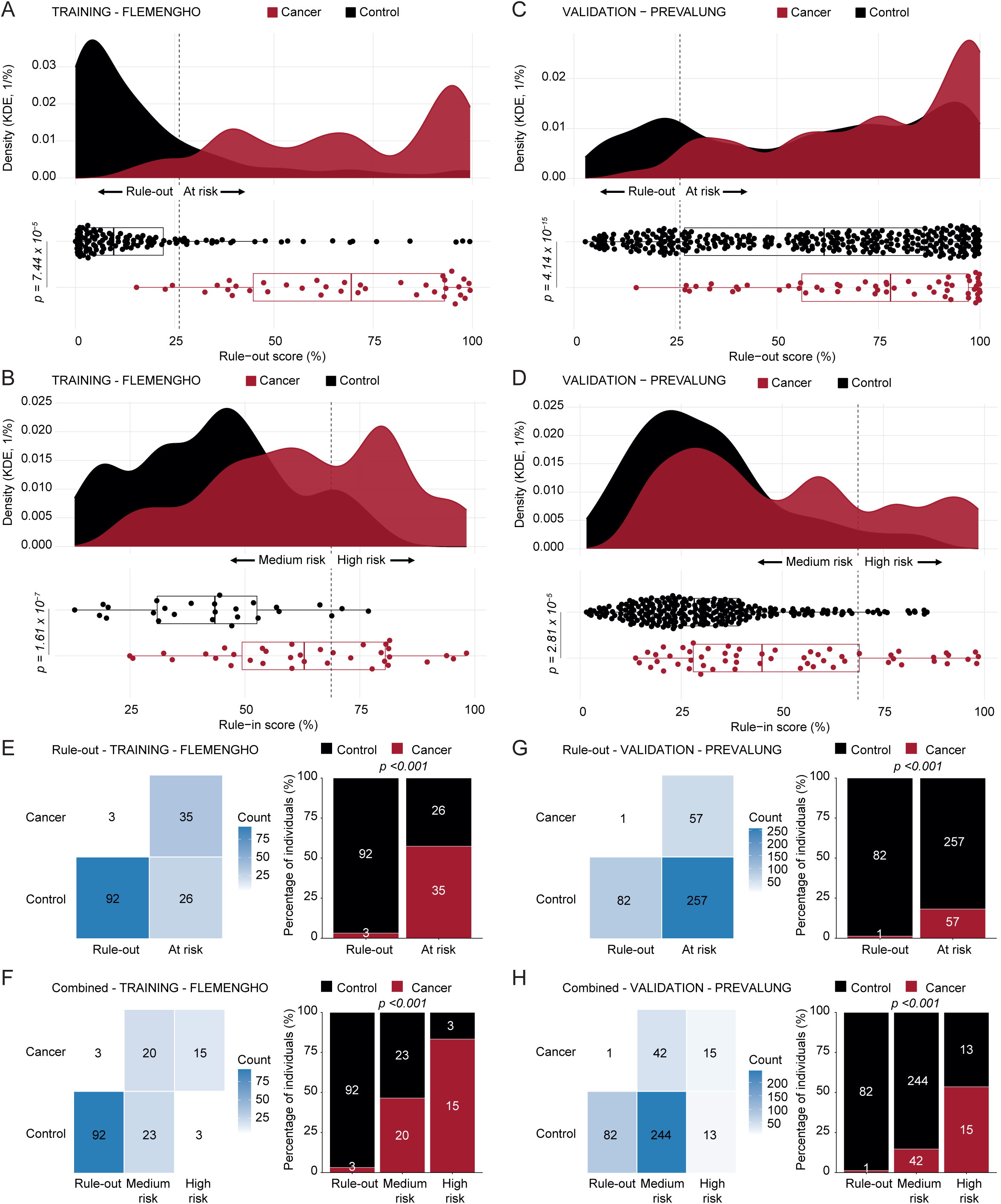
Distribution of the rule-out and rule-in scores in the discovery and transfer cohorts according to cancer status. **a-d**, The distribution of the rule-out score (**a and c**) and of the rule-in score (**b and d**) were compared between participants with incidental cancer (red) and control individuals (black) in the training (FLEMENGHO) cohort (N = 156) (**a-b**) and in the validation (PREVALUNG) cohort (N = 397) (**c-d**). The upper panel shows kernel density estimates (KDE) of the rule-out or rule-in score for each group. The lower panel displays individual participant values using a boxplot representing the median and interquartile range. Each dot represents one participant. The vertical dashed line indicates the predefined rule-out threshold (26.13%) and the rule-in threshold (68.73%). Differences between groups were assessed using a two-sided Wilcoxon rank-sum test. The corresponding *P* value is displayed on the figure. **e-h**, Confusion matrix (**left**) and contingency plots (**right**) of the rule-out score (**e and g**) and combined with rule-in score (**f and h**) classification in the training (FLEMENGHO) cohort (**e-f**) and in the validation (PREVALUNG) cohort (**g-h**). For confusion matrix (**left**), participants were classified according to their true clinical status (Control or Cancer) and their predicted rule-out category (Rule-out or At risk) or combined with the rule-in score (Rule-out, Medium risk or High risk). The heatmap displays the number of participants in each category combination. Cell color intensity is proportional to the number of individuals, and the exact counts are indicated within each cell (Count). For the contingency plots (**right**), the stacked bar chart shows the proportion of participants classified as Control (black) or Cancer (red) within category (Rule-out and At risk or Rule-out, Medium risk and High risk). Bars represent the percentage of individuals in each predicted category, and the absolute number of participants is displayed within each segment. The association between predicted rule-out category and clinical status was assessed using Pearson’s chi-squared test. The corresponding two-sided *P* value is shown in the figure.

Although the program measured extensive plasma proteomics and complementary metabolomic, clonal-hematopoiesis, immunophenotypic and gut-microbiome variables, the deployable classifier was reduced to four plasma proteins. Low plasma concentrations of O-GlcNAcase (OGA) and interleukin-6 (IL-6) together defined a stratum at very low risk of developing lung cancer in the first stage. Chymotrypsin C (CTRC) and β-microseminoprotein (MSMB) determined a second rule-in stage. The latter separated the non-ruled-out population into medium risk (high CTRC/low MSMB) and high risk (low CTRC/high MSMB). In FLEMENGHO, the OGA/IL-6 rule-out score differed strongly between future cases and controls (**Fig. 2a**). Low OGA and low IL-6 assigned 95 participants to the rule-out group, comprising 92 controls and only three future cancers. Thus, Stage 1 retained 35 of 38 cancers in the at-risk group (sensitivity, 92.1%) and achieved a negative predictive value (NPV) of 96.8% (**Fig. 2e**). Among the 61 participants retained; an outcome-dependent subsample carrying 35 of the 38 events, in which the rule-in arm was both built and evaluated; the CTRC/MSMB rule-in score also separated cases from controls (**Fig. 2b**): the medium-risk group contained 20 cancers and 23 controls, whereas the high-risk group contained 15 cancers and only three controls, yielding a high-risk positive predictive value (PPV) of 83.3% (**Fig. 2f**). The combined cancer frequencies were therefore 3.2%, 46.5% and 83.3% in the rule-out, medium-risk and high-risk groups, respectively (**Fig. 2f**). These frequencies are not population risks. The FLEMENGHO analytical sample was built by design: 38 cases, 52 healthy non-smoking controls, 64 ACVD controls. They therefore reflect the case–control sampling fraction. FLEMENGHO is a discovery set throughout; the clinical claims rest on PREVALUNG.

This ordered separation transferred to PREVALUNG. The rule-out score again separated cancers from controls (**Fig. 2c**). Only one of 58 future cancers, together with 82 controls, fell into the rule-out group, corresponding to 98.3% sensitivity for all cancers and an overall NPV of 98.8% (**Fig. 2g**). Among the 314 participants proceeding to Stage 2, 42 cancers and 244 controls were classified as medium risk, whereas 15 cancers and 13 controls were classified as high risk (score comparison in **Fig. 2d**; categorical association in **Fig. 2h**). The high-risk PPV was 53.6% and Stage 2 specificity was 94.9%. Accordingly, cancer prevalence increased from 1.2% in the rule-out group to 14.7% in the medium-risk group and 53.6% in the high-risk group; a 3.7-fold enrichment over cohort-wide prevalence and a 44-fold gradient between the extreme strata (**Fig. 2h**). Predictive values are conditional on the 14.6% cancer frequency of this cohort and will fall in a general screening population; they do not transport without recalibration. Exact binomial intervals for each operating characteristic, and performance restricted to the 26 incident lung cancers, are given in **Supplementary Table S3**.

In conclusion, a two-stage classifier ruling out low-risk patients (with low OGA and IL-6) and then ruling in high-risk patient (with low CTRC and high MSMB) provides a method to predict the risk of future cancer development among high-risk patients with CVD exposure.

### Biological correlates of the low-risk classifier: low OGA and IL-6

Circulating OGA was modestly higher in individuals who subsequently developed cancer than in cancer-free controls (*P = 0.02*; **Fig. 3a**). To determine whether plasma OGA reflected its abundance in circulating immune cells, OGA was quantified by immunoblot technology (**Extended data Fig. 1**) or flow cytometry (**Fig. 3b-c**) in PBMCs. Plasma OGA correlated positively with cellular OGA across total PBMCs and all major lymphoid compartments examined, with the strongest associations in CD3^+^ and CD4^+^ T cells and no significant association in monocytes (**Fig. 3b**). Consistently, participants with plasma OGA above the median had higher intracellular OGA in total PBMCs (*P = 0.0002*), CD19^+^ B cells (*P = 0.024*), CD3^+^ T cells (*P = 0.00079*), CD4^+^ T cells (*P = 0.0015*), CD8^+^ T cells (*P = 0.0009*) and CD56^+^ natural killer (NK) cells (*P = 0.0075*; **Fig. 3c**). Circulating OGA was therefore associated with a coordinated immune-cell phenotype rather than behaving as an isolated plasma signal. Plasma OGA was embedded in a broader glycometabolic network.^49^ Its strongest positive protein correlates included GPI, GNPDA1, AMDHD2, GNE, NAGK and UXS1, together with numerous glycosyltransferases and glycan-modifying enzymes, including FUT3/FUT5, FUT8, B4GALT1, GCNT1, several GALNT family members, ST3GAL1, ST6GAL1, B3GNT7, B3GAT3 and EXTL1 (**Fig. 3d**). Metabolite correlations were weaker but converged on related chemistry,^50,51^ encompassing glucosamine and sugar phosphates as well as N-acetylated amino acids and multiple acetylated polyamine derivatives, including N-acetylputrescine, N-acetylspermine and N-acetylspermidines (**Fig. 3e**). Collectively, these associations link OGA to systemic hexosamine utilization, glycan remodeling and acetylated amine metabolism.

**Figure 3.**
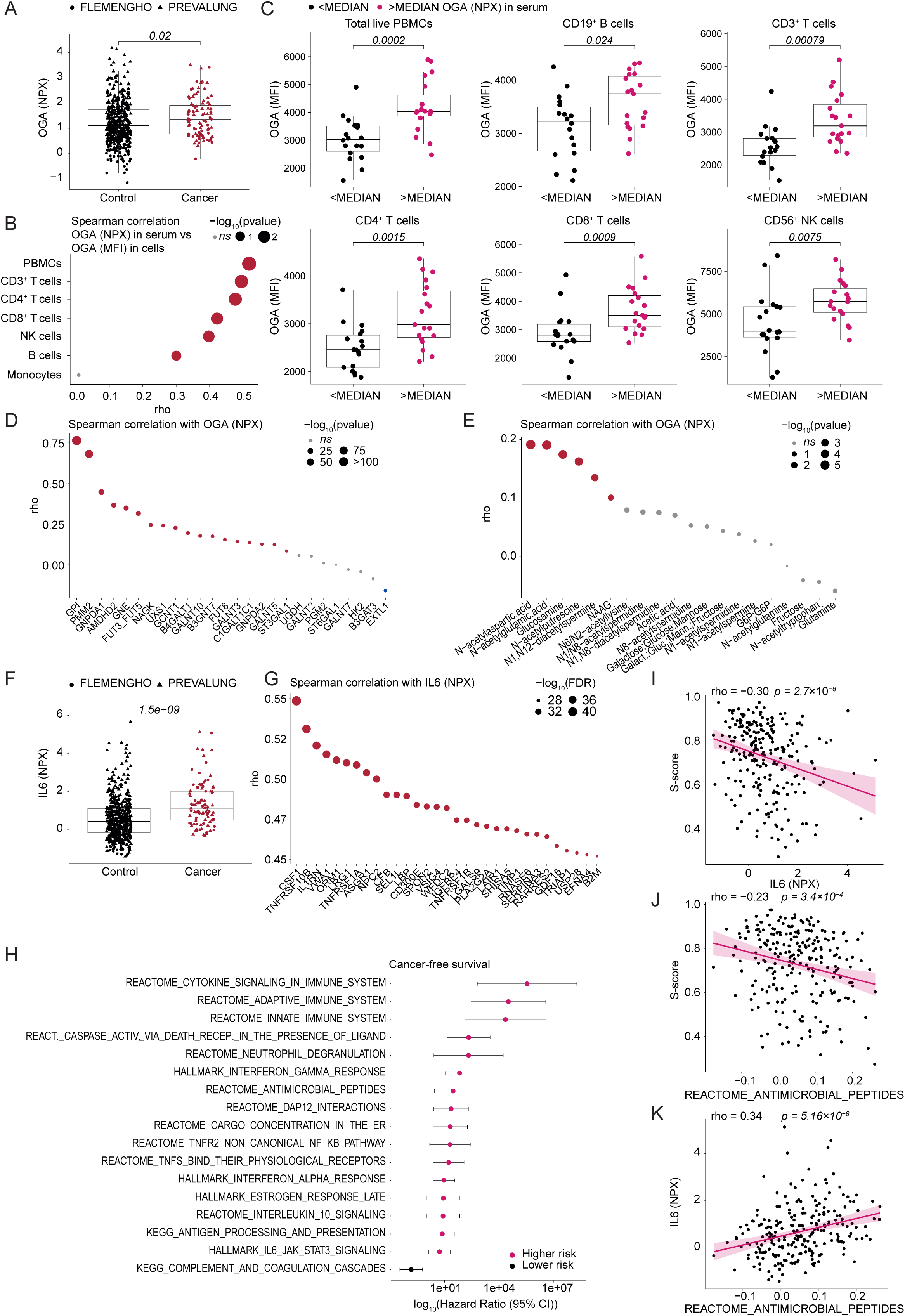
Multiomic biological insights into rule-out score biomarkers. **a**, OGA protein levels measured by Olink proteomics (NPX) in the plasma. Participants were stratified according to their clinical status (Control or Cancer). Boxplots display the median and interquartile range, while each dot represents individual participants. Each individual from the FLEMENGHO (N = 154) and PREVALUNG (N = 396) cohorts is represented by solid circles and by triangles, respectively. Difference between groups is assessed using two-sided Wilcoxon rank-sum tests, and the corresponding *P* value is shown on the figure. **b-c**, Association between plasma OGA protein levels and immune-cell OGA expression measured by flow cytometry (N = 37 patients). **b**, Spearman correlation analyses were performed between plasma OGA protein abundance measured by Olink proteomics (NPX) and OGA median fluorescence intensity (MFI) measured by flow cytometry in the indicated immune cell populations from peripheral blood mononuclear cells (PBMCs). Each dot represents one immune-cell population. The x-axis indicates the Spearman correlation coefficient (rho), while dot size is proportional to the statistical significance (−log_10_(*P* value)). Dot color represents the direction and magnitude of the correlation, with blue and red indicating negative and positive correlations, respectively. Non-significant (*ns*) correlations are indicated in grey. **c**, Participants were stratified according to plasma OGA protein abundance into two groups using the cohort median (<MEDIAN and >MEDIAN). OGA expression measured by flow cytometry (MFI) was compared between groups for each immune-cell population. Boxplots display the median and interquartile range, while each dot represents individual participants. Differences between groups were assessed using two-sided Wilcoxon rank-sum tests, and the corresponding *P* values are shown on each panel. **d-e**, Correlations between plasma OGA protein abundance and proteins (**d**) or metabolites (**e**) involved in the hexosamine biosynthetic pathway (HBP) and related glycosylation pathways. Spearman rank correlation analyses were performed between plasma OGA protein abundance and a curated panel of proteins involved in the hexosamine biosynthetic pathway (HBP), O-GlcNAc cycling, monosaccharide metabolism, nucleotide-sugar metabolism, and downstream glycosylation processes (**d**) or metabolites involved in, or functionally associated with, the HBP, including carbohydrates, amino acids, acetylated metabolites, and polyamine derivatives (**e**), in individuals from FLEMENGHO (N = 154) and PREVALUNG (N = 396) cohorts. Each dot represents one protein (**d**) or one metabolite (**e**). The x-axis indicates the Spearman correlation coefficient (rho), while dot size is proportional to the statistical significance (−log_10_(*P* value)). Proteins or metabolites significantly correlated with OGA (*P value ≤ 0.05*) are color-coded according to the direction of the correlation (blue, negative; red, positive), whereas non-significant (*ns*) correlations are displayed in grey. **f**, same as in a for IL6 protein levels measured by Olink proteomics (NPX) in the plasma. **g**, Spearman correlation analyses were performed between plasma IL6 protein abundance and proteins consistently associated with IL6 across the FLEMENGHO and PREVALUNG cohorts. The protein panel corresponds to the intersection of proteins significantly correlated with IL6 after Benjamini–Hochberg false discovery rate (FDR) correction and with absolute rho value >0.3 in both cohorts (n = 187 proteins; refer to Extended data Fig. 2a). Each dot represents one protein. The x-axis indicates the Spearman correlation coefficient (rho), while dot size is proportional to the statistical significance (−log_10_(FDR)). Significant positive correlations are in red. **h**, Association between IL6-associated pathway activity scores and cancer-free survival in the combined FLEMENGHO (N = 154) and PREVALUNG (N = 396) cohorts. Pathway activity scores were calculated using single-sample Gene Set Enrichment Analysis (ssGSEA) from Hallmark, KEGG, and Reactome pathways identified from the common set of 187 proteins significantly associated with plasma IL6 (refer to Extended data Fig. 2a). Associations between continuous pathway activity scores and cancer-free survival were evaluated in the pooled FLEMENGHO and PREVALUNG cohorts using univariable Cox proportional hazards regression. Forest plots display hazard ratios (log_10_(HR)) and 95% confidence intervals (95% CI) for a one-unit increase in pathway activity score. Only pathways with a *P* value < 0.05 are shown. Pathways associated with an increased cancer risk (HR > 1) are shown in pink, whereas pathways associated with a reduced cancer risk (HR < 1) are shown in black. The vertical dashed line indicates HR = 1. **i-k**, Spearman correlation analyses were performed in the PREVALUNG cohort to assess the associations between plasma IL6 protein abundance measured by Olink proteomics (NPX), the microbiome-derived S-score of dysbiosis, and the REACTOME_ANTIMICROBIAL_PEPTIDES pathway activity score from ssGSEA. Each dot represents one participant. The solid line indicates the linear regression fit with the corresponding 95% confidence interval (shaded area). Spearman correlation coefficients (rho) and two-sided nominal *P* values are displayed for each comparison.

IL-6 defined a complementary inflammatory axis. IL-6 was increased before diagnosis in participants who developed cancer (*P = 1.5 × 10^−9^*; **Fig. 3f**) and correlated reproducibly with 187 plasma proteins in both cohorts (**Fig. 3g** and **Extended data Fig. 2a**). Prominent correlates included CSF1, TNFRSF10B, IL1RN, LRG1, TNFRSF1A, VSIG4, LBP, WFDC2, TNFRSF1B, LGALS9, TIMP1, RNASE6, SERPINA3, GDF15, LAIR1 and B2M. Pathway analysis connected this shared protein set to cytokine signaling, innate and adaptive immunity, neutrophil degranulation, interferon responses, TNF-family signaling, complement and antimicrobial peptides (**Extended data Fig. 2b**). Many of these IL-6-associated programs were themselves associated with shorter cancer-free survival, particularly cytokine, innate and adaptive immune programs and caspase activation through death receptors (**Fig. 3h**). This inflammatory state also intersected with the gut microbiota. IL-6 correlated inversely with the continuous S-score derived from the balance between the previously defined immunosuppressive SIG1 and immunostimulatory SIG2 microbial communities (rho = −0.30, *P = 2.7 × 10^−6^*; **Fig. 3i**).^52^ The plasma antimicrobial-peptide score similarly correlated inversely with the S-score (rho = −0.23, *P = 3.4 × 10^−4^*; **Fig. 3j**) and positively with IL-6 (rho = 0.34, *P = 5.16 × 10^−8^*; **Fig. 3k**). Accordingly, IL-6 and the antimicrobial-peptide program were highest in SIG1-classified participants and lowest in the SIG2 group, with intermediate values in the intermediate (grey) category (**Extended data Fig. 2c–f**). These data connect low IL-6 to a distinct microbiota-associated, low-inflammatory systemic state.

The combined low-OGA/low-IL-6 phenotype extended across the plasma proteome. Principal-component analysis separated ruled-out from at-risk participants in both FLEMENGHO (PERMANOVA R² = 0.036, *P = 0.001*) and PREVALUNG (R² = 0.028, *P = 0.001*; **Fig. 4a**). Of the proteins differentiating these categories, 523 were shared across cohorts, compared with 249 and 447 cohort-specific proteins, respectively (**Fig. 4b**). Concordant pathway shifts involved adaptive immunity, Toll-like-receptor and B-cell-receptor signaling, TNF and interferon-γ signaling, RIPK1-mediated regulated necrosis and death-receptor-dependent caspase activation; these programs were increased in future cancer cases and negatively associated with cancer-free survival (**Fig. 4c, d and Extended data Fig. 3**). In FLEMENGHO and PREVALUNG, B-cell-receptor and TNF-signaling scores rose nearer to diagnosis (rho = −0.36 and −0.33, respectively; **Fig. 4e, f**), compatible with progressive immune activation during the prediagnostic interval and a pioneering report.^53^ The rule-out score was also correlated with a proteomic aggregate score reflecting a regulatory T-cell (Treg) signature, derived from 128 proteins shared with single-cell sequencing data from human cancer liver tissue,^54^ including Treg-associated proteins such as IL2RA (CD25), CTLA4, TIGIT, TNFRSF4 (OX40), IKZF2 (HELIOS), LAYN, and TNFRSF9 (4-1BB) (rho = 0.43, *P < 0.0001*), which was higher in future cancer cases (*P = 8.1 × 10^−8^*; **Fig. 4g, h**). Immunophenotyping confirmed increased CD25^hi^CD127^lo^ Treg cells among CD4^+^ cells and increased CCR4^+^ regulatory T cells among PBMCs in cases (*P = 0.014* and *P = 0.041*, respectively; **Fig. 4i, j and Supplementary Fig. 3 for gating strategy**). Ruled-out participants also had uniformly lower concentrations of six lung-cancer-relevant, potentially actionable proteins: GDF15, IL4R, TREM2, MAP2K1, ACBP/DBI and BCL2L1 (**Extended data Fig. 4a**). Jointly, this cluster predicted cancer-free survival (**Extended data Fig. 4b-c**, hazard ratio, 2.45; 95% confidence interval, 1.62–3.71; *P < 0.0001*), and five of the six proteins were individually prognostic (**Extended data Fig. 4d-i**).

**Figure 4.**
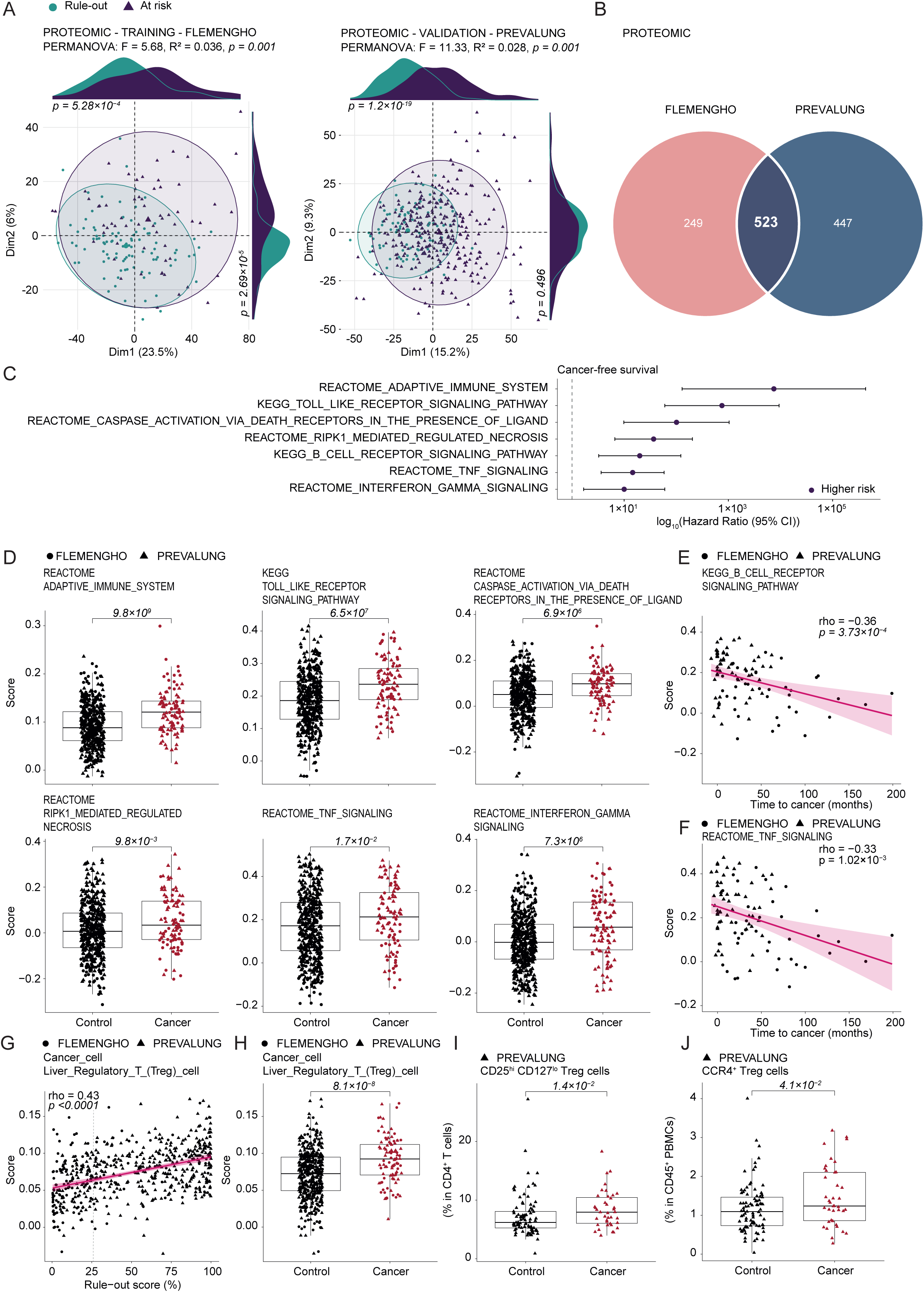
Unsupervised identification of biological and immune pathways associated with cancer risk stratification by the rule-out score. **a-b**, Unsupervised proteomic profiling according to rule-out score-defined cancer risk groups in the FLEMENGHO and PREVALUNG cohorts. **a**, Principal component analyses (PCA) were performed using the Olink Explore 3072 proteomic panel to assess global plasma proteomic patterns associated with rule-out score-defined cancer risk groups in FLEMENGHO (**left**, N = 155) and in PREVALUNG (**right**, N = 397). PCA plots display individual samples projected onto the first two principal components and colored according to the rule-out score categories (Rule-out and At risk). Ellipses represent the 90% confidence interval for each group. Differences in global proteomic profiles between groups were assessed using permutational multivariate analysis of variance (PERMANOVA) based on Euclidean distances. The proportion of variance explained (R²), F statistic, and nominal *P* value are indicated. Marginal density plots along the x- and y-axes represent the distribution of samples along principal components 1 and 2, respectively, for each risk group. Pairwise differences in principal component coordinates were assessed using Welch’s t-test, with corresponding *P* values shown for dimensions (Dim) 1 and 2. **b**, Venn diagram showing the overlap between PCA-contributive proteins identified independently in the FLEMENGHO and PREVALUNG cohorts. Proteins were selected based on their contribution to significantly discriminating principal components between rule-out score categories. In each cohort, only principal components with significant differences between groups according to Welch’s t-test were considered (Dim 1 and 2 in FLEMENGHO; Dim 1 in PREVALUNG). Proteins contributing above the theoretical PCA contribution threshold and significantly associated with rule-out score categories after Benjamini–Hochberg FDR correction (FDR < 0.05) were retained. The intersection represents proteins consistently associated with rule-out score stratification across both independent cohorts (n = 523 proteins; refer to **Supplementary Table S5**). **c**, Pathways significantly associated with cancer-free survival in both FLEMENGHO and PREVALUNG (*P* < 0.05 in each cohort) were retained for pooled univariable Cox proportional hazards regression. Forest plots show HRs and 95% CIs for continuous pathway activity scores; the dashed line indicates HR = 1. **d**, Participants were stratified according to their clinical status (Control or Cancer) and pathway activity scores significantly associated with cancer-free survival were compared between groups. Boxplots display the median and interquartile range, while each dot represents individual participants. Each individual from the FLEMENGHO (N = 154) and PREVALUNG (N = 396) cohorts is represented by solid circles and by triangles, respectively. Difference between groups is assessed using two-sided Wilcoxon rank-sum tests, and the corresponding *P* value is shown on the figure. **e-f**, Spearman correlation analyses were performed to assess the associations between time to cancer diagnosis in months and pathway activity scores. Each dot represents one participant. The solid line indicates the linear regression fit with the corresponding 95% confidence interval (shaded area). Each individual from the FLEMENGHO (N = 154) and PREVALUNG (N = 396) cohorts is represented by solid circles and by triangles, respectively. Spearman correlation coefficients (rho) and two-sided nominal *P* values are displayed for each comparison. **g**, Spearman correlation analysis was performed to assess the associations between the cell-type signature score and the continuous rule-out score. The Regulatory T cell (*Liver, cancer tissue*) signature was significantly associated with the rule-out score. Each dot represents one participant. The solid line indicates the linear regression fit with the corresponding 95% confidence interval (shaded area). Each individual from the FLEMENGHO (N = 154) and PREVALUNG (N = 396) cohorts is represented by solid circles and by triangles, respectively. The dashed line indicates the rule-out score threshold of 26.13%. Spearman correlation coefficients (rho) and two-sided nominal *P* values are displayed for each comparison. **h**, Participants were stratified according to their clinical status (Control or Cancer) and the Regulatory T cell (*Liver, cancer tissue*) pathway score was compared between groups. Boxplots display the median and interquartile range, while each dot represents individual participants. Each individual from the FLEMENGHO (N = 154) and PREVALUNG (N = 396) cohorts is represented by solid circles and by triangles, respectively. Difference between groups is assessed using two-sided Wilcoxon rank-sum tests, and the corresponding *P* value is shown on the figure. **i-j**, Participants were stratified according to their clinical status (Control or Cancer), and the percentage of regulatory T cells (Tregs) among circulating CD4^+^ T cells (**i**) and of CCR4^+^ Tregs among total PBMCs (**j**) was compared between groups in the PREVALUNG cohort, for which spectral flow cytometry data were available (refer to **Supplementary Figure 3** for gating strategy). Boxplots display the median and interquartile range, and each dot represents an individual participant. Each participant is represented by a triangle (N = 129). Difference between groups is assessed using two-sided Wilcoxon rank-sum tests, and the corresponding *P* value is shown on the figure.

Finally, comparison of our dataset with several external biomarker frameworks^8,9,16,17^ showed that several established inflammatory and lung-cancer-risk proteins mapped predominantly to the at-risk proteomic field (**Extended data Fig. 5a-d**). In our dataset, IL-6 recurred in four independent measurements, whereas OGA provided a distinct, nonredundant glycometabolic signal (**Extended data Fig. 5a**). IL-6 has been confirmed in two other datasets (**Extended data Fig. 5b, d**), ^8,9,16,17^ while OGA was unique to our rule-out model. Together, low OGA and low IL-6 identify a biologically coherent state characterized by attenuated glycometabolic remodeling, systemic inflammation, antimicrobial defense and regulatory immune activation.

### Biological correlates of the high-risk classifier: low CTRC and high MSMB

To define the biological context of the two proteins retained for the rule-in step, we first examined their tissue distribution, comparing them to the close-to-ubiquitous proteins OGA and IL-6. Analysis of GTEx transcript abundance and Human Protein Atlas immunohistochemistry showed sharply contrasting patterns (**Extended data Fig. 6a, b**). CTRC expression was essentially confined to the pancreas, consistent with its production by exocrine acinar cells.^55^ By contrast, MSMB was detectable across multiple non-prostatic tissues, including arteries, heart, lung and skin, indicating that the circulating protein cannot be assumed to originate exclusively from the male genitourinary tract despite its conventional designation.

We next adapted a plasma-proteomic deconvolution strategy^56^, in which proteins significantly correlated or anticorrelated with the circulating factor of interest are assigned to tissues according to their expression profiles. CTRC-associated proteins were overwhelmingly enriched in the pancreas **(Extended data Fig. 6c)**, and this specificity became even more pronounced when the analysis was restricted to correlates shared by both cohorts (**Extended data Fig. 6d**). In contrast, MSMB showed a distributed tissue-correlation pattern, with shared positive correlates assigned to several tissues, most prominently lung, skin, arteries and brain, as well as blood and spleen (**Extended data Fig. 6e, f**). Cell-type-specific proteomic annotations independently supported these assignments: the CTRC network showed expression patterns compatible with a pancreatic acinar origin, whereas the MSMB network was associated with signatures of diverse immune and stromal populations, including monocytes, microglia, regulatory T cells, plasmacytoid and pulmonary dendritic cells, vascular endothelial cells and bone-marrow myeloid cells (**Extended data Fig. 6g, h**). Of note, plasma MSMB did not differ between men and women in either FLEMENGHO and PREVALUNG, again arguing against a dominant sex-specific source (**Extended data Fig. 6i**).

Circulating CTRC was significantly reduced in participants who subsequently developed cancer across the two cohorts (*P = 1.2 × 10^−6^*; **Fig. 5a**). Cohort-wise correlation analysis identified 263 CTRC-associated proteins unique to FLEMENGHO, one unique to PREVALUNG and 20 shared between them (**Fig. 5b**). These reproducible correlates were dominated by pancreatic digestive proteins, including PNLIP, CELA3A, SERPINI2, CTRL, CTRB1, CPB1, CPA1, PNLIPRP1, AMY2A/AMY2B, PRSS2 and CLPS (**Fig. 5c**). Accordingly, pathway enrichment converged on pancreatic acinar-cell development, exocrine-pancreas lineage specification, digestion and absorption, starch and sucrose metabolism, glycerolipid metabolism, and vitamin or fat-soluble-vitamin metabolism (**Fig. 5d**). Higher scores for these CTRC-linked pathways were consistently associated with a lower hazard of future cancer, with all selected signatures displaying hazard ratios below one for cancer-free survival (**Fig. 5e**). Thus, low CTRC marked the attenuation of a highly coherent circulating exocrine-pancreatic program rather than an isolated change in one protease.

**Figure 5.**
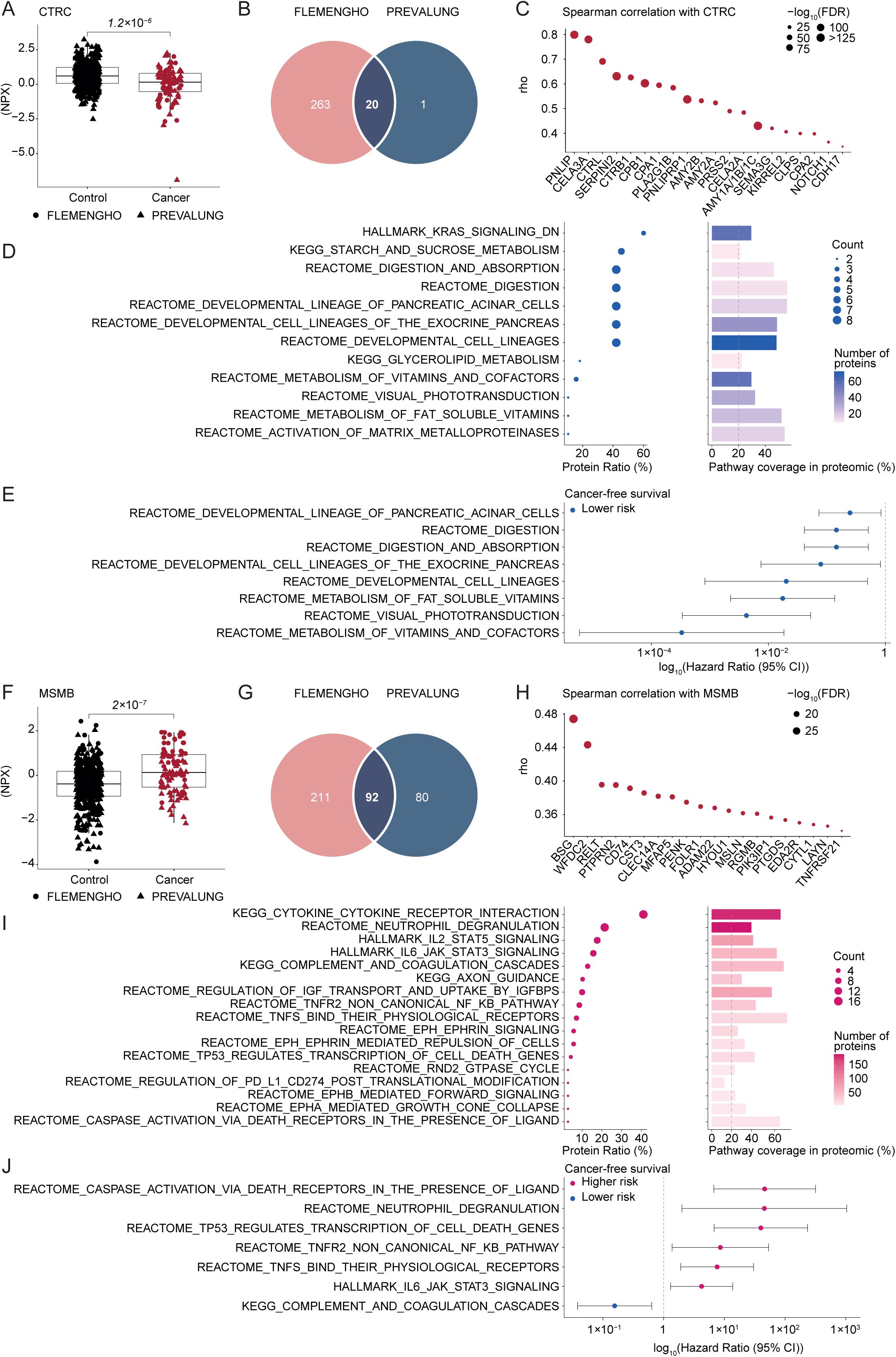
Biological correlates of the high-risk classifier from the rule-in score. **a**, CTRC protein levels measured by Olink proteomics (NPX) in the plasma. Participants were stratified according to their clinical status (Control or Cancer). Boxplots display the median and interquartile range, while each dot represents individual participants. Each individual from the FLEMENGHO (N = 154) and PREVALUNG (N = 396) cohorts is represented by solid circles and by triangles, respectively. Difference between groups is assessed using two-sided Wilcoxon rank-sum tests, and the corresponding *P* value is shown on the figure. **b-c**, Venn diagram showing the overlap between CTRC-correlated proteins identified independently in the FLEMENGHO and PREVALUNG cohorts. Proteins were selected based on significant correlations with plasma CTRC in each cohort (Benjamini–Hochberg FDR *< 0.05* and absolute Spearman rho > 0.3) and retained only when consistently identified in both cohorts (**b**). Spearman rank correlation analyses were performed between plasma CTRC levels and the 20 common proteins identified in FLEMENGHO (N = 154) and PREVALUNG (N = 396) participants (**c**). Each dot represents one protein. The x-axis indicates the Spearman correlation coefficient (rho), and dot size is proportional to statistical significance (−log_10_(FDR)). Proteins significantly correlated with CTRC (Benjamini–Hochberg FDR *< 0.05*) are color-coded according to the direction of the association (blue, negative correlation; red, positive correlation), whereas non-significant (ns) correlations are shown in grey. **d**, Pathway enrichment and coverage analysis were performed on the 20 proteins consistently correlated with plasma CTRC in the FLEMENGHO and PREVALUNG cohorts. Hallmark, KEGG Legacy, and Reactome pathways enriched by over-representation analysis (ORA) are shown. The left panel displays the proportion of CTRC-associated proteins assigned to each pathway (Protein Ratio (%)), with dot size indicating the number of contributing proteins (Count). The right panel shows pathway coverage within the Olink Explore 3072 proteomic platform, calculated as the percentage of proteins defining each pathway in MSigDB that were quantified by Olink. Bar color indicates the number of detected pathway proteins. The dashed line indicates the 20% pathway coverage threshold used for pathway selection in downstream analyses. Only pathways with *P* value < 0.05 are displayed. **e**, Pathway activity scores derived from ssGSEA were calculated for pathways identified in **d** (at least 10 proteins, coverage >20%, and *P* value < 0.05). Associations between continuous pathway activity scores and cancer-free survival were evaluated in the pooled FLEMENGHO (N = 154) and PREVALUNG (N = 396) cohorts using univariable Cox proportional hazards regression models. Forest plots display hazard ratios (HRs) and 95% confidence intervals (95% CIs) for a one-unit increase in pathway activity score. The x-axis is displayed on a logarithmic scale, and the vertical dashed line indicates HR = 1. Pathways associated with significant decreased cancer risk (HR < 1 and FDR < 0.05) are shown in blue. **f**, Same as in a for MSMB levels. **g-h**, Same as in b-c MSMB-correlated proteins identified independently in the FLEMENGHO and PREVALUNG cohorts. Spearman rank correlation analyses were performed between plasma MSMB levels and the 92 common proteins identified in FLEMENGHO (N = 154) and PREVALUNG (N = 396) participants. The 20 most strongly correlated shared proteins are shown. (**h**). **i**, Same as in d for the 92 proteins identified in g. **j**, Same as in e. Forest plots display hazard ratios (HRs) and 95% confidence intervals (95% CIs) for a one-unit increase in pathway activity score. The x-axis is displayed on a logarithmic scale, and the vertical dashed line indicates HR = 1. Pathways associated with significant increased (HR > 1 and *P* value < 0.05) or decreased cancer risk (HR < 1 and FDR < 0.05) are shown in pink or in blue, respectively.

MSMB exhibited the opposite direction: its plasma concentration was significantly increased in future cancer cases (*P = 2 × 10^−7^*; **Fig. 5f**), consistent with high MSMB contributing to the high-risk branch. MSMB also belonged to a broader and more reproducible network than CTRC, with 211 correlates unique to FLEMENGHO, 80 unique to PREVALUNG and 92 shared between cohorts (**Fig. 5g**). The 20 strongest common correlates included BSG, WFDC2, CD74, CLEC14A, MFAP5, PENK, FOLR1, ADAM22, MSLN, among others (**Fig. 5h**). Pathway analysis linked this network to cytokine–receptor interactions, neutrophil degranulation, IL-2–STAT5 and IL-6–JAK–STAT3 signaling, complement and coagulation, TNF–TNFR2/NF-κB signaling, ephrin pathways, TP53-regulated cell-death genes and death-receptor-mediated caspase activation (**Fig. 5i**). Most selected inflammatory and cell-death signatures predicted increased cancer hazard, whereas the complement/coagulation signature was associated with lower risk (**Fig. 5j**).

Finally, comparison with four literature-derived rule-in protein sets ^8,9,16,17^ revealed no protein-level overlap with our rule-in algorithm (**Extended data Fig. 7a–d**). The CTRC–MSMB rule-in axis therefore captured a distinct biological configuration combining loss of an exocrine-pancreatic program with elevation of a multisystem inflammatory, stromal and cell-death-associated network.

### Potential clinical utility of the two-step classifier

Beyond its biological implications, the sequential four-protein classifier stratified risk to a degree that could be clinically meaningful. In the FLEMENGHO discovery set, the OGA–IL-6 rule-out phase achieved an AUC of 0.924 (95% CI, 0.883–0.965), whereas the CTRC–MSMB rule-in phase, applied to individuals not eliminated during the first step, yielded an AUC of 0.789 (95% CI, 0.677–0.901). Integration of both phases into the complete classifier preserved high apparent discrimination, with an AUC of 0.909 (95% CI, 0.852–0.966; **Fig. 6a–c**). Thus, the classifier combined a highly effective first step that excluded individuals unlikely to develop lung cancer with a complementary second step that identified a smaller subgroup at particularly elevated risk. This discrimination translated into striking differences in cancer-free survival. In FLEMENGHO, participants retained after the rule-out step exhibited a 15.48-fold higher hazard of lung cancer than ruled-out individuals (95% CI, 4.75–50.46; *P<0.0001*; **Fig. 6d**). Subsequent rule-in classification generated three separated trajectories: compared with the ruled-out group, the hazard ratio was 12.95 (95% CI, 3.83–43.82) for the medium-risk group and 23.39 (95% CI, 6.76–80.92) for the high-risk group. Even within the population retained after rule-out, high-risk individuals experienced approximately twice the hazard of those at medium risk (HR, 2.04; 95% CI, 1.03–4.03; **Fig. 6e**).

**Figure 6.**
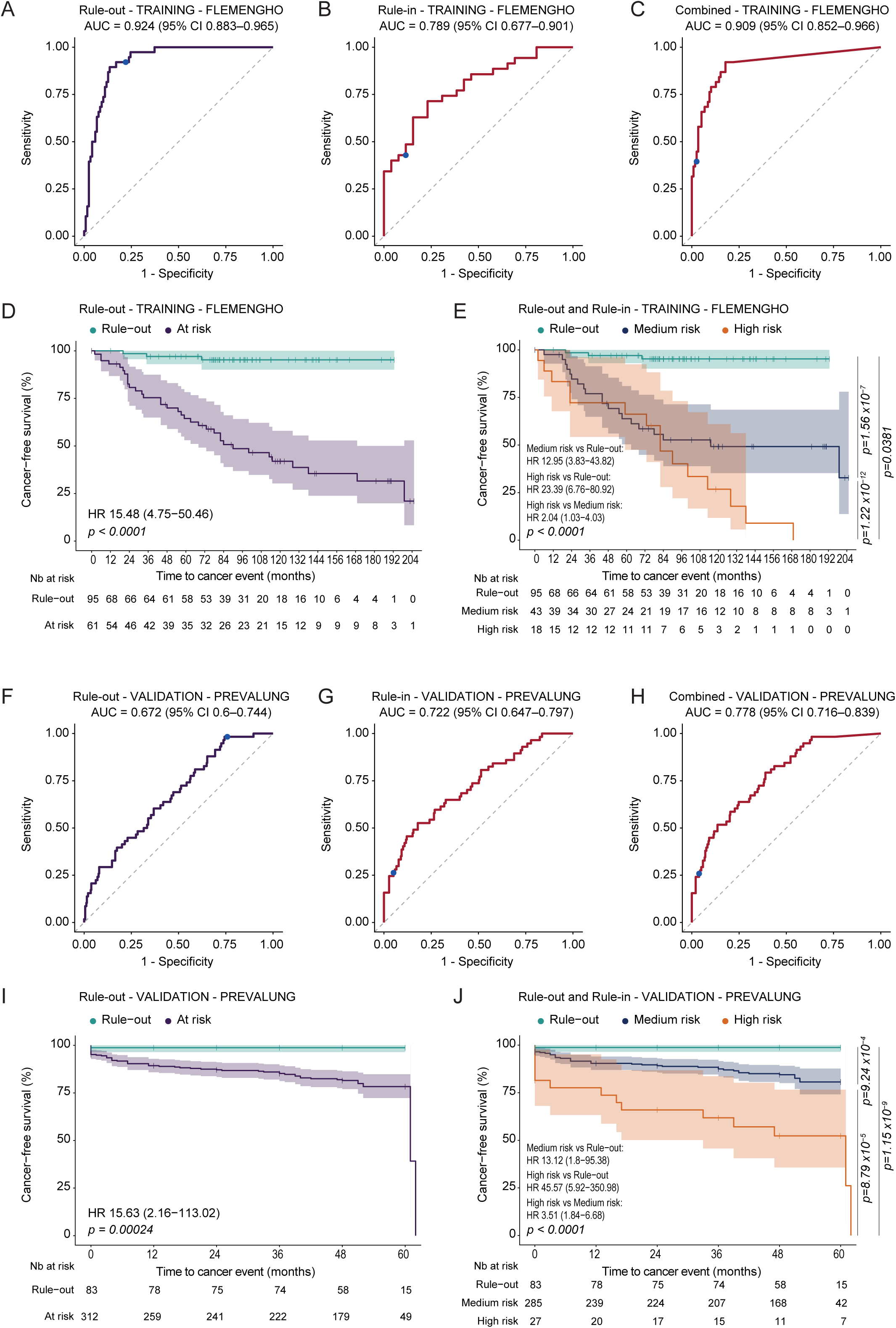
Performance evaluation and prognostic stratification of the two-stage cancer risk score. **a-c**, Receiver operating characteristic (ROC) curve analyses evaluating the discrimination of cancer status in the FLEMENGHO training cohort (N = 156). Performance of the rule-out score using the predefined threshold of 26.13% (**a**). Performance of the rule-in score among individuals classified as at risk after the rule-out step (rule-out score ≥26.13%) using the predefined threshold of 68.73% (**b**). Combined two-stage model integrating the rule-out classification and the continuous rule-in score using the predefined threshold of 68.73% (**c**). ROC curves display sensitivity versus 1-specificity, with the corresponding area under the curve (AUC) and 95% confidence interval (CI) indicated. The blue dot represents the predefined score threshold. **d-e**, Kaplan-Meier analyses of cancer-free survival according to risk-score categories in the FLEMENGHO training cohort (N = 156). Cancer-free survival according to rule-out score categories (Rule-out versus At risk). (**d**). Cancer-free survival according to the combined two-stage risk classification: Rule-out (rule-out score <26.13%), Medium risk (rule-out score ≥26.13% and rule-in score <68.73%), and High risk (rule-out score ≥26.13% and rule-in score ≥68.73%) (**e**). Curves represent cancer-free survival probabilities over follow-up time, with the number of participants at risk displayed below each plot. Global and pairwise log-rank test *P* values are shown. Hazard ratios (HRs) and 95% CIs were estimated using univariable Cox proportional hazards regression models. **f-j**, Validation analyses performed in the PREVALUNG cohort (N = 397) following the same methodology as in a-e. ROC curve analyses evaluating the rule-out score, rule-in score, and combined two-stage model, respectively (**f-h**). Kaplan-Meier analyses of cancer-free survival according to rule-out categories and combined two-stage risk classification, respectively (**i-j**).

The classifier retained performance on transfer to PREVALUNG. The rule-out, rule-in and combined AUCs were 0.672 (95% CI, 0.600–0.744), 0.722 (95% CI, 0.647–0.797) and 0.778 (95% CI, 0.716–0.839), respectively (**Fig. 6f–h**). The survival analyses were particularly compelling: participants not ruled out had a 15.63-fold higher cancer hazard than the ruled-out group (95% CI, 2.16–113.02; *P = 0.00024*; **Fig. 6i**). In the complete three-tier classification, medium- and high-risk individuals had hazard ratios of 13.12 (95% CI, 1.80–95.38) and 45.57 (95% CI, 5.92–350.98), respectively, relative to the ruled-out population. High-risk participants also had a 3.51-fold greater hazard than medium-risk participants (95% CI, 1.84–6.68; overall *P<0.0001*; **Fig. 6j**). Performance depended on the interval between sampling and diagnosis, and we report it by horizon rather than as a single figure. Cancers diagnosed within 12 months of sampling were separated from controls only weakly (AUC 0.589, 95% CI 0.485–0.692); cancers diagnosed at 12 months or later were separated at AUC 0.766 (0.671–0.857), with discrimination highest for cancers arising between 12 and 36 months, though that window contains only nine events (AUC 0.863, 0.761–0.953). The two quantities answer different questions and we report the time-dependent one as primary: cumulative/dynamic AUC, which contrasts every case arising by a given horizon with participants still event-free beyond it, rose from 0.577 at 12 months to 0.629 at 24 and 0.636 at 36 months; the window-specific values above instead contrast cases arising within one interval against all event-free controls, and are complementary rather than comparable. Late cases scored above early ones (*P = 0.011*). The direction held within site: lung cancers gave 0.547 within 12 months against 0.818 thereafter, though only eight lung cancers occurred at 12 months or later and that estimate is correspondingly imprecise; non-lung cancers gave 0.649 against 0.738. These findings argue against performance being driven predominantly by prevalent or near-term disease: discrimination sits in cancers arising a year or more after sampling. This distinction is important when interpreting the headline performance metric. PREVALUNG scheduled low-dose computed tomography within seven months of inclusion and the median interval from sampling to lung-cancer diagnosis was 2 months, so the all-cancer sensitivity of 98.3% is carried in part by the cancers this classifier separates least well.

These results compared favorably with established clinical selection approaches. The PLCO_m2012_ threshold yielded AUCs of only 0.700 (95% CI 0.603–0.798) in FLEMENGHO and 0.668 (95% CI 0.593–0.743) in PREVALUNG, with corresponding hazard ratios of 2.46 (95% CI 1.27−4.79) and 2.29 (95% CI 1.26−4.17). In PREVALUNG the NELSON and NLST criteria discriminated poorly as binary scores (AUCs 0.559 and 0.539) and did not separate cancer-free survival. These are eligibility rules, not continuous risk models: the comparison describes how current criteria behave in a cardiovascular smoking population and is not a formal test of them (**Extended data Fig. 8**).

Clonal hematopoiesis of indeterminate potential (CHIP) is associated with an increased risk of LC independently of other clinical parameters besides aging. Targeted sequencing using a panel of 40 genes recurrently mutated in myeloid malignancies was successfully performed on genomic DNA in 85 PREVALUNG patients (detailed in **Supplementary table S4a**) An extended NGS panel including 67 genes and 32 additional genes (among which ATM, ASXL2, CHEK2) was used to screen 40 LC, 67 CVD, and 34 HV patients from the FLEMENGHO cohort (**Supplementary table S4b**). 33.3% and 27.5% of PREVALUNG and FLEMENGHO cancer patients exhibited at least one CHIP mutation with a variant allele frequency (VAF) ≥2% respectively, versus 18.5% and 20% of cancer-free CVD individuals (**Extended data Fig. 9a, b**). These percentages were slightly above the ones described in the literature on ACVD (19/113, i.e., 17% of patients with coronary disease),^57^ despite a similar median age of the two populations. The most commonly mutated genes were DNMT3A, TET2, and TP53 at a VAF ranging from 2 to 60% (**Extended data Fig. 9a, b**). The accumulation of concomitant CHIP mutations as well as the CHIP mutation VAFs were not associated with our cancer risk (**Extended data Fig. 9c, d**).

PREVALUNG also shows directly how the classifier would redistribute imaging. Of 397 classified participants, 83 were ruled out, 286 were medium risk and 28 were high risk. The observed quantity is this: 20.9% of the cohort fell into the very-low-risk stratum, and 98.3% of incident cancers diagnosis were retained outside it. We deliberately stop there. No outcome data support any surveillance interval for the high-risk stratum, scan count is one component of screening cost, and a scan withheld is not exchangeable with a scan added elsewhere.

The intended-use population is narrow and we state it: adults aged 45–75 with daily tobacco exposure and established atherosclerotic cardiovascular disease, in whom low-dose computed tomography is already under consideration. The classifier has not been evaluated in never-smokers, in smokers without cardiovascular disease, in people whose only risk factor is chronic obstructive pulmonary disease, or across ancestries. It is not a general cancer-risk test. Nothing here justifies withholding, deferring or intensifying imaging for any individual: the test is not ready for clinical use, and the endpoint that matters; stage distribution and cancer-specific mortality under biomarker-guided allocation; has not been studied.

## Discussion

This study describes a minimal sequential strategy for stratifying incident cancer risk in people already enriched for atheromatous cardiovascular disease, and reports its reproducibility in a second cohort. The first step uses low O-GlcNAcase (OGA) and interleukin-6 (IL-6) to identify individuals at very low risk; the second applies low chymotrypsin C (CTRC) and high β-microseminoprotein (MSMB) to rule in a small group at high risk among those not excluded. Four proteins and two routinely available clinical variables (i.e., smoking status and personal cancer history) generate three operational strata. The resulting assay is small, transferable in principle and reasonably accurate: on transfer to the PREVALUNG cohort, the complete classifier achieved an AUC of 0.778, while the rule-out group had a negative predictive value of 98.8% and the rule-in group a positive predictive value of 53.6%. Importantly, the plasma proteome was not merely the richest discovery layer but the only omic layer required for prediction. Metabolomics, immunophenotyping, clonal-hematopoiesis analyses and metagenomics illuminated the biology of the score, yet none yielded a single biomarker that reproducibly improved the two-step, four-protein model across cohorts. This negative result is valuable because it limits analytical complexity, missingness and cost without discarding the mechanistic information provided by multi-omics.

The four proteins appear to sample complementary physiological compartments. OGA, an intracellular enzyme that removes O-linked N-acetylglucosamine, was detected across lymphoid populations, and its circulating concentration tracked OGA abundance in T, B and natural killer cells. This accords with the central role of O-GlcNAc cycling in lymphocyte activation and innate inflammatory signalling.^58^ Its correlates connected nutrient sensing and glycan remodeling to cytokine signaling and both adaptive and innate immunity, although further studies will be required to understand the link between circulating plasma OGA levels and the risk of developing cancer. IL-6 is an inducible product of myeloid, lymphoid and stromal cells rather than a tissue-specific secretion; it coordinates acute-phase, TLR, B-cell and T-cell programs. ^59^ Here, it additionally marked the immunosuppressive/proinflammatory SIG1 microbiota configuration, antimicrobial peptides and death-receptor-associated immune turnover. Thus, both rule-out markers report widely distributed immune states, although neither should be considered exclusively immune-derived. CTRC, by contrast, tracked almost uniquely with pancreatic acinar-cell signatures and a coherent digestive-enzyme programme. These analyses are compatible with a pancreatic origin for circulating CTRC; they do not establish one. ^55^ Its network also intersected KRAS-related signaling, but this systems-level association does not establish a direct role for CTRC in KRAS biology. MSMB was far more promiscuous, mapping to pulmonary, vascular, stromal and immune compartments. Its network encompassed cytokine–receptor, IL-6–JAK–STAT3 and TNF signaling, as well as TP53-regulated death genes. TP53 signaling may represent the clearest cell-autonomous component, whereas death-receptor signatures accompanying several proteins more plausibly reflect immune-cell activation or turnover.

These patterns resonate with the dual pathogenesis of lung cancer. One arm is cell autonomous: tobacco carcinogens and inflammation-associated reactive species create mutations, genomic instability and selection of fitter epithelial clones. The other is ecological: transformed cells must escape immunosurveillance. Immune activation is already evident in pre-invasive lung lesions, followed by progressive immune escape before invasion.^60^ The TRACERx study by Swanton’s group subsequently demonstrated neoantigen loss, antigen-presentation defects and other routes of immune evasion in untreated NSCLC.^61^ Inflammation bridges these processes. Recurrent damage and repair generate oxidants, proliferation and selective pressure, while chronic cytokine signaling recruits suppressive myeloid cells and regulatory lymphocytes and disables effective immune predation. At least three classifier components, OGA, IL-6 and MSMB, are embedded in immune or inflammatory networks. CTRC appears orthogonal, perhaps reporting compromised exocrine, digestive or metabolic fitness. Yet low CTRC was unaccompanied by major metabolomic shifts or weight loss, arguing against overt malnutrition, cachexia or clinically manifest pancreatic insufficiency.

At least one of the classifier proteins might be actionable. CTRC and MSMB currently lack compelling preventive or therapeutic evidence, and OGA inhibition may even be counterproductive: pharmacological OGA inhibition increased O-GlcNAcylation and cisplatin resistance in lung-cancer cells. ^62^ IL-6, however, is pharmacologically tractable. IL-6 blockade suppressed tumor promotion, STAT3 activation, proliferation and angiogenesis in murine KRAS-driven lung cancer. ^63^ Nevertheless, no trial has shown that IL-6 inhibition prevents lung cancer, and its pleiotropic functions in host defense preclude treating an elevated risk marker as an indication. The defensible conclusion is therefore that none is a clinically validated interception target, while IL-6 constitutes a biologically credible exception deserving prospective investigation.

Potential intervention points instead emerged from the proteomic field associated with failure of rule-out. Six increased proteins are actionable in principle. GDF15 suppresses antitumor immunity; its neutralization with visugromab restored responses to PD-1 blockade in some patients, including those with NSCLC.^64^ IL-4Rα signaling drives tumor-promoting myelopoiesis, and adding dupilumab to PD-(L)1 blockade reduced circulating monocytes and expanded intratumoral CD8 T cells in a small NSCLC study.^65^ TREM2 marks suppressive macrophages that exclude or disable cytotoxic lymphocytes; its blockade remodels the myeloid compartment and augments checkpoint therapy, while lung-specific work links TREM2-positive macrophages to NK-cell paucity.^66,67^ MAP2K1/MEK1 transmits oncogenic RAS signaling and is druggable, although the negative SELECT-1 trial illustrates that targetability does not guarantee efficacy in KRAS-mutant NSCLC.^68^ Extracellular ACBP/DBI restrains autophagy and immunosurveillance; its neutralization reduced experimental carcinogenesis, making it a candidate interception target.^14,15^ Finally, BCL2L1/BCL-xL protects damaged cells from mitochondrial apoptosis, and its inhibition sensitized lung-cancer models to MEK blockade.^69^ These molecules should be viewed as hypotheses generated by the at-risk proteome, not as therapies selected by the classifier.

Our approach differs from previous lung-cancer risk models in two fundamental respects. First, both cohorts focus on individuals with cardiovascular risk factors or events. Because atheromatous cardiovascular disease and cancer share smoking exposure, ageing, clonal hematopoiesis, inflammation and metabolic dysfunction,^70–72^ comparison within this background should enrich signals that discriminate asymptomatic or future cancer rather than merely recapitulate generic cardiometabolic risk. It does not, however, formally establish lung-cancer specificity. Second, most published models generate one continuous probability from clinical variables or multiplexed biomarkers. Our architecture is deliberately asymmetric: a sensitive first gate identifies a very-low-risk stratum, whereas a specific second gate concentrates risk among the remainder. This structure maps onto distinct clinical actions that remain hypothetical.

Clinical implementation now requires three sequential demonstrations. First, the locked assay and thresholds must be validated in additional prospective cohorts of smokers with comparable cardiovascular profiles, using standardized sampling, low-throughput quantitation methods and prespecified predictive horizons. Second, a randomized trial should test biomarker-guided LDCT allocation: no or less frequent imaging for the ruled-out group, guideline-frequency LDCT for the intermediate group and approximately threefold more frequent LDCT for the high-risk group, with safeguards against interval cancers. Third, this strategy must improve stage distribution and ultimately lung-cancer mortality, quality of life and cost-effectiveness, rather than merely increase diagnoses and procedures. Several limitations bound these conclusions. Also, these biomarkers may be tested as biological surrogates of cancer risk evolution associated with clinical risk reduction following lifestyle changes such as smoking cessation and increase in physical activity. The cohorts are modest and the intervals correspondingly wide, most acutely for the high-risk stratum of 28 PREVALUNG participants. Discovery used incident lung cancer; validation used all incident cancers, fewer than half of them lung. The classifier may therefore report a systemic susceptibility state rather than a lung-specific signal, and lung-specific performance rests on 26 events. FLEMENGHO was assembled as a nested case–control set including healthy non-smoking controls: its apparent discrimination is inflated relative to a population setting. PREVALUNG participants underwent a low-dose chest CT scan within seven months of enrollment; some cancers were therefore pre-existing (prevalent) rather than incident, while remaining asymptomatic. Horizon analyses therefore separated diagnoses within 12 months from later cancers; discrimination was greater for later cancers, but the event counts preclude a definitive separation of susceptibility from occult disease. Predictive values are prevalence-dependent. The four proteins were quantified on a single affinity-proteomic platform, so orthogonal assay validation and recalibration precede any deployment. Competing cardiovascular mortality further complicates absolute risk in this population. Confounding deserves separate mention in a cardiovascular population. Statins, antiplatelet and antihypertensive agents, metformin, corticosteroids, diabetes, renal and hepatic impairment, chronic obstructive pulmonary disease, body composition, recent infection and interval cardiovascular events all plausibly move the proteins we measured. IL-6 is a non-specific inflammatory readout and we have not established what it adds over C-reactive protein and leukocyte counts. Low CTRC may equally reflect exocrine pancreatic, nutritional or alcohol-related states. Personal cancer history is a model covariate, so the cohort includes survivors in whom incident events may be recurrences or second primaries. The two cohorts differ in country, collection era, storage duration and assay batch, and preprocessing constants were transferred without a bridging study. Finally, the cascade is a three-category decision rule, not a calibrated probability model: calibration and net benefit are not yet assessed, and the strata should not be read as calibrated risk levels. The horizon analysis reported above bears directly on this. Discrimination increased rather than decreased with time to diagnosis, a pattern more consistent with a predisposing systemic state than with detection of prevalent asymptomatic disease. We note the converse cost: near-term cancers, which dominate the PREVALUNG lung-cancer set, are the ones the cascade separates least well, and no part of this work supports its use for early detection.

## Supporting information

Doc. 1. REMARK reporting checklist.

Doc. 2. TRIPOD+AI reporting checklist.

## Acknowledgments and Funding

We thank Lena Lindbom, Anissa Lamarque, and Marijana Rucevic from Olink and Vadim Puller, Raynald de Lahondès, and Etienne Formstecher from GMT Science. We thank Bertrand Routy, Thomas Marron, Pedro Romero, Solange Peters, Ignacio Melero, Carlijn M van der Aalst, Harry J de Koning, Tarek Ben-Ahmed and Suzette Delaloge for their criticisms. LZ and GK are supported by the SEERAVE Foundation, the European Union’s Horizon Europe research and innovation program under grant agreement No 101095604 [project acronym: PREVALUNG-EU, project title: Personalized lung cancer risk assessment leading to stratified Interception], the European Union’s Horizon 2020 research and innovation program under grant agreement No 825410 (ONCOBIOME project), Institut National du Cancer (INCa), ANR Ileobiome - 19-CE15-0029-01, ANR RHU5 “ANR-21-5 RHUS-0017” IMMUNOLIFE”, MAdCAM INCA_ 16698, Ligue contre le cancer, the LabEx Immuno-Oncology (ANR-18-IDEX-0001), la direction generale de l’offre de soins (DGOS), Université Paris-Saclay, PACRI network, Ligue contre le Cancer (équipes labellisées, Program “Equipe labelisée LIGUE”; no. EL2016.LNCC (VT/PLP)); Agence Nationale de la Recherche (ANR) – Projets blancs; ANR under the frame of E-Rare-2, the ERA-Net for Research on Rare Diseases; AMMICa US23; Association pour la recherche sur le cancer (ARC); Association “Le Cancer du Sein, Parlons-en!”; Cancéropôle Ile-de-France; Chancelerie des universités de Paris (Legs Poix), Fondation pour la Recherche Médicale (FRM); a donation by Elior; European Research Area Network on Cardiovascular Diseases (ERA-CVD, MINOTAUR); Gustave Roussy Odyssea, Fondation Carrefour; INCa; Inserm (HTE); Institut Universitaire de France; LeDucq Foundation; and the SIRIC Cancer Research and Personalized Medicine (CARPEM). MF is supported by the SEERAVE Foundation and the European Union’s Horizon Europe research and innovation program under grant agreement No 101095604 [project acronym: PREVALUNG-EU, project title: Personalized lung cancer risk assessment leading to stratified Interception]. FM and AD acknowledge the research infrastructure France Exposome. We thank Carlijn M van der Aalst, Harry J de Koning, Sam M Janes, Francesco Passiglia, Silvia Novello, and Morten L. Isaksen for their contribution in the PREVALUNG-EU program.

## Author contributions

**Conceptualization**: MF, DB, NP, GK, and LZ,; **Methodology**: MF, TK, ND, DB, GK, LZ, and NP; **Software**: MF, TP, NP, TD, MD, and AR; **Validation**: NP, MF, and TP; **Formal analysis**: MF, TP, TD, AR, and NP; **Investigation**: MF, DB, AG, LP, SD, FA, and AR; **Resources**: DB, PA, PP, CC, OH, TW, and TK; **Data curation**: MF, DB, TP, TK, and NP; **Writing – original draft**: MF, DB, TP, LZ, NP and GK; **Visualization**: MF, TP, AG, and NP; **Supervision**: MF, DB, NP, GK, and LZ; **Project administration**: MF, DB, MM, TK, and LZ; **Funding acquisition**: LZ.

## Competing interests

LZ is a cofounder of everImmune, the President of everImmune SAB, and holds patents covering the treatment of cancer and the therapeutic manipulation of the microbiota by LBP. LZ has held research contracts with Biomérieux, Daichi Sankyo, Glaxo Smyth Kline, Incyte, Lytix, Kaleido, Pileje, Transgene, 9m, Tusk Pharma, Merus, Roche and Innovate Pharma, and now has current research support from Biomérieux, Daiichi Sankyo, everImmune, and Pileje. LZ is in the SAB of Hookipa. LZ was in the Board of Directors of Transgene and is currently in the SAB of Hookipa. GK has been holding research contracts with Daiichi Sankyo, Eleor, Kaleido, Lytix Pharma, PharmaMar, Osasuna Therapeutics, Samsara Therapeutics, Sanofi, Sotio, Tollys, Vascage and Vasculox/Tioma. GK has been consulting for Reithera. GK is on the Board of Directors of the Bristol Myers Squibb Foundation France. GK is a scientific co-founder of Osasuna Therapeutics, Samsara Therapeutics and Therafast Bio. GK is the inventor of patents covering therapeutic targeting of aging, cancer, cystic fibrosis and metabolic disorders. GK’s brother, Romano Kroemer, was an employee of Sanofi and now consults for Boehringer-Ingelheim. The funders had no role in the design of the study; in the writing of the manuscript, or in the decision to publish the results. N.P. is CEO, co-founder and shareholder of TheraPanacea, and holds advisory and/or equity interests in Reveal Genomics, Nuclivision and Artedrone. He is a member of the Scientific Council of Safran. DB is consultant for Kenvue and received research grant from Kenvue, Astra Zeneca and Medtronic. PA received speaker fees and research grants from Astra-Zeneca, Netcancer, MSD, Janssen, Takeda, Novonordisk, as well as travel grants from Roche, Pierre Fabre, Eli Lilly, Pfizer.

## Materiel and methods

The study is reported per TRIPOD+AI^73^ and REMARK^74^; the completed checklists are supplied as Supplementary files.

### Cohort description

#### Medical centers and regulatory approvals for translational research

**The Flemish study on Environment, Genes and Health Outcomes (FLEMENGHO)** received ethical approval from the Ethics Committee of the University of Leuven (S64406). The FLEMENGHO cohort represents a random population sample stratified by sex and age from a geographically defined area in northern Belgium. The initial response rate was 78%. All participants provided written informed consent prior to study participation. The participants (n=3,343) were repeatedly examined at a local examination center. During regular re-examination, standardized and validated questionnaires were administered to collect detailed information about each participant’s medical history, use of medication, socio-economic class and lifestyle including information on smoking habit. Phenotyping at the examination center included clinical and anthropometric measurements. Blood samples (i.e. serum) are available. Every year, the vital status of all registered persons and any changes in their place of residence are requested from the National Register in Brussels. The competent service of the Flemish Community then provides the immediate and underlying causes of death, as registered on the death form. To confirm the diagnoses on the questionnaires (non-fatal endpoints) or on the death certificate, the study team contacts the GP or the treating specialist, provided that the participants gave their prior consent.^43^ Participants eligible for inclusion in this retrospective validation study were selected from the FLEMENGHO cohort. For 38 selected lung cancer cases the blood samples were of a good quality for proteomics/metabolomics. These were samples obtained from consequently examined participants starting from 1996 till 2015. Median follow-up time between examination/ blood sampling and incidence of lung cancer is 4.3 years (interquartile range, 1.9 to 6.9 years). We matched these cases by age, sex, body mass index (BMI), with 53 healthy non-smoker controls from the same population. We also included 65 subjects with ACVD (atherosclerosis events) who did not develop lung cancer (or any other forms of cancer) during follow-up period (4.9 years). These ACVD controls have been paired on age, gender, BMI, and by tobacco use (current or ex-smokers) with lung cancer cases (**Supplementary Table 1**).

**Prevalence of Lung Cancer (PREVALUNG)** study is a monocentric and prospective study enrolling adult patients in their follow up post-acute cardiovascular events. The inclusion criteria were (1) age 45-75 years old, (2) history of atheromatous cardiovascular disease (ACVD) and (3) history of daily tobacco consumption for 10 years prior to onset of ACVD. Exclusion criteria were symptoms of lung cancer (LC), existing follow-up for pulmonary nodule, fibrosis, pulmonary hypertension, resting dyspnea and active pulmonary infectious disease. 508 patients were included in this study from November 18th, 2019 up to May 18th, 2021 at Marie Lannelongue hospital (Le Plessis-Robinson, France).^44^ After inclusion, patients were scheduled for a low-dose chest CT-scan within 7 months. Blood and feces samples were collected concomitantly with the CT-scan. Each patient was scheduled for a follow-up by phone visits at month 3, month 6 and month 12 after the CT-scan. LC prevalence was the primary endpoint. National Lung Screening Trial and Dutch-Belgian Randomized Lung Cancer Screening Trial (NELSON) trial eligibility criteria, radiation, positive screening, false positivity, rate of localized LC diagnosis, quality of life with the Short Form 12 (SF-12) and anxiety with the Spielberger State-Trait Anxiety Inventory A and B (STAI-YA and STAI-YB, respectively), smoking cessation and onset of cardiovascular and oncological events within 1 year of follow-up were recorded. A case-control study nested in the cohort was performed to identify clinical or biological candidate biomarkers of LC or tobacco-associated cancer. The study was approved according the French Jardé law; the study is referenced at the French ‘Agence Nationale de Sécurité du Médicament et des Produits de Santé’ (reference ID RCB: 2019-A00262-55) and registered on clinicaltrial.gov (NCT03976804). Proteomics data were available for 397 patients and clinical follow-up for cancer diagnosis and ACVD events was up to 5 years after inclusion with a median follow-up time of 4 years (**Supplementary Table 2**).

### Method details

#### Collection of serum, plasma and PBMC samples from blood

Blood samples were drawn from patients. Serum was collected after centrifugation of serum separator tubes. Plasma was collected after centrifugation of lithium heparin tubes without Ethylenediaminetetraacetic acid (EDTA), and peripheral blood mononuclear cells (PBMC) were separated using a density gradient medium. Serum, plasma, as well as whole blood, when available, were stored at −80°C or lower for long-term storage. Only samples with a good quality for subsequent analysis were selected. PMBCs were stored in liquid nitrogen (−196°C) in a medium composed of 90% fetal bovine serum (FBS) and 10% dimethylsulfoxide (DMSO).

#### Collection of patient stools, DNA extraction and sequencing (PREVALUNG)

Fecal samples were prospectively collected at the same time as LDCT LC screening following the International Human Microbiome Standards (IHMS) guidelines. For metagenomic analysis, the stools were processed for total DNA extraction and sequencing with Ion Proton technology following MetaGenoPolis (INRAE) France, as previously reported.^52,75–77^

#### Proteomics analysis on serum

For each patient, 100µl of serum was provided to Olink Proteomics AB and the full Explore 3072 panel was performed, which consists in eight separated panels: Oncology, Oncology II, Cardiometabolic, Cardiometabolic II, Inflammation, Inflammation II, Neurology, and Neurology II. Samples randomization was done using Olink Analyze R package. Relative quantification was calculated from Ct values and Log2 scaled into normalized protein expression (NPX). Before conducting several statistical analysis and machine learning models, proteomics features were first filtered by selecting proteins measured in both cohorts

#### Metabolomics analysis on plasma or serum

For each patient, 50 μL of crude plasma or serum was mixed with 500 μL of ice-cold extraction mixture (methanol/water, 9/1 spiked with a cocktail of internal standards) and centrifuged after 5 min of incubation (10 min at 15,000g, 4°C). 150 µL aliquots of supernatant were transferred to glass vials and evaporated to be used for Gas Chromatography combined with Mass Spectrometry (GC-MS). Then, 50 μL of methoxyamine (20 mg/ml in pyridine) was added on dried extracts, which were stored at room temperature in the dark, overnight. The day after, 80 µL of N-Methyl-N-trimethylsilyl-trifluoroacetamide were added, and the final derivatization was made at 40°C during 30 minutes. Samples were then transferred into vials and directly injected into the GC-MS system. GC-MS/MS was performed with a 7890A gas chromatograph (Agilent Technologies, Waldbronn, Germany) coupled with a triple quadrupole 7000C (Agilent) equipped with a high sensitivity electronic impact source (EI) operating in positive mode. This method was inspired by developments reported previously.^78^ The scan mode was the MRM for biological samples. Peak detection and integration of the analytes were performed using the Agilent Mass Hunter quantitative software (B.07.01).^79^ All targeted treated data were merged and cleaned with a dedicated R package (version 4.0) (@Github/Kroemerlab/GRMeta). A daily qualification of the instrumentation was set up with automatic tune. These qualifications were completed with double injections of standard mixes, at the beginning and at the end of the run, in addition to blank samples to control the background impurities. Moreover, pools of tested samples were used to check the column before the analysis with the proper biological matrix. The same pool was re-injected during the batch to monitor and correct analytical bias occurring through the acquisition and post-acquisition treatment of the signal (m/z, retention time, and sensitivity drifts). For each analyte, the corrected peak area was considered to be proportional to its concentration

#### Orthogonal validation of OGA protein levels in circulating immune cells (PBMCs)

##### OGA protein levels in circulating immune cells measured by flow cytometry

Peripheral blood mononuclear cells (PBMCs) were analyzed by multiparameter flow cytometry to assess intracellular OGA levels. PBMCs were seeded into V-bottom 96-well plates and centrifuged at 1200 rotations per minute (rpm) for 4 min at 4°C. Cells were resuspended in 50 μL per well of a surface-staining antibody cocktail prepared in FACS buffer (PBS 1X + 2% FBS + 0.5M EDTA). The antibody cocktail consisted of viability dye APC-Cy7 (ThermoFisher Scientific, cat. no. L34994) (1:1000), anti-CD56-BUV737 (1:200) (clone NCAM16.2; BD Biosciences, cat. no. 564447), anti-CD3-PerCP-Cy5.5 (1:50) (clone UCHT1; BioLegend, cat. no. 300429), anti-CD4-BV785 (1:200) (clone RPA-T4; BioLegend, cat. no. 300553), anti-CD8-BV421 (1:100) (clone SK1; BD Biosciences, cat. no. 740093), anti-CD19-PE-CF594 (1:100) (clone HIB19; BD Biosciences, cat. no. 562294) and anti-CD14-BV650 (1:100) (clone M5E2; BioLegend, cat. no. 301836). For 1900 μL of staining mixture, 1.9 μL of viability dye, 9.5 μL of anti-CD56, 38 μL of anti-CD3, 9.5 μL of anti-CD4, 19 μL of anti-CD8, 19 μL of anti-CD19 and 19 μL of anti-CD14 were combined with 1,784 μL of FACS buffer. Cells were incubated for 20-30 min at 4°C and subsequently washed with 100 μL of FACS buffer. Plates were centrifuged at 1200 rpm for 2 min at 4°C.

For intracellular staining, cells were fixed and permeabilized by incubation with 100 μL of Cytofix/Cytoperm (BD Biosciences, cat. no. 554714) for 20 min at 4°C. Cells were then washed with 80 μL of 1× Perm/Wash buffer (BD Biosciences, cat. no. 554723). Cells were centrifuged at 2000 rpm for 2 min at 4°C and resuspended in 100 μL of 1× Perm/Wash buffer before overnight incubation at 4°C.

The following day, cells were centrifuged at 2000 rpm for 2 min at 4°C and resuspended in 50 μL of the corresponding intracellular antibody cocktail. For OGA detection, cells were stained with purified anti-OGA antibody (1:100) (clone 11E7; Invitrogen, cat. no. MA572964,). For each staining condition, the antibody cocktail was prepared in a final volume of 1900 μL using 1× Perm/Wash buffer. Cells were incubated for 30 min at room temperature and washed with 100 μL of 1× Perm/Wash buffer, followed by centrifugation at 2000 rpm for 2 min at 4°C.

A secondary staining step was performed using AF647-conjugated anti-rabbit IgG (1:50) (Jackson Immuno Research, cat. no. 711-606-152) prepared in 1× Perm/Wash buffer. Cells were incubated for 30 min at room temperature, washed with 100 μL of 1× Perm/Wash buffer and centrifuged at 2000 rpm for 2 min at 4°C. Finally, cells were resuspended in 70 μL of FACS buffer and transferred to microcentrifuge tubes for flow cytometric acquisition.

Stained samples were acquired using a BD LSR Fortessa 5L flow cytometer. Flow cytometry and mean fluorescence intensity (MFI) data were analyzed using *FlowJo* v10.8.1 (BD Biosciences).

| REAGENT OR RESOURCE | SOURCE | IDENTIFIER |
| --- | --- | --- |
| REAGENT |  |  |
| Viability dye | ThermoFisher Scientific | Cat. No. L34994 |
| Cytofix/Cytoperm | BD Biosciences | Cat. No. 554714 |
| Perm/Wash buffer | BD Biosciences | Cat. No. 554723 |
| Antibodies |  |  |
| anti-OGA antibody | Invitrogen | Cat. No. MA572964 |
| anti-CD56-BUV737 | BD Biosciences | Cat. No. 564447,<br>RRID: AB_2744432 |
| anti-CD19-PE-CF594 | BD Biosciences | Cat. No. 562294,<br>RRID: AB_11154408 |
| anti-CD14-BV650 | BioLegend | Cat. No. 301836,<br>RRID: AB_2563799 |
| anti-CD3-PerCP-Cy5.5 | BioLegend | Cat. No. 300429,<br>RRID: AB_893301 |
| anti-CD4-BV785 | BioLegend | Cat. No. 300553,<br>RRID: AB_2564381 |
| anti-CD8-BV421 | BD Biosciences | Cat. No. 740093,<br>RRID: AB_2739852 |
| PE-conjugated anti-rabbit IgG | Jackson ImmunoResearch | Cat. No. 711-606-152,<br>RRID: AB_2340625 |

##### Orthogonal validation analyses between plasma OGA protein levels and circulating immune-cell OGA expression

Associations between plasma OGA protein abundance quantified by Olink proteomics (NPX) and OGA expression measured by flow cytometry (median fluorescence intensity, MFI) in multiple immune-cell populations were evaluated using Spearman’s correlation coefficient (rho) in 37 patients from the PREVALUNG cohort. For each immune-cell population, a two-sided Spearman correlation test was performed independently. Correlation coefficients (rho) and nominal *P* values were calculated. Results were summarized using a bubble plot in which the x-axis represents the Spearman correlation coefficient (rho), dot size corresponds to −log_10_(*P* value), and dot color reflects the direction of the correlation, with blue indicating negative correlations and red indicating positive correlations. Non-significant (*ns*) correlations are indicated in grey.

Also, participants were dichotomized according to the median plasma OGA protein level measured by Olink proteomics in the PREVALUNG cohort, generating two groups (<MEDIAN and >MEDIAN). For each immune-cell population, OGA expression quantified by flow cytometry (MFI) was compared between groups using a two-sided Wilcoxon rank-sum test. Boxplots were used to display the median and interquartile range, with each dot representing individual participants.

R Packages: *dplyr* (v1.2.1), *ggplot2* (v4.0.3), *ggpubr* (v0.6.3), and *stats* (v4.5.2).

##### OGA protein levels in circulating immune cells measured by Jess Automated Western Blot System

OGA protein levels were assessed in frozen PBMC dry pellets using the Jess Automated Western Blot System (ProteinSimple, Bio-Techne), a capillary-based automated Western blotting platform. Frozen PBMC pellets were lysed and protein extracts were prepared according to the manufacturer’s instructions. Protein samples were diluted in fluorescent master mix containing dithiothreitol and fluorescent molecular weight standards, denatured, and loaded onto Jess capillary cartridges.

OGA was detected using a rabbit monoclonal anti-OGA antibody (MGEA5, clone E9C5U; Cell Signaling Technology, cat. no. 60406) at the optimized dilution. Following electrophoretic separation and protein immobilization within the capillaries, samples were sequentially incubated with the primary antibody, HRP-conjugated anti-rabbit secondary antibody and chemiluminescent detection reagents according to the manufacturer’s protocol. Electropherograms and quantitative peak areas were generated using Compass for Simple Western software (ProteinSimple, Bio-Techne).

To account for differences in protein loading between samples, OGA signal was normalized to total protein measured in each capillary using the Jess Protein Normalization Assay. This approach quantifies the total protein content of each sample independently of housekeeping proteins and enables normalization of the OGA peak area to the corresponding total protein signal.

Two independent analytical runs were performed using frozen PBMC dry pellets. Run 1 included 17 participants and run 2 included 16 participants. Representative electropherograms from each analytical run were generated using Compass software.

#### Immunophenotyping by spectral flow cytometry

Cryopreserved peripheral blood mononuclear cells (PBMCs) were stored in fetal calf serum (FCS)/dimethyl sulfoxide (DMSO) and aliquots containing 5 to 10×10^6^ cells were thawed, washed, and resuspended in FACS buffer (Brilliant Stain Buffer Plus, BD Biosciences, supplemented with 5% FCS and 2 mM EDTA). Cell viability was assessed using the LIVE/DEAD™ Fixable Blue Dead Cell Stain Kit (Invitrogen), and dead cells were excluded from subsequent analyses.

Cells were stained with a panel of fluorophore-conjugated antibodies targeting CCR7 (2-L1-A), CADM-1/TSLC1 (3E1), CCR4 (1G1), CCR6 (11A9), CCR9 (L053E8), CD11b (M1/70), CD123 (6H6), CD127 (HIL-7R-M21), CD14 (63D3), CD141 (M80), CD16 (3G8), CD163 (GHI/61), CD169 (7-239), CD19 (HIB19), CD1c (F10/21A3), CD206 (15-2), CD25 (M-A251), CD3 (UCHT1), CD301 (H037G3), CD4 (SK3), CD40 (5C3), CD45 (2D1), CD45RA (HI100), CD5 (UCHT2), CD69 (FN50), CD8α (SK1), CD86 (FUN-1), CD88 (S5/1), CD89 (A59), CTLA-4 (14D3), CXCR3 (1C6/CXCR3), CXCR5 (RF8B2), FcεRIα (AER-37), HLA-DR (L243), LPAM-1 (FAB10078V), PD-1 (EH12.1), PD-L1 (MIH1), and SLAN (DD-1). Antibodies were obtained from BD Biosciences, R&D Systems, MBL, BioLegend, Miltenyi Biotec, and Cytek.

Stained samples were acquired using a Cytek Aurora five-laser spectral flow cytometer (Cytek Biosciences). Flow cytometry data were analyzed using *FlowJo* v10.8.1 (BD Biosciences).

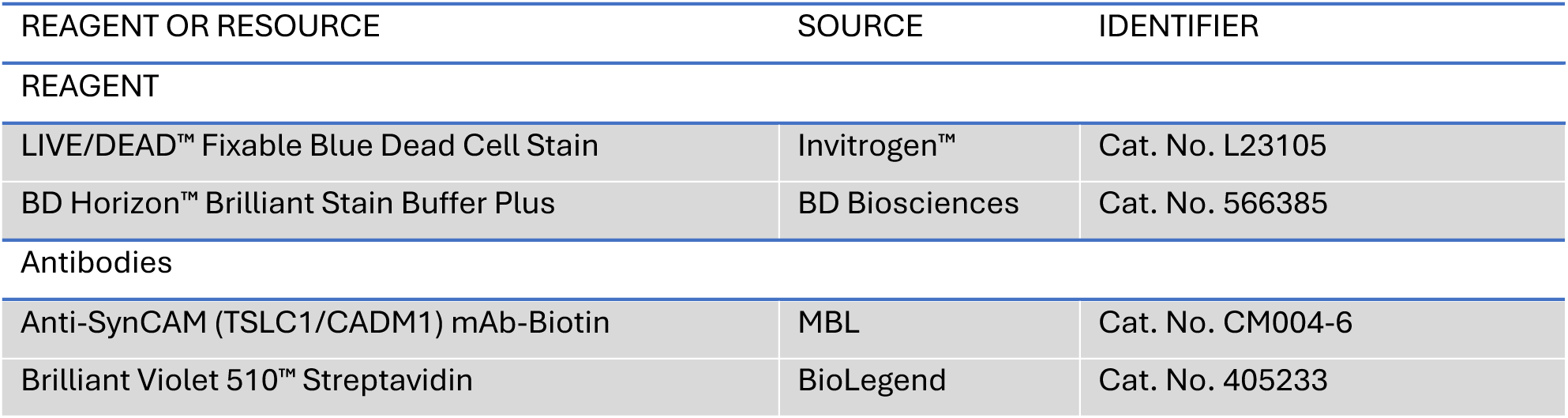
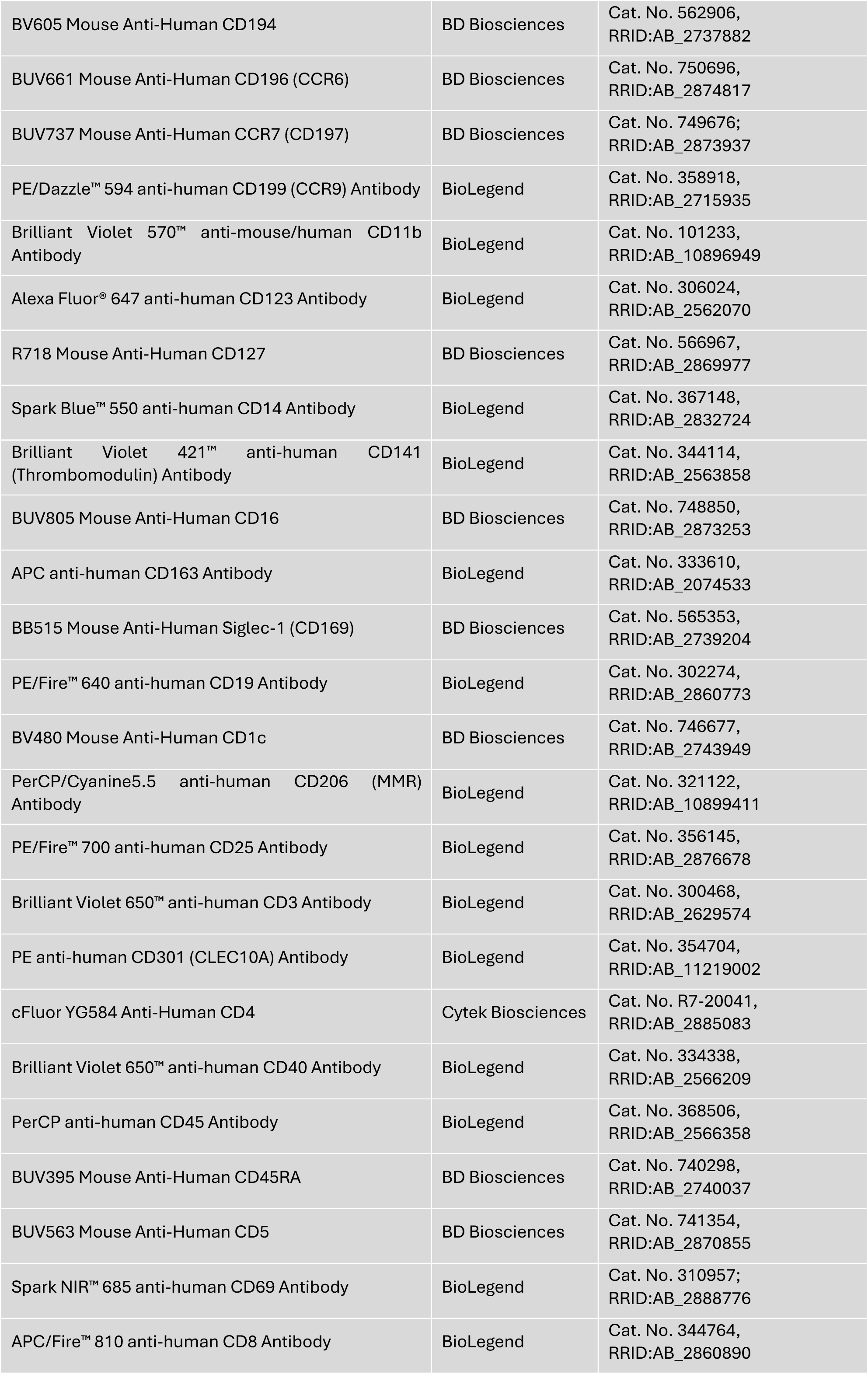
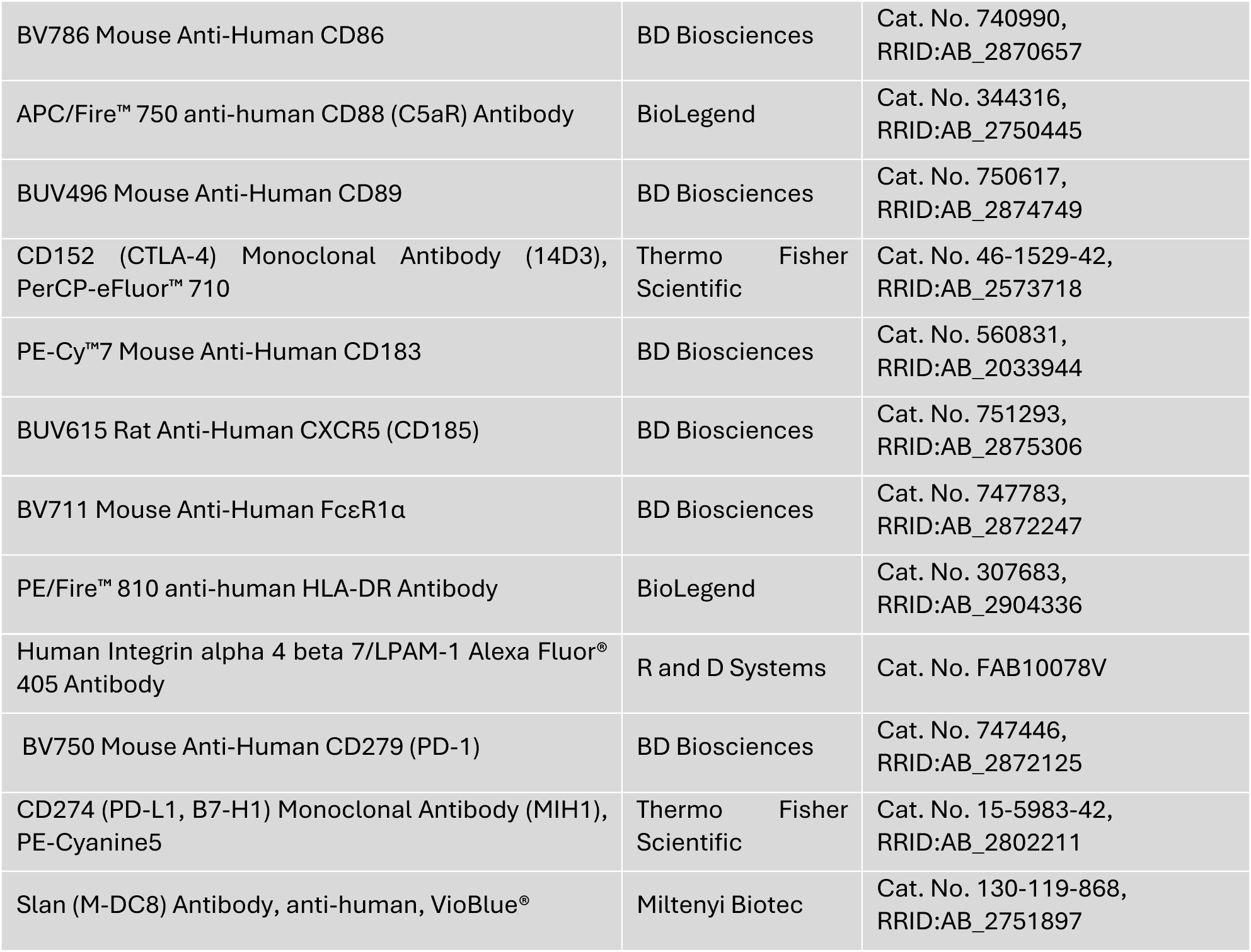

##### Manual gating strategies

Flow cytometric data were pre-processed by sequential exclusion of debris, doublets, and dead cells, followed by selection of viable CD45^+^ cells (refer to **Supplemental Fig. 3**). New .fcs files containing only viable CD45^+^ cells were generated using FlowJo v10.8.1 and used for subsequent dimensionality reduction and automated cell-type annotation analyses.

##### Dimensionality reduction by UMAP

Uniform Manifold Approximation and Projection (UMAP) was performed on viable CD45^+^ cells using the *Scanpy* package^80^ for dimensionality reduction and visualization of the immune-cell landscape.

##### Automated cell-type annotation by Scyan

Cell-type annotation was performed using the Single-cell Cytometry Annotation Network (*Scyan*)^81^, a biology-driven model that incorporates prior biological knowledge of cell types for automated annotation. *Scyan* identified nine major immune-cell populations: basophils, natural killer (NK) cells, B cells, monocytes, dendritic cells, CD4^+^ T cells, CD8^+^ T cells, double-positive (DP) T cells, and double-negative (DN) T cells (refer to **Supplemental Fig. 3**). Following automated annotation, individual .fcs files corresponding to each of the nine major populations were generated for each participant. Subpopulations were subsequently identified by manual gating using *FlowJo* v10.8.1, and their relative frequencies were retained for subsequent statistical analyses.

### Targeted deep sequencing and CHIP-related mutation analysis

Clonal hematopoiesis of indeterminate potential (CHIP) related-mutations were assessed by targeted deep sequencing of the exonic regions of 40 genes commonly mutated in myeloid malignancies. DNA from PBMC was extracted with QIAamp DNA Mini Kit (Qiagen). Ion AmpliSeqTM Custom Panel Primer Pools (Life Technologies) were used (10 ng of gDNA per primer pool) to perform multiplex PCR but the protocol to generate libraries was modified by adding indexed paired-end adaptors (NEXTflex, Bioo Scientific) to sequence on an Illumina Novaseq 6000 SP and MiSeq flow cells using the onboard cluster method, as paired-end sequencing (2×250 bp reads) (Illumina, San Diego, CA). Briefly, PCR amplicons after Fupa digestion were end-repaired, Supplemental with an ‘A’ base on the 3’end, ligated with indexed paired-end adaptors (NEXTflex, Bioo Scientific) using the Bravo Platform (Agilent), amplified by PCR for 6 cycles and purified with AMPure XP beads (Beckman Coulter).

Myeloid panel (165 kb) gene name and targeted exons are indicated: ASXL1 (NM_015338.6): 11-15 12; BCOR (NM_017745.6): 2-15; BCORL1 (NM_021946.4): 1-12; BRAF (NM_004333.6): 15; CALR (NM_004343.3): 8-9; CBL (NM_005188.4): 1-9; CEBPA (NM_004364.4): 1; CSF3R (NM_156039.3): 13-17; CSNK1A1 (NM_001892.6): 2-4; CUX1 (NM_181552.4): 1-24 and (NM_001913.5): 1-23; DDX41 (NM_016222.4): 1-17; DNMT3A (NM_175629.2): 2-23 and (NM_153759.3): 1-19; ETNK1 (NM_018638.5): 2; ETV6 (NM_001987.5): 1-8; EZH2 20 (NM_004456.5): 2-20; FLT3 (NM_004119.3): 14-20; GATA2 (NM_032638.5): 4-6; HRAS (NM_005343.4): 2-5; IDH1 (NM_005896.3): 4, 6; IDH2 (NM_002168.3): 4, 7; JAK2 (NM_004972.3): 8, 12-15; KDM6A (NM_021140.3): 1-29; KIT (NM_000222.2): 1-21; KRAS (NM_004985.5): 1-4; MPL (NM_005373.3): 1-12; MYD88 (NM_002468.5): 4; NPM1 (NM_002520.6): 11; NRAS (NM_002524.5): 1-4; PHF6 (NM_032458.3): 2-10; PPM1D 25 (NM_003620.4): 6; PTEN (NM_000314.8): 1-9; PTPN11 (NM_002834.4): 1-15; RAD21 (NM_006265.3): 2-14; RIT1 (NM_006912.6): 4-5; RUNX1 (NM_001754.4): 1-9; SETBP1 (NM_015559.3): 4-5; SF3B1 (NM_012433.3): 1-25; SRSF2 (NM_003016.4): 1-2; STAG2 (NM_006603.5): 2-33; TET2 (NM_001127208.2): 3-11; TP53 (NM_000546.5): 2-11; U2AF1 (NM_006758.2): 2, 6; WT1 (NM_024426.6): 1, 7; ZRSR2 (NM_005089.3): 1-11.

Genome DNA sequence and annotations were download from Ensembl. Pyfaidx^82^ was used to filter non-cannonical chromosomes. Agat^83^ was used to correct common issues found in Ensembl genome annotation files, filter non-cannonical chromosomes, and remove transcripts with TSL being equal to NA. Samtools^84^ and Picard^85^ were used to index genome sequences. Raw fastq file quality was assessed with FastQC^86^. Raw fastq files were trimmed using Fastp^87^. Cleaned reads were aligned over indexed Ensembl genome with Bowtie 2^88^. Sambamba^89^ was used to sort, filter, mark duplicates, and compress aligned reads. Quality controls were done on cleaned, sorted, deduplicated aligned reads using Picard^85^ and Samtools^84^. Goleft^90^, RSeQC^91^ and ngsderive^92^ were used to assess quality of mapping. Read groups were re-defined over aligned reads with Picard^85^. GATK Mutect2^93^ was used to perform germline calling, following the ‘GATK Best practices’^94^ and the method described in the GATK germline best practices publication^95^. VariantEval and BCFTools^96^ were used to assess variant calling quality. All quality reports produced during both trimming and mapping steps have been aggregated with MultiQC^97^. The whole pipeline was powered by Snakemake^98^.

### NGS data analysis

Somatic variants with at least 10 mutated reads and a variant allele frequency (VAF) ≥ 2% were reported and retained for further analysis. All pathogenic or likely pathogenic variants were manually checked using Integrative Genomics Viewer (IGV) software. Clinical interpretation was supported by public variant databases such as COSMIC (Catalogue Of Somatic Mutations In Cancer), cBioPortal, and ClinVar. The PredictSNP consensus classifier was utilized to refine the functional impact prediction of missense variants,^99^ especially if not listed in the above-mentioned databases.

### Statistical analysis

*Statistical analyses and data visualization were performed using R (R Foundation for Statistical Computing, Vienna, Austria) and Python*.

#### Machine-learning signature

The signature was derived from FLEMENGHO plasma proteomics through a three-stage discovery pipeline (consensus multivariate ranking → mRMR shortlisting → classifier-coupled ablation) designed for the 2,910 candidate proteins for 156 participants and 38 incident cancers. The three stages separate three problems: estimating per-feature multivariate predictive content (consensus multivariate ranking), controlling redundancy across features (mRMR shortlisting), and selecting a compact signature against a deployment objective (classifier-coupled ablation) (**Figure 1**). All stages were run on FLEMENGHO. The pipeline was applied twice to build a sequential two-arm cascade. In the first pass, the full pipeline was run on the whole FLEMENGHO cohort to derive the rule-out arm. Participants cleared by the rule-out test were then set aside, and the pipeline was run a second time on the participants not cleared to derive the rule-in arm.

##### Data curation and standardisation

Circulating protein concentrations were measured by Olink proximity extension assay (Explore 3072; eight panels, 2,966 protein assays per sample) and reported as normalized protein expression (NPX) on a log2 scale, following Olink’s standard sample-preparation, plate-level normalization and cross-plate bridging QC. Proteins and samples failing QC were removed, retaining ≈2,910 cascade-eligible proteins. Missing NPX values were imputed with FLEMENGHO training-set medians. Continuous features were standardized with a StandardScaler estimated on the FLEMENGHO training partition and applied to PREVALUNG. Two clinical covariates including smoking status and personal cancer history were encoded identically across the two cohorts. The outcome was any incident cancer during follow-up (Control vs Cancer). Every incident cancer used for development in FLEMENGHO was a lung cancer. In PREVALUNG the outcome comprised all incident cancers: 26 lung, 14 other tobacco-related, 18 non-tobacco-related. Validation is therefore reported for all cancers and, separately, for each subgroup.

##### Stage 1 — Consensus multivariate ranking

Each protein received a scalar consensus relevance score estimating its multivariate predictive information about the outcome, aggregated over B = 1,000 stratified bootstrap replicates of FLEMENGHO. Five complementary importance estimators with distinct failure modes were computed per bootstrap: (i) bootstrap-stabilized LASSO selection frequency; (ii) random-forest out-of-bag permutation importance; (iii) mean absolute TreeSHAP attribution; (iv) ℓ2-penalised (ridge) logistic-regression coefficient magnitude; and (v) LightGBM split-gain. Each component was median-aggregated across bootstraps and percentile-normalized to [0, 1]; the five percentile scores were pooled by equal-weight arithmetic mean to give the consensus score. The majority direction of effect (up- or down-regulated in the cancer class) was recorded per protein. The top 50 proteins by consensus score were retained as the Stage-2 candidate pool.

##### Stage 2 — mRMR shortlisting

The top-50 pool was reordered by the Minimum Redundancy Maximum Relevance (mRMR) criterion of Peng, Long and Ding to penalize features duplicating information already captured. mRMR was run in its Mutual Information Difference (MID) form, substituting the Stage-1 consensus score for per-feature relevance and the absolute Spearman rank correlation for pairwise redundancy (redundancy weight λ = 1.0), the default appropriate for heavy-tailed, non-Gaussian NPX distributions. Features were selected sequentially: at each step the protein maximizing relevance minus mean redundancy against the current selection was added, to produce the ordered shortlist used as the ablation starting set. Selection is deterministic given the pool, similarity matrix and λ; ties break by feature index.

##### Stage 3 — Classifier-coupled ablation and locking

Within each pass, the 20-feature mRMR shortlist, augmented with the two locked clinical covariates (smoking status and personal cancer history), was refined by classifier-coupled backward elimination in the robust variant. The second pass produced the rule-in arm and ran in the FLEMENGHO subset not cleared by the rule-out test (n = 61, 35 events). That subset was itself defined by a model fitted to the same outcome: the rule-in arm was developed in an outcome-dependent sample. We state its effective sample size and event count so the dependency can be judged. Candidate signatures were scored with a soft-voting ensemble of Gaussian Naïve Bayes and L2-regularised logistic regression, evaluated over repeated stratified 80/20 splits of FLEMENGHO with standard scaling re-fit within each split; the selection metric was ROC AUC. Backward elimination returned the full performance–parsimony trajectory and the beam of competing signatures at each size, from which the two-protein arm signature was selected against the arm’s deployment objective.

The rule-out threshold (26.13%) was the mean of 11 thresholds, each derived from a 5-fold stratified cross-validation of FLEMENGHO with a different random seed (seeds 1000–1010), selecting in each replicate the highest threshold attaining sensitivity ≥ 0.925 on the out-of-fold predictions. The rule-in threshold (68.73%) was derived identically, with a sensitivity constraint of ≥ 0.40 applied to the FLEMENGHO subset not cleared by the rule-out test. Both thresholds were fixed, with the imputation constants, scaler and classifier, before PREVALUNG outcomes were examined for performance.

#### Signature validation on PREVALUNG

The locked pipeline including preprocessing constants, classifier, stage assignments and both thresholds developed on FLEMENGHO cohort was applied to the PREVALUNG transport cohort. The resulting scores were used to classify participants according to the prespecified rule-out and rule-in criteria. Discrimination of each stage score was summarized by ROC AUC with 1,000-resample percentile-bootstrap 95% confidence intervals.

#### Visualization of the rule-out and rule-in scores distribution

Participants in each cohort were categorized as Control or Cancer according to the cancer diagnosis during the follow-up. The distribution of the rule-out and rule-in scores were compared between the two groups using a two-sided Wilcoxon rank-sum test, a non-parametric test appropriate for comparing independent groups without assuming normality.

Two complementary graphical representations were generated: (i) a kernel density estimation (KDE) using a Gaussian kernel with a bandwidth of 5 to visualize the continuous distribution of the rule-out score; (ii) a dot plot displaying individual observations, overlaid with boxplots summarizing the median and interquartile range.

A predefined rule-out threshold (26.13%) and rule-in threshold (68.73%) was represented by a vertical dashed line in both panels.

R Packages: *dplyr* (v1.2.1), *ggplot2* (v4.0.3), *ggbeeswarm* (v0.7.3), *ggpubr* (v0.6.3), and *patchwork* (v1.3.2)

#### Evaluation and visualization of rule-out and rule-in score classifications

Participants were classified according to their observed clinical status (Control or Cancer) during the follow-up and their predicted rule-out category (Rule-out or At risk) or combined with rule-in score category (Rule-out, Medium risk or High risk). A contingency table was generated to summarize the distribution of participants across the different classifications. The confusion matrix was visualized as a heatmap in which each cell represented the number of participants corresponding to a specific combination of observed and predicted categories. Cell color intensity was proportional to the observed frequency, and absolute counts were displayed within each cell. The contingency table was represented as a stacked proportional bar chart. For each predicted category, the relative proportion of participants belonging to each clinical group was displayed, while absolute participant counts were shown within each bar segment.

The association between clinical status and rule-out score alone or combined with rule-in score classification was evaluated using Pearson’s chi-squared test. This test was applied when all expected cell counts exceeded five; otherwise, Fisher’s exact test would be considered. Expected cell counts were calculated from the contingency table to verify the assumptions of the chi-squared test.

R Packages: *dplyr* (v1.2.1), *ggplot2* (v4.0.3), *scales* (v 1.4.0).

#### Correlation analyses between plasma OGA abundance and HBP-related proteins and metabolites

To investigate the molecular environment associated with plasma OGA abundance, Spearman correlation analyses were performed between plasma OGA protein levels (NPX) and curated panels of proteins or metabolites related to the hexosamine biosynthetic pathway (HBP), quantified by Olink proteomics and mass spectrometry, respectively, in the FLEMENGHO and PREVALUNG cohorts. The protein panel included enzymes involved in O-GlcNAc cycling, HBP metabolism, monosaccharide interconversion, nucleotide-sugar synthesis, and glycosyltransferase-mediated glycosylation. The metabolite panel comprised intermediates of carbohydrate metabolism, glutamine-related metabolites, acetylated amino acids, acetylated polyamines, and additional metabolites participating in metabolic pathways connected to HBP activity. For each protein or metabolites, a two-sided Spearman correlation coefficient (rho), and *P* value were calculated independently. Results were summarized as a bubble plot in which the x-axis represents the Spearman correlation coefficient, point size corresponds to −log_10_(*P* value)), and point color indicates the direction of statistically significant correlations (blue, negative; red, positive), while non-significant (*ns*) associations are displayed in grey.

R Packages: *dplyr* (v1.2.1), *ggplot2* (v4.0.3), and *stats* (v4.5.2).

#### Pathway over-representation analysis associated with circulating IL6

Spearman correlation analyses were performed between plasma IL6 protein abundance and proteins consistently associated with IL6 across the FLEMENGHO and PREVALUNG cohorts. The protein panel corresponds to the intersection of proteins significantly correlated with IL6 after Benjamini–Hochberg false discovery rate (FDR) correction in both cohorts and with absolute rho value >0.3.

##### Over-representation analysis (ORA) of pathways

To identify biological pathways consistently associated with circulating IL6, the over-representation analysis (ORA) was performed using the *clusterProfiler* package (v 4.18.4) in R. The background universe consisted of all proteins quantified by the Olink Explore 3072 platform after removing panel-specific suffixes and duplicate protein identifiers. Gene sets were retrieved from the Molecular Signatures Database (MSigDB) using the *msigdbr* package (v 26.1.0), including the Hallmark (H), KEGG Legacy (C2:CP:KEGG_LEGACY), and Reactome (C2:CP:REACTOME) collections. ORA was performed using the *enricher()* function and enriched pathways (*P* value < 0.05) were subsequently ranked according to the gene ratio. The gene ratio was calculated as the proportion of input proteins assigned to each enriched pathway, and dot size represents the number of proteins contributing to pathway enrichment.

R packages *clusterProfiler* (v 4.18.4), *msigdbr* (v26.1.0), *dplyr* (v1.2.1), *ggplot2* (v4.0.3).

##### Assessment of pathway coverage by the Olink proteomic platform

To evaluate how comprehensively biological pathways were represented by the Olink Explore 3072 proteomic platform, pathway coverage was calculated independently for all Hallmark, KEGG Legacy, and Reactome gene sets retrieved from the Molecular Signatures Database (MSigDB) using the *msigdbr* package (v 26.1.0).

First, all protein identifiers measured by the Olink platform were harmonized by removing panel-specific suffixes (e.g., Inflammation, Oncology, Cardiometabolic, and Neurology) to obtain unique gene symbols. For each pathway, the total number of annotated genes were extracted from MSigDB and compared with the number of pathway genes represented among Olink-measured proteins.

Pathway coverage was calculated as the ratio between the number of pathway genes (i.e., proteins) quantified by the Olink platform and the total number of genes defining the corresponding pathway in MSigDB.

Coverage estimates were subsequently integrated with the over-representation analysis (ORA) results obtained from the common IL6-associated protein signature. For each significantly enriched pathway, two complementary metrics were displayed: (i) the protein ratio (GeneRatio), corresponding to the proportion of input proteins assigned to the pathway by ORA, and (ii) the pathway coverage, reflecting the proportion of the entire biological pathway represented by the Olink proteomic panel.

Visualization consisted of paired plots. Dot plots displayed the GeneRatio, with point size proportional to the number of proteins contributing to pathway enrichment. Adjacent horizontal bar plots represented pathway coverage, with bar color indicating the number of pathway proteins detected by the Olink platform. A dashed horizontal reference line was drawn at 20% pathway coverage to facilitate interpretation of pathway representation across the proteomic panel. For subsequent analyses only pathways with a coverage above 20% and containing at least 10 proteins represented in the Olink proteomic dataset were retained.

R packages *clusterProfiler* (v 4.18.4), *msigdbr* (v26.1.0), *dplyr* (v1.2.1), *ggplot2* (v4.0.3).

##### Calculation of pathway activity scores

To quantify pathway activity at the individual sample level, single-sample Gene Set Enrichment Analysis (ssGSEA) was performed using the *GSVA* package (v2.4.9) in R. Gene sets were obtained from the Molecular Signatures Database (MSigDB) using the *msigdbr* package (v26.1.0) and included the Hallmark (H), KEGG Legacy (C2:CP:KEGG_LEGACY), and Reactome (C2:CP:REACTOME) collections. Only pathways previously identified as significantly enriched and meeting the predefined coverage criteria (>20% pathway coverage and ≥10 proteins represented by the Olink proteomic panel) were retained for score calculation. Protein expression matrices were generated from Olink Explore 3072 normalized protein expression (NPX) values. Proteins with more than 50% missing values across samples were excluded. Remaining missing values were imputed using the median NPX value of the corresponding protein across all samples. Sample-specific pathway activity scores were then computed using the ssGSEA algorithm implemented in the *GSVA* package, generating one enrichment score per pathway and per individual. Higher scores indicate greater coordinated abundance of pathway-associated proteins within a given sample.

R packages *GSVA* (v 2.4.9), *msigdbr* (v26.1.0), *dplyr* (v1.2.1), *ggplot2* (v4.0.3).

##### Association between pathway activity scores and cancer-free survival

Associations between pathway activity scores and cancer-free survival were assessed in the pooled FLEMENGHO and PREVALUNG cohorts. For each pathway, continuous ssGSEA activity scores were evaluated as independent variables in univariable Cox proportional hazards regression models. Cancer-free survival was defined as the time from cohort inclusion to cancer diagnosis or last follow-up, with cancer diagnosis considered as the event. For each Cox model, hazard ratios (HRs), 95% confidence intervals (CIs), and *P* values were extracted. HRs were displayed on a logarithmic scale, with pathways associated with increased cancer incidence (HR > 1) highlighted in pink and pathways associated with decreased cancer incidence (HR < 1) shown in black.

R packages *dplyr* (v1.2.1), *ggplot2* (v4.0.3), and *survival* (v3.8-6)

#### S-score and TOPOSCORE calculation

##### Metagenomic analysis of patient stools

Filtered high-quality reads were mapped with an identity threshold of 95% to the gene catalogs using Bowtie2 v2.3.4 88 included in METEOR version 3.2 (https://github.com/metagenopolis/meteor) software, and then a gene abundance profiling table was generated by means of a two-step procedure. First, reads mapped to a unique gene in the catalog were attributed to their corresponding genes. Second, reads that mapped with the same alignment score to multiple genes in the catalog were attributed according to the ratio of their unique mapping counts. Microbial genomes were recapitulated in >4,000 species-level genome bins (SGBs), and the gene abundance table was processed using MetaPhlAn4 package.^100^ S-score and TOPOSCORE were calculated from gut microbiota metagenomic profiles according to the methodology described by *Derosa et al*.^52^

#### Principal component analysis of plasma proteomic profiles

To evaluate whether rule-out score-defined cancer risk groups exhibited distinct global plasma proteomic profiles, unsupervised principal component analyses (PCA) were performed independently in the FLEMENGHO and PREVALUNG cohorts using Olink Explore 3072 proteomic data.

Proteomic features corresponding to Olink protein abundance values (NPX) were used as input variables. Proteins with more than 50% missing values across samples were excluded. PCA was performed using the *PCA()* function implemented in the *FactoMineR* package (v2.16).

Samples were visualized according to the first two principal components and colored by rule-out score categories (Rule-out versus At risk). Group separation was further evaluated using permutational multivariate analysis of variance (PERMANOVA) based on Euclidean distances calculated from scaled proteomic data, with 999 permutations. The pseudo-F statistic (F), proportion of variance explained (R²) and significance level (*P* value) were reported.

The contribution of individual proteins to principal components was calculated from PCA loading contributions. Proteins contributing above the theoretical contribution threshold (100 divided by the total number of total proteins) were considered major contributors to the corresponding principal component.

##### Identification of PCA-contributive proteins shared between FLEMENGHO and PREVALUNG cohorts

To identify proteins contributing to the proteomic separation associated with rule-out score categories, PCA-contributive proteins were determined independently in the FLEMENGHO and PREVALUNG cohorts. For each cohort, proteins were considered major contributors when their contribution to principal component 1 and/or principal component 2 exceeded the theoretical contribution threshold. Only principal components showing a significant difference between rule-out score categories, as assessed by Welch’s t-test, were considered for downstream protein selection (*i.e*., Dim 1 and 2 in FLEMENGHO, and Dim 1 in PREVALUNG). For each cohort, proteins were further filtered based on differential level between rule-out score categories using Kruskal–Wallis testing with Benjamini–Hochberg false discovery rate (FDR) correction, retaining proteins with FDR < 0.05. The overlap between FLEMENGHO and PREVALUNG contributive protein lists was assessed using Venn diagram analysis (n = 523 proteins; refer to **Supplementary Table S5**).

To identify biological pathways consistently associated with the selected proteins, over-representation analyses (ORA), assessment of pathway coverage by the Olink proteomic platform, and calculation of pathway activity scores were performed as described above. Associations between pathway activity scores and cancer-free survival were first assessed independently in the FLEMENGHO and PREVALUNG cohorts using univariable Cox proportional hazards regression models. Only pathways significantly associated with cancer-free survival in both cohorts (*P* value < 0.05 in each cohort) were retained for subsequent analysis. These pathways were then evaluated in the pooled FLEMENGHO and PREVALUNG cohorts using univariable Cox proportional hazards regression models.

R packages *FactoMineR* (v2.16), *factoextra* (v2.1.0), *dplyr* (v1.2.1), *ggplot2* (v4.0.3), *ggExtra* (v0.11.0), *vegan* (v 2.7-5), *ggvenn* (v0.1.19), *survival* (v3.8-6), and *survminer* (v0.5.2)

#### Cell-type marker-based pathway analysis and activity score calculation

Human cell-type marker sets were obtained from the <u>CellMarker 2.0 database</u> (*Human cell markers; Cell markers of different cell types from different tissues in human database*), which provides curated cell-type markers across different human tissues. Marker genes with an available UniProt identifier were retained for subsequent analyses. Cell-type marker sets were defined by combining tissue class, tissue type, cell type (Normal or Cancer cells), and cell name to generate unique cell-type pathway identifiers.

The coverage of CellMarker 2.0 cell-type marker sets by the Olink proteomic platform was assessed by calculating, for each cell-type marker set, the proportion of marker genes quantified in the Olink panel relative to the total number of genes defining the corresponding cell-type marker set. Cell-type marker sets with >20% coverage and at least 10 detected proteins were retained for subsequent activity score calculation.

Cell-type pathway activity scores were calculated using single-sample gene set enrichment analysis (ssGSEA). Olink NPX data were used as the expression matrix, with proteins detected in at least 50% of participants retained. Missing values were imputed using the median value of the corresponding protein across participants. For each retained cell-type marker set, only genes quantified by the Olink platform were included in the gene set. Cell-type activity scores were then calculated using the GSVA R package (v2.4.9) with the ssGSEA method. Resulting scores were obtained for each participant and each retained cell-type marker set and used for subsequent statistical analyses.

R packages *GSVA* (v 2.4.9), *dplyr* (v1.2.1), and *ggplot2* (v4.0.3).

#### Actionable protein selection and unsupervised clustering

Proteins identified as consistently associated with rule-out score stratification across the FLEMENGHO and PREVALUNG cohorts were further evaluated for their potential clinical relevance. Among the 523 proteins identified from the overlap of cohort-specific PCA-contributive proteins, proteins annotated as clinical-stage targets or as having strong lung-cancer biological relevance but not yet clinically validated as lung-cancer targets were selected for downstream analysis.

For the selected proteins, plasma protein abundance data from the FLEMENGHO and PREVALUNG cohorts were combined and subjected to unsupervised hierarchical clustering. Protein abundance values (NPX) were organized into a participant-by-protein matrix and clustered using Euclidean distance and Ward’s minimum-variance method (Ward.D2). The resulting hierarchical tree of patients was cut into two clusters to identify distinct molecular profiles based on plasma protein abundance. Cluster assignments were subsequently integrated with clinical and survival data.

##### Survival analysis according to molecular clusters

Kaplan–Meier survival curves were generated to compare cancer-free survival between the two clusters, and differences in survival distributions were assessed using the two-sided log-rank test. The associations between individual proteins characterizing the poor-prognosis cluster and cancer-free survival were further evaluated using univariable Cox proportional hazards regression models. Hazard ratios (HRs) and corresponding 95% confidence intervals (CIs) were estimated for Cluster 1 versus Cluster 2. Statistical analyses were performed in the pooled FLEMENGHO and PREVALUNG cohorts.

R packages *dplyr* (v1.2.1), *ggplot2* (v4.0.3), *pheatmap* (v1.0.13), *scales* (v1.4.0), *cowplot* (v 1.2.0), *survival* (v3.8-6), and *survminer* (v0.5.2)

#### Pathway over-representation analysis associated with circulating CTRC and MSMB

Spearman correlation analyses were performed between plasma CTRC or MSMB protein levels and all other proteins quantified by the Olink Explore 3072 platform. Proteins consistently correlated with CTRC or MSMB across the FLEMENGHO and PREVALUNG cohorts were selected. The final protein panels corresponded to the intersection of proteins significantly correlated with CTRC or MSMB in both cohorts after Benjamini-Hochberg false discovery rate (FDR) correction (FDR < 0.05) and showing an absolute Spearman correlation coefficient (rho) > 0.3.

To identify biological pathways consistently associated with the selected proteins, over-representation analyses (ORA), assessment of pathway coverage by the Olink proteomic platform, and calculation of pathway activity scores were performed as described previously. In addition, to identify cell type signatures consistently associated with the selected proteins (rho >0.3), ORA were performed using gene sets from the cell type signature (C8) collection of MSigDB. The C8 collection comprises gene sets representing cell type-specific transcriptional signatures and was used to characterize the cellular signatures associated with the selected proteins.

Associations between pathway activity scores and cancer-free survival were first assessed independently in the FLEMENGHO and PREVALUNG cohorts using univariable Cox proportional hazards regression models. Only pathways significantly associated with cancer-free survival in both cohorts (*P* value < 0.05 in each cohort) were retained for subsequent analysis. These pathways were then evaluated in the pooled FLEMENGHO and PREVALUNG cohorts using univariable Cox proportional hazards regression models.

R packages *FactoMineR* (v2.16), *factoextra* (v2.1.0), *dplyr* (v1.2.1), *ggplot2* (v4.0.3), *ggExtra* (v0.11.0), *vegan* (v 2.7-5), *ggvenn* (v0.1.19), *survival* (v3.8-6), and *survminer* (v0.5.2)

#### Tissue expression analysis

##### Transcriptomic tissue expression analysis using GTEx

To characterize the tissue distribution of genes encoding proteins of interest, bulk RNA expression data from the Genotype-Tissue Expression (GTEx) project (<u>GTEx Analysis Release v10</u>; *Median gene-level TPM by tissue. Median expression was calculated from the file GTEx_Analysis_v10_RNASeQCv2.4.2_gene_tpm.gct.gz*) were analyzed. Gene-level median expression values across normal human tissues were retrieved from the GTEx RNA-sequencing dataset and expressed as transcripts per million (TPM). Tissues were grouped into broader anatomical categories, including adipose tissue, arteries, bladder, breast, brain, colon, esophagus, heart, kidney, liver, lung, pancreas, skin, skeletal muscle, small intestine, spleen, stomach, thyroid, and whole blood. Gender-specific tissues, including testis, ovary, prostate, uterus, and vagina, were excluded from the analysis. When multiple GTEx anatomical compartments were available for a given organ, the mean expression value across corresponding tissues was calculated. Genes of interest were selected based on the corresponding protein candidates identified in the plasma proteomic analyses (Olink dataset). Expression values were log2-transformed for visualization. Heatmap was generated to display the expression levels of selected genes across healthy human tissues.

##### Protein tissue expression analysis using the Human Protein Atlas

Protein-level tissue expression patterns were evaluated using immunohistochemistry-based protein expression data from the Human Protein Atlas (<u>HPA, version 19.3</u>; *normal_tissue.tsv.zip*). Proteins corresponding to selected proteins of interest were extracted based on the annotated gene symbols. Protein expression levels across normal human tissues were converted into ordinal values (Not detected = 0, Low = 1, Medium = 2, High = 3). When multiple annotations were available for a given gene and tissue pair, the highest expression level was retained. Protein expression patterns across normal tissues were visualized using an heatmap.

##### Integration of GTEx tissue enrichment and plasma proteomic associations

To investigate whether proteins circulating in plasma displayed tissue-specific expression patterns, GTEx-based tissue enrichment profiles were integrated with plasma proteomic correlation analyses. Genes corresponding to proteins quantified by the Olink Explore 3072 platform were annotated according to their tissue enrichment profile derived from GTEx RNA-sequencing data. Tissue enrichment was assessed by calculating the fold-change (FC) of gene expression in each tissue relative to the geometric mean expression across all other analyzed tissues. Genes were considered tissue-enriched when the maximal tissue-specific FC was ≥4. Gender-specific tissues (testis, ovary, prostate, uterus, and vagina) were excluded from this analysis. Plasma protein levels of CTRC or MSMB were correlated with all available Olink proteins in the pooled FLEMENGHO and PREVALUNG cohorts using Spearman correlation analyses. Multiple testing correction was performed using the Benjamini-Hochberg (BH) false discovery rate (FDR) method. For each tissue category, the proportion of tissue-enriched proteins significantly associated with CTRC or MSMB was calculated. Positive correlations were defined as proteins showing Spearman correlation coefficients rho > 0.3 and FDR < 0.05, whereas negative correlations were defined as rho < −0.1 and FDR < 0.05. Proteins with FDR < 0.05 and correlation coefficients between −0.1 and 0.1 were considered to display neutral associations. The percentage of significantly correlated proteins among tissue-enriched proteins was calculated separately for each tissue category and visualized using bar plots.

R packages *tidyr* (v1.3.2), *tibble* (v3.3.1), *dplyr* (v1.2.1), *ggplot2* (v4.0.3), and *stringr* (v1.6.0)

#### Differential plasma proteins levels according to rule-out or rule-in score categories

Participants from the FLEMENGHO and PREVALUNG cohorts were pooled and stratified according to rule-out score (Rule-out vs At risk) or the rule-in (Medium risk vs High risk) categories. Plasma protein abundance was assessed using the Olink Explore 3072 proteomic platform. Proteins with more than 50% missing values were excluded from the analysis. For each protein, differences in plasma abundance between rule-out or rule-in score categories were evaluated using two-sided Wilcoxon rank-sum tests. As Olink NPX values are reported on a log2 scale, the effect size was calculated as the difference between the median protein levels in the At risk and Rule-out groups, corresponding to the log2 fold-change (log2FC).

Multiple testing correction was performed using the Benjamini–Hochberg (BH) false discovery rate (FDR) procedure. Volcano plots display the log2FC on the x-axis and the −log_10_(FDR) on the y-axis. Proteins were considered significantly differentially abundant when FDR < 0.05 and the absolute log2FC was ≥0.5. Proteins of particular interest were annotated individually on the volcano plot.

R packages *dplyr* (v1.2.1), *stats* (v4.5.2), *ggplot2* (v4.0.3), and *ggrepel* (v0.9.8)

#### Evaluation of rule-out and rule-in scores by ROC and survival analyses

##### Performance evaluation of the risk scores

The discriminative performance of the rule-out and rule-in scores was assessed using receiver operating characteristic (ROC) curve analyses. ROC curves were generated independently in the FLEMENGHO and PREVALUNG cohorts using cancer status (Control versus Cancer) as the outcome. For the rule-out score, the predefined threshold of 26.13% was evaluated, whereas for the rule-in score, only individuals classified as “At risk” after the rule-out step (rule-out score ≥26.13%) were included, and the predefined rule-in threshold of 68.73% was assessed. For the combined score, individuals classified as rule-out were assigned a value of 0, while the continuous rule-in score was retained for individuals classified as at risk after the rule-out step. Sensitivity, specificity, positive predictive value (PPV), and negative predictive value (NPV) were calculated for each threshold. Areas under the ROC curves (AUCs) with 95% confidence intervals were estimated using the DeLong method.

##### Cancer-free survival analyses

Associations between risk-score categories and cancer-free survival were evaluated using Kaplan-Meier survival analyses. Time-to-event was defined as the interval between baseline assessment and cancer diagnosis or last follow-up, and survival distributions were compared using the log-rank test. Hazard ratios (HRs) and 95% confidence intervals (CIs) were estimated using univariable Cox proportional hazards regression models. For the combined two-stage risk stratification, participants were classified into three groups: rule-out (rule-out score <26.13%), medium risk (rule-out score ≥26.13% and rule-in score <68.73%), and high risk (rule-out score ≥26.13% and rule-in score ≥68.73%). Pairwise Cox regression models and pairwise log-rank tests were performed to compare risk groups, with P values adjusted using the Benjamini–Hochberg method.

##### Performance evaluation of the PLCO_m2012_ risk score and eligibility criteria from lung cancer screening trials (NELSON and NLST)

The modified 2011 lung cancer risk prediction model derived from the Prostate, Lung, Colorectal, and Ovarian (PLCO) Cancer Screening Trial (PLCO_m2012_), which estimates the 6-year probability of developing lung cancer,^3^ was calculated for participants from the FLEMENGHO and PREVALUNG cohorts.

For each participant a linear predictor (logit) was computed as a weighted sum of the model’s covariates: age (centered at 62), race/ethnicity, education level, body-mass index (centered at 27), self-reported chronic obstructive pulmonary disease (COPD), personal history of cancer, family history of lung cancer, smoking status (current vs. former), smoking intensity, smoking duration (years, centered at 27) and time since smoking cessation (years, centered at 10), each multiplied by its published PLCO β-coefficient and added to the model intercept. PLCO_m2012_ score was obtained from the logistic link function and expressed as a percentage: *PLCO (%) = exp(logit) / [1 + exp(logit)] × 100*.^3^

For the NELSON trial, eligibility criteria included an age of 50-75 years and a smoking history of either more than 15 cigarettes per day for more than 25 years or more than 10 cigarettes per day for more than 30 years, with participants being either current smokers or having stopped smoking less than 10 years before inclusion. In the NLST, eligibility criteria included an age of 55-74 years and a cumulative smoking exposure of at least 30 pack-years, with participants being either current smokers or having quit smoking within the previous 15 years. Participants meeting all eligibility criteria were classified as “Yes” for the corresponding NELSON or NLST criteria; all other participants were classified as “No”.

ROC curves were generated independently in the FLEMENGHO and PREVALUNG cohorts using cancer status (Control versus Cancer) as the outcome. Sensitivity, specificity, PPV, and NPV were calculated at the predefined PLCO_m2012_ risk threshold of 1.5%^3^ in both cohorts and according to NELSON and NLST eligibility status (Yes versus No) in the PREVALUNG cohort. AUCs and corresponding 95% confidence intervals (CIs) were estimated using the DeLong method. Cancer-free survival was evaluated according to PLCO_M2012_ risk categories (<1.5% versus ≥1.5%) and NELSON and NLST eligibility status (Yes versus No) using Kaplan-Meier survival analyses.

##### Statistical analyses and visualization

ROC curves were generated using the *pROC* (v1.19.0.1) package. Kaplan-Meier curves and Cox proportional hazards regression analyses were performed using the *survival* (v3.8-6) and *survminer* (v0.5.2) packages.

R packages *dplyr* (v1.2.1), *ggplot2* (v4.0.3), *pROC* (v1.19.0.1), *survival* (v3.8-6), and *survminer* (v0.5.2)

## Legends to extended figures

**Extended Data Fig. 1.**
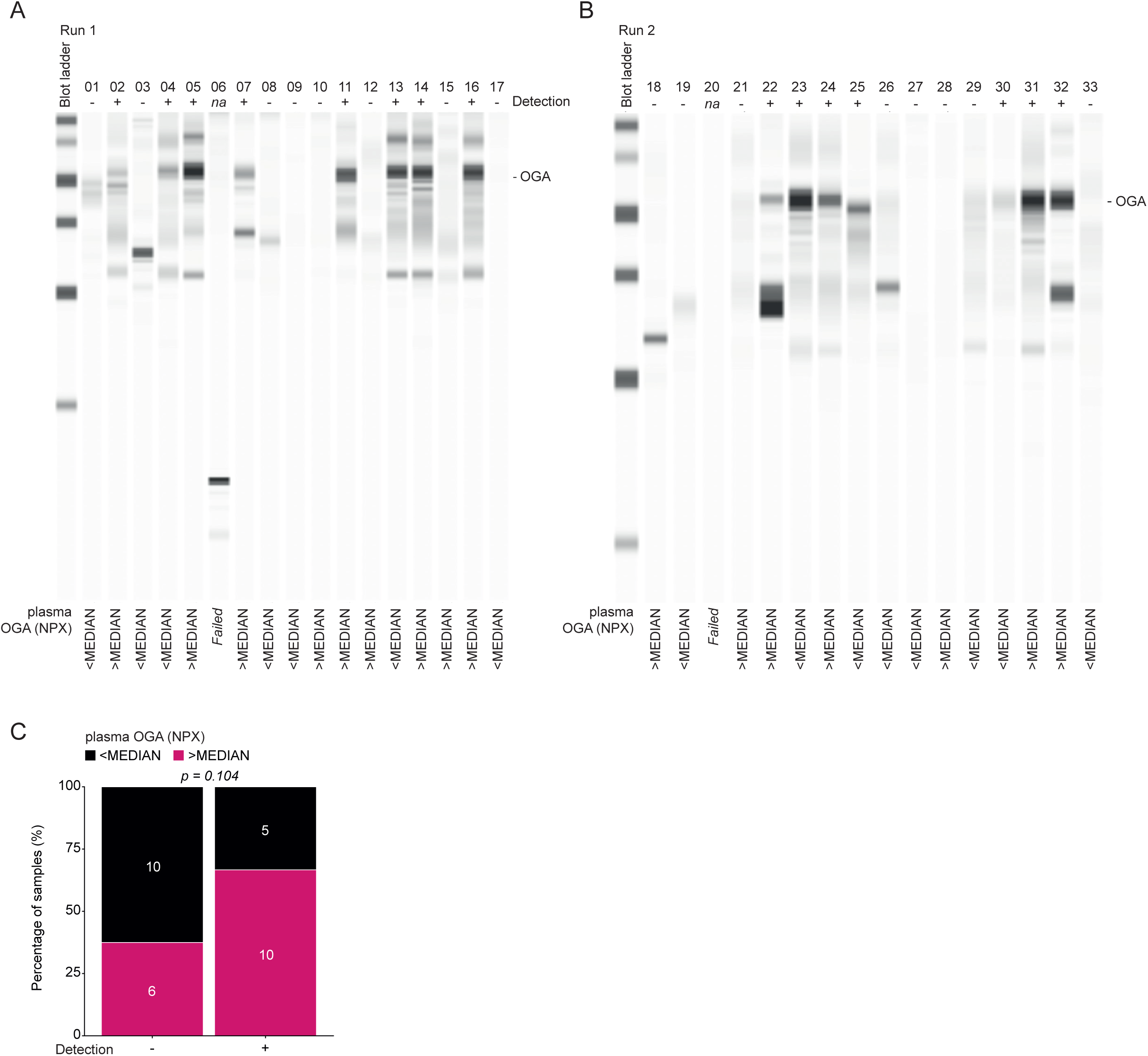
Orthogonal validation of OGA protein expression in circulating immune cells by automated Western blotting. **a-b**, Representative electropherograms generated using the Jess Automated Western Blot System (Jess) for detection of OGA protein in frozen PBMC dry pellets from two independent analytical runs: run 1 (a, n = 17) and run 2 (b, n = 16). **c**, Contingency analysis comparing OGA detection by Jess (detected versus not detected) with circulating serum OGA levels dichotomized according to the median value (< versus > the median in the cohort). The association between OGA detection in PBMCs and serum OGA category was assessed using a contingency-table analysis. Overall, N = 35 participants from the PREVALUNG cohort were included across the two analytical runs. The stacked bar chart shows the proportion of samples classified as <MEDIAN (black) or >MEDIAN (pink) within Jess detection category (detected versus not detected). Bars represent the percentage of samples in each category, and the absolute number is displayed within each segment. The association was assessed using Pearson’s chi-squared test. The corresponding two-sided *P* value is shown in the figure.

**Extended Data Fig. 2.**
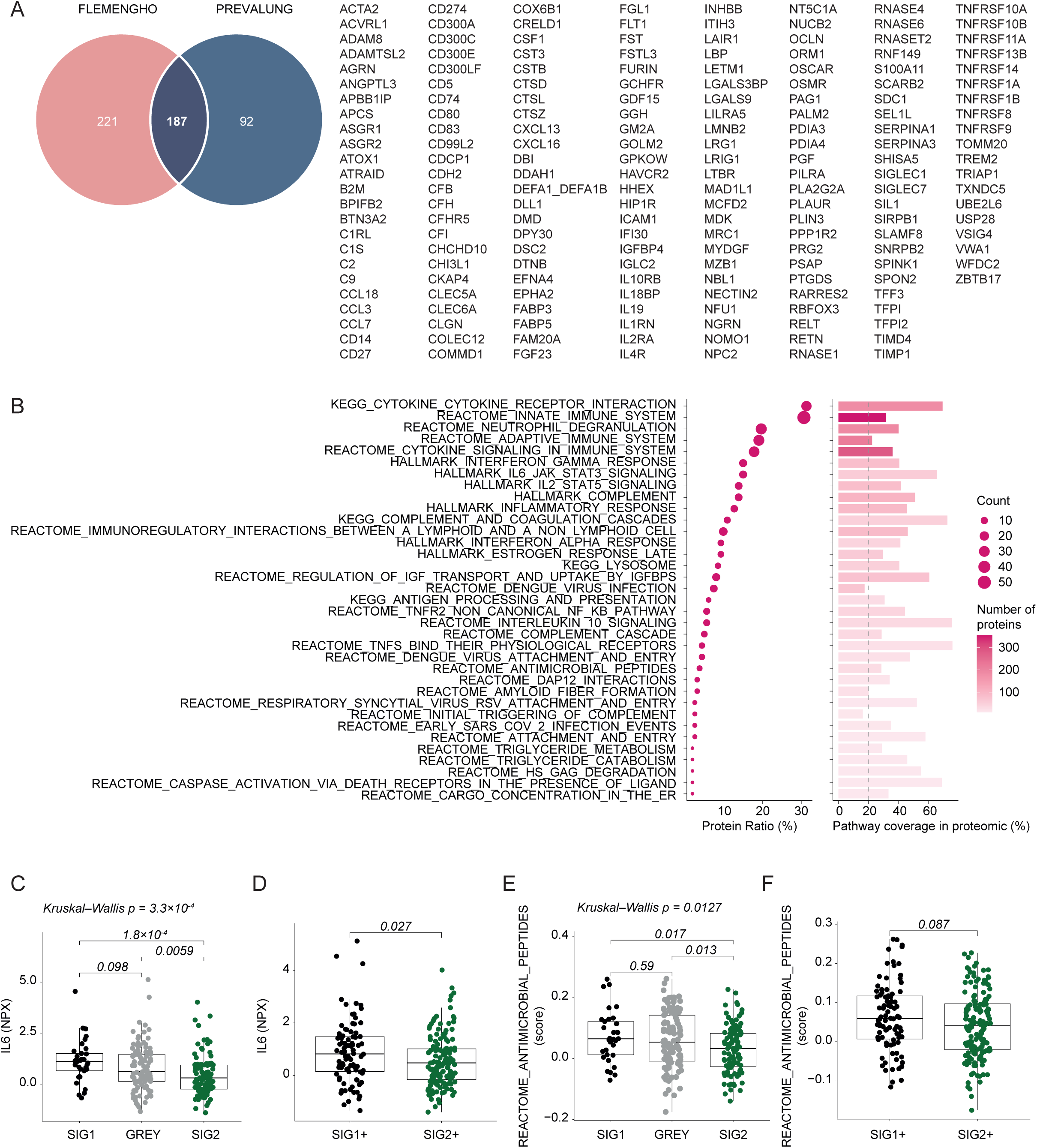
Biological function analysis associated with circulating IL6. **a**, Venn diagram representing the intersection of proteins significantly correlated with IL6 after Benjamini–Hochberg false discovery rate (FDR) correction and with absolute rho value >0.3 in both cohorts across the FLEMENGHO (N = 154) and PREVALUNG (N = 396) cohorts (n = 187 proteins; refer to Fig. 3g-h). **b**, Pathway enrichment and coverage analysis were performed on the 187 proteins consistently correlated with plasma IL6 in the FLEMENGHO and PREVALUNG cohorts. Hallmark, KEGG Legacy, and Reactome pathways enriched by over-representation analysis (ORA) are shown. The left panel displays the proportion of IL6-associated proteins assigned to each pathway (Protein Ratio (%)), with dot size indicating the number of contributing proteins (Count). The right panel shows pathway coverage within the Olink Explore 3072 proteomic platform, calculated as the percentage of proteins defining each pathway in MSigDB that were quantified by Olink. Bar color indicates the number of detected pathway proteins. The dashed line indicates the 20% pathway coverage threshold used for pathway selection in downstream analyses. Only pathways with *P* value < 0.05 are displayed. **c-f**, Participants from the PREVALUNG cohort (N = 241) were stratified according to S-score categories (SIG1, Grey, and SIG2) (C and E) or TOPOSCORE categories (SIG1+ and SIG2+) (D and F). Plasma IL6 levels (**c-d**) and REACTOME_ANTIMICROBIAL_PEPTIDES pathway activity scores derived from ssGSEA (**e-f**) were compared between groups. Boxplots display the median and interquartile range, while each dot represents an individual participant. Group differences were assessed using Kruskal-Wallis (KW) tests followed by pairwise two-sided Wilcoxon rank-sum tests with Benjamini-Hochberg (BH) correction for multiple comparisons, or two-sided Wilcoxon rank-sum tests for two-group comparisons. KW *P* values and BH-adjusted *P* values, or nominal *P* values for two-group comparisons are shown.

**Extended Data Fig. 3.**
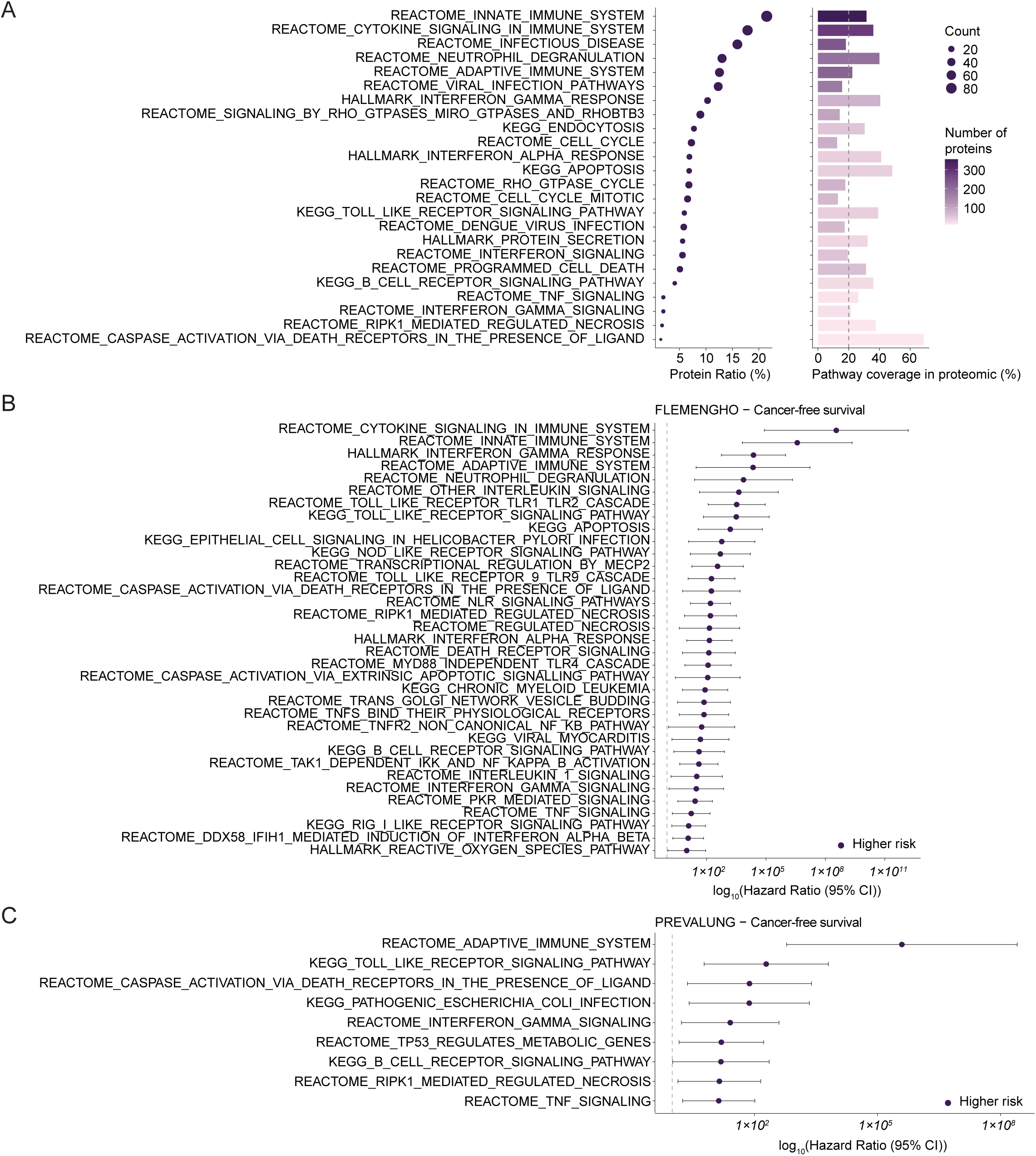
Identification of biological pathways associated with cancer risk stratification by the rule-out score. **a**, Pathway enrichment and coverage analysis were performed on the 523 proteins consistently associated with rule-out score stratification across both FLEMENGHO (N = 154) and in PREVALUNG (N = 396) independent cohorts (refer to Fig. 4a-b). Hallmark, KEGG Legacy, and Reactome pathways enriched by over-representation analysis (ORA) are shown. The left panel displays the proportion of proteins assigned to each pathway (Protein Ratio (%)), with dot size indicating the number of contributing proteins (Count). The right panel shows pathway coverage within the Olink Explore 3072 proteomic dataset, calculated as the percentage of proteins defining each pathway in MSigDB that were quantified by Olink. Bar color indicates the number of detected pathway proteins. The dashed line indicates the 20% pathway coverage threshold used for pathway selection in downstream analyses. Displayed pathways have a *P* value < 0.05. **b-c**, Forest plots display univariable Cox proportional hazards regression analyses assessing the associations between continuous pathway activity scores and cancer-free survival in the FLEMENGHO (N = 154) (**b**) and PREVALUNG (N = 396) (**c**) cohorts (*P* value < 0.05). Hazard ratios (HRs) and 95% confidence intervals (CIs) are shown on a log10 scale. The dashed line indicates HR = 1.

**Extended Data Fig. 4.**
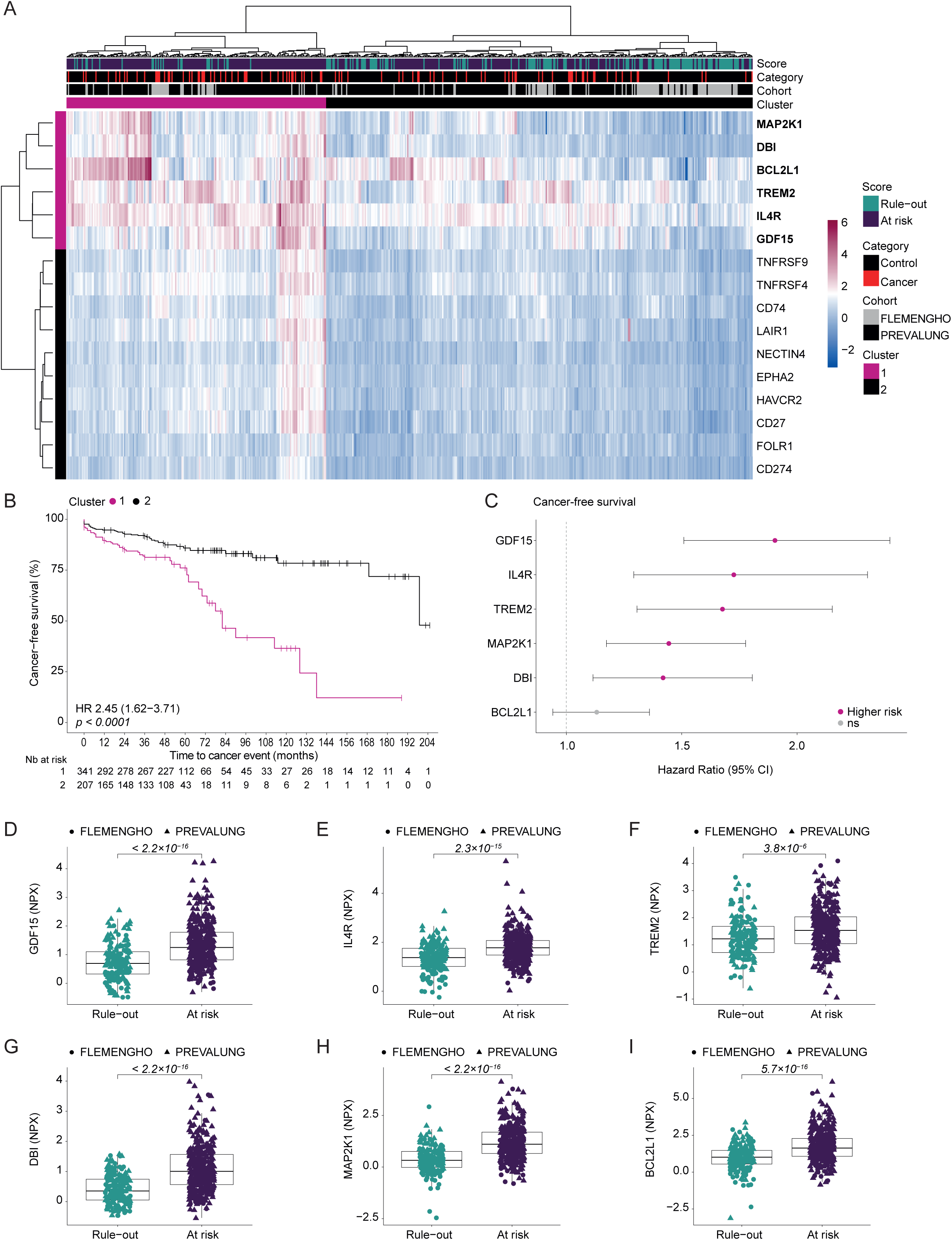
Identification and prognostic characterization of molecular clusters based on actionable plasma proteins. **a**, Heatmap showing the unsupervised hierarchical clustering of participants from the combined FLEMENGHO (N = 154) and in PREVALUNG (N = 396) cohorts based on the plasma levels of selected actionable proteins. Columns represent individual participants and rows represent proteins. Participants and proteins were hierarchically clustered using Euclidean distance and Ward.D2 linkage, and the resulting patient tree was cut into two clusters (k = 2). Clinical and rule-out score annotations are shown above the heatmap. **b**, Kaplan-Meier analysis of cancer-free survival according to the two clusters. The cancer-free survival probability is shown over follow-up, with the number of participants at risk indicated below the plot. The hazard ratio (HR) and 95% confidence interval (CI) for Cluster 1 versus Cluster 2, as well as the global log-rank *P* value, are indicated. **c**, Univariable Cox proportional hazards regression analyses of individual proteins characterizing the poor-prognosis cluster (Cluster 1) in relation to cancer-free survival. Hazard ratios and 95% CIs are shown for a continuous increase in protein level. Pink dots indicate statistically significant associations (*P* < 0.05), whereas non-significant associations (ns) are shown in grey. The vertical dashed line indicates HR = 1. **d**, Participants were stratified according to their rule-out score classification (Rule-out or At risk) and levels of the proteins characterizing the poor-prognosis cluster (Cluster 1) were compared between groups. Boxplots display the median and interquartile range, while each dot represents individual participants. Each individual from the FLEMENGHO (N = 154) and in PREVALUNG (N = 396) cohorts is represented by solid circles and by triangles, respectively. Difference between groups is assessed using two-sided Wilcoxon rank-sum tests, and the corresponding *P* value is shown on the figure.

**Extended Data Fig. 5.**
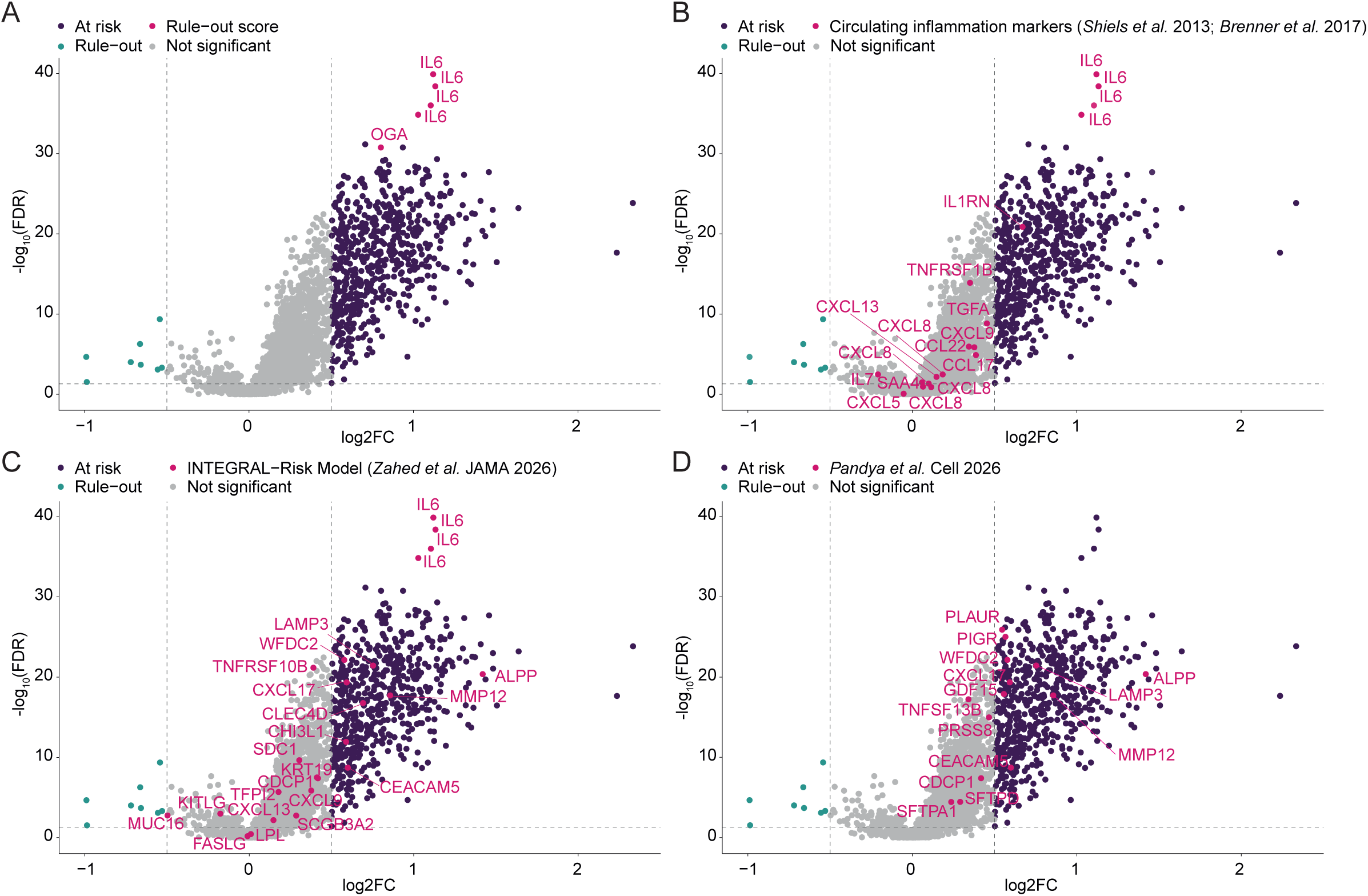
Differential plasma proteins levels according to rule-out score categories. **a-d**, Volcano plot of plasma protein levels according to rule-out score categories in the combined FLEMENGHO (N = 156) and PREVALUNG (N = 397) cohorts. Participants were stratified according to rule-out score categories (Rule-out vs At risk), and differential plasma protein levels was assessed using two-sided Wilcoxon rank-sum tests. The x-axis represents the log2 fold-change (log2FC) protein values (NPX) between At risk and Rule-out participants, whereas the y-axis represents −log_10_(FDR) of the Benjamini–Hochberg adjusted *P* value. Proteins with FDR < 0.05 and |log2FC| ≥ 0.5 were considered significantly differentially abundant. Proteins enriched in At risk participants are shown in purple, whereas proteins enriched in Rule-out participants are shown in blue. Non-significant proteins are displayed in grey. Dashed horizontal and vertical lines indicate the significance and fold-change thresholds, respectively. Proteins included in the rule-out score signature (**a**), or in circulating inflammation markers (*Shiels et al*. 2013; *Brenner et al*. 2017)^8,9^ (**b**), or in the INTEGRAL−Risk Model (*Zahed et al*. JAMA 2026)^16^ (**c**), or in *Pandya et al.* Cell 2026^17^ (**d**) are highlighted in pink and labelled.

**Extended Data Fig. 6.**
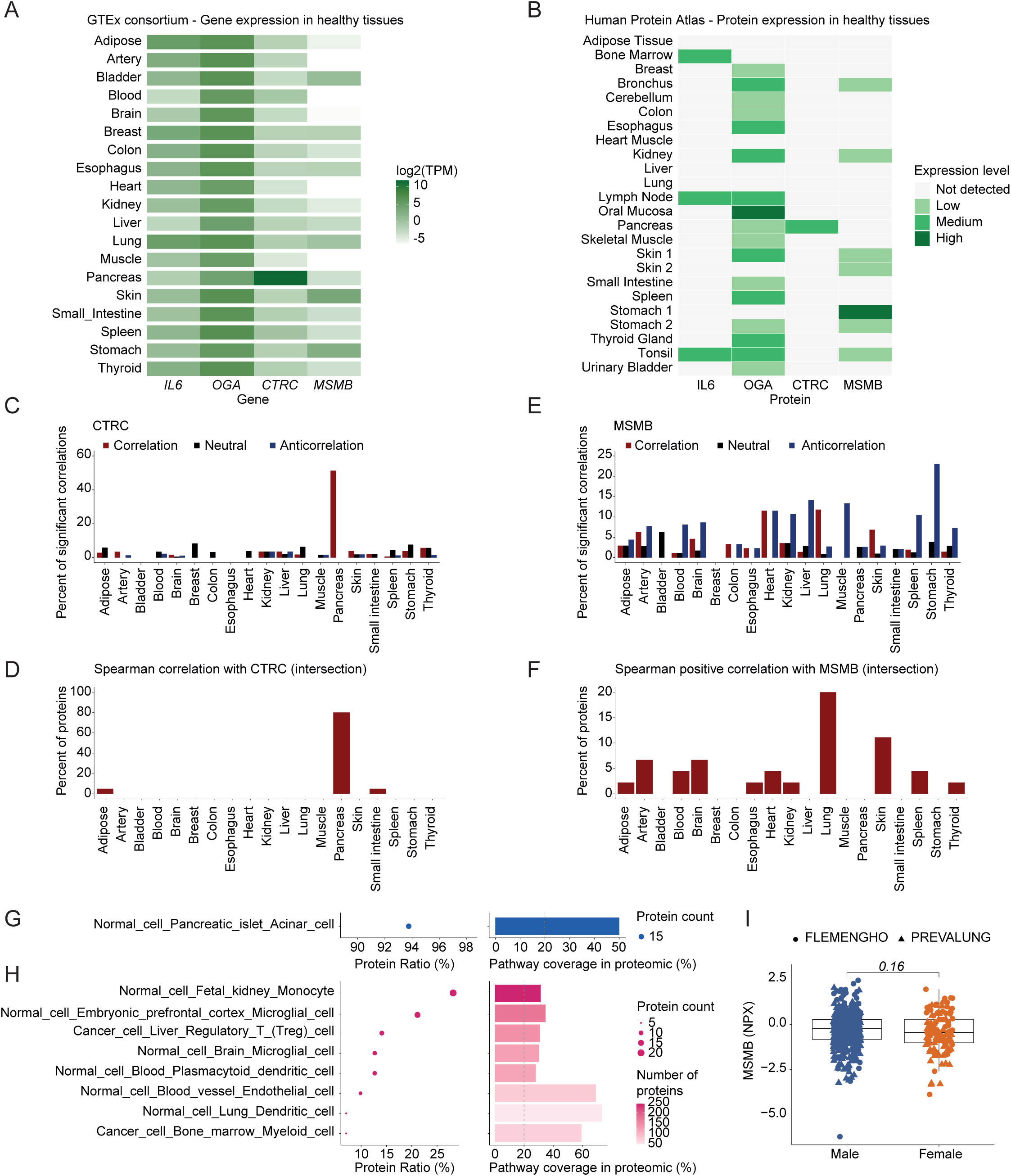
Tissue expression landscape of genes and proteins encoding circulating biomarkers. **a**, Heatmap showing the transcriptomic expression levels of genes encoding selected circulating biomarkers (IL6, OGA (MGEA5), CTRC, and MSMB) across healthy human tissues from the GTEx consortium. Expression values represent log2-transformed median TPM values. **b**, Heatmap showing protein expression levels of corresponding proteins across normal human tissues based on immunohistochemistry-derived annotations from the Human Protein Atlas (HPA). Expression categories were defined as not detected, low, medium, or high according to HPA criteria. **c-f**, Tissue enrichment profiles of genes corresponding to Olink-measured proteins were established using GTEx RNA-sequencing data. Genes were considered tissue-enriched when the maximal tissue-specific FC was ≥4. These tissue-specific gene expression annotations were integrated with plasma proteomic correlation analyses performed in the pooled FLEMENGHO (N = 154) and PREVALUNG (N = 396) cohorts. Protein correlations were assessed using Spearman correlation tests, and *P* values were adjusted using the Benjamini-Hochberg FDR method. For each tissue category, the proportion of tissue-enriched proteins showing significant associations with circulating CTRC (**c**) or MSMB (**e**) plasma levels was calculated. Distribution of tissue-enriched proteins according to their correlation patterns with circulating CTRC (**c**) or MSMB (**e**) levels in the pooled FLEMENGHO and PREVALUNG cohorts. The same analyses were subsequently performed using only the 20 proteins commonly correlated with CTRC (**d**) and the 92 proteins commonly correlated with MSMB (**f**) in the FLEMENGHO and PREVALUNG cohorts (Refer to Fig. 5b and g). Bar plots show the percentage of tissue-enriched proteins displaying positive correlations in red (Spearman rho > 0.3, FDR < 0.05), neutral associations in black (−0.1 < rho < 0.1, FDR < 0.05), or anticorrelations in bleu (rho < −0.1, FDR < 0.05). **g-h**, Cell type signatures enrichment and coverage analysis were performed on the 20 proteins commonly correlated with CTRC (**g**) and the 92 proteins commonly correlated with MSMB (**h**) in the FLEMENGHO and PREVALUNG cohorts (Refer to Fig. 5b and g). Cell type signatures from the C8 collection of MSigDB enriched by over-representation analysis (ORA) are shown. The left panel displays the proportion of CTRC or MSMB-associated proteins assigned to each cell type signature (Protein Ratio (%)), with dot size indicating the number of contributing proteins (Count). The right panel shows signature coverage within the Olink proteomic platform, calculated as the percentage of proteins defining each signature in MSigDB that were quantified by Olink. Bar color indicates the number of detected proteins contributing to each signature. The dashed line indicates the 20% pathway coverage threshold. Only cell type signatures with *P* value < 0.05 are displayed. **I**, Participants were stratified according to their gender (Male or Female) and MSMB levels (NPX) was compared between groups. Boxplots display the median and interquartile range, while each dot represents individual participants. Each individual from the FLEMENGHO (N = 156) and PREVALUNG (N = 397) cohorts is represented by solid circles and by triangles, respectively. Difference between groups is assessed using two-sided Wilcoxon rank-sum tests, and the corresponding *P* value is shown on the figure.

**Extended Data Fig. 7.**
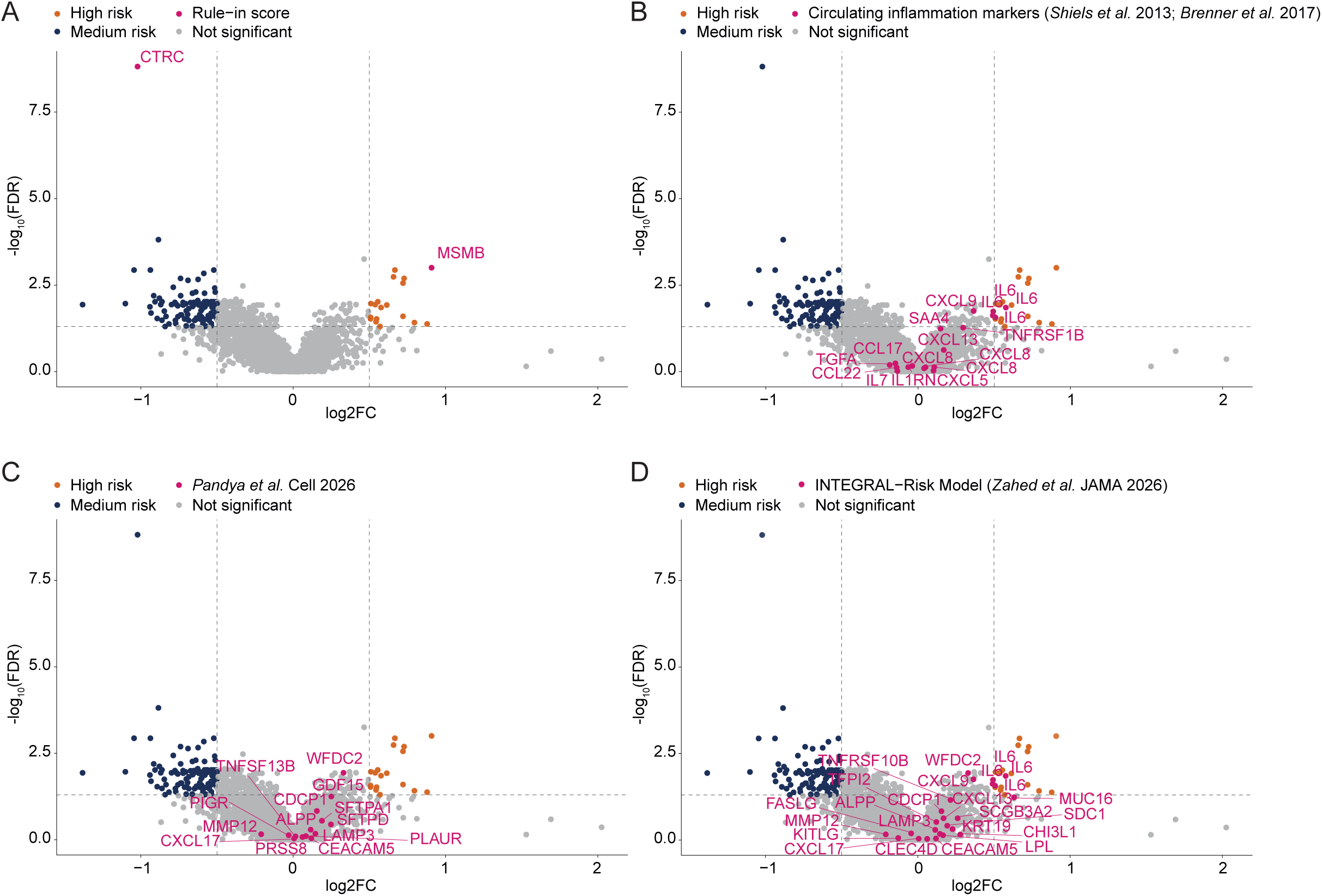
Differential plasma proteins levels according to rule-in score categories. **a-d**, Volcano plot of plasma protein levels according to rule-in score categories in the combined FLEMENGHO (N = 61) and PREVALUNG (N = 314) cohorts. Among participants classified as At risk after the rule-out step, individuals were stratified according to rule-in score categories (Medium risk vs High risk), and differential plasma protein levels was assessed using two-sided Wilcoxon rank-sum tests. The x-axis represents the log2 fold-change (log2FC) protein values (NPX) between At risk and Rule-in participants, whereas the y-axis represents −log_10_(FDR) of the Benjamini–Hochberg adjusted *P* value. Proteins with FDR < 0.05 and |log2FC| ≥ 0.5 were considered significantly differentially abundant. Proteins enriched in High risk participants are shown in orange, whereas proteins enriched in Medium risk participants are shown in blue. Non-significant proteins are displayed in grey. Dashed horizontal and vertical lines indicate the significance and fold-change thresholds, respectively. Proteins included in the rule-out score signature (**a**), or in circulating inflammation markers (*Shiels et al*. 2013; *Brenner et al*. 2017) (**b**), or in *Pandya et al.* Cell 2026 (**c**), or in the INTEGRAL−Risk Model (*Zahed et al*. JAMA 2026) (**d**) are highlighted in pink and labelled.

**Extended Data Fig. 8.**
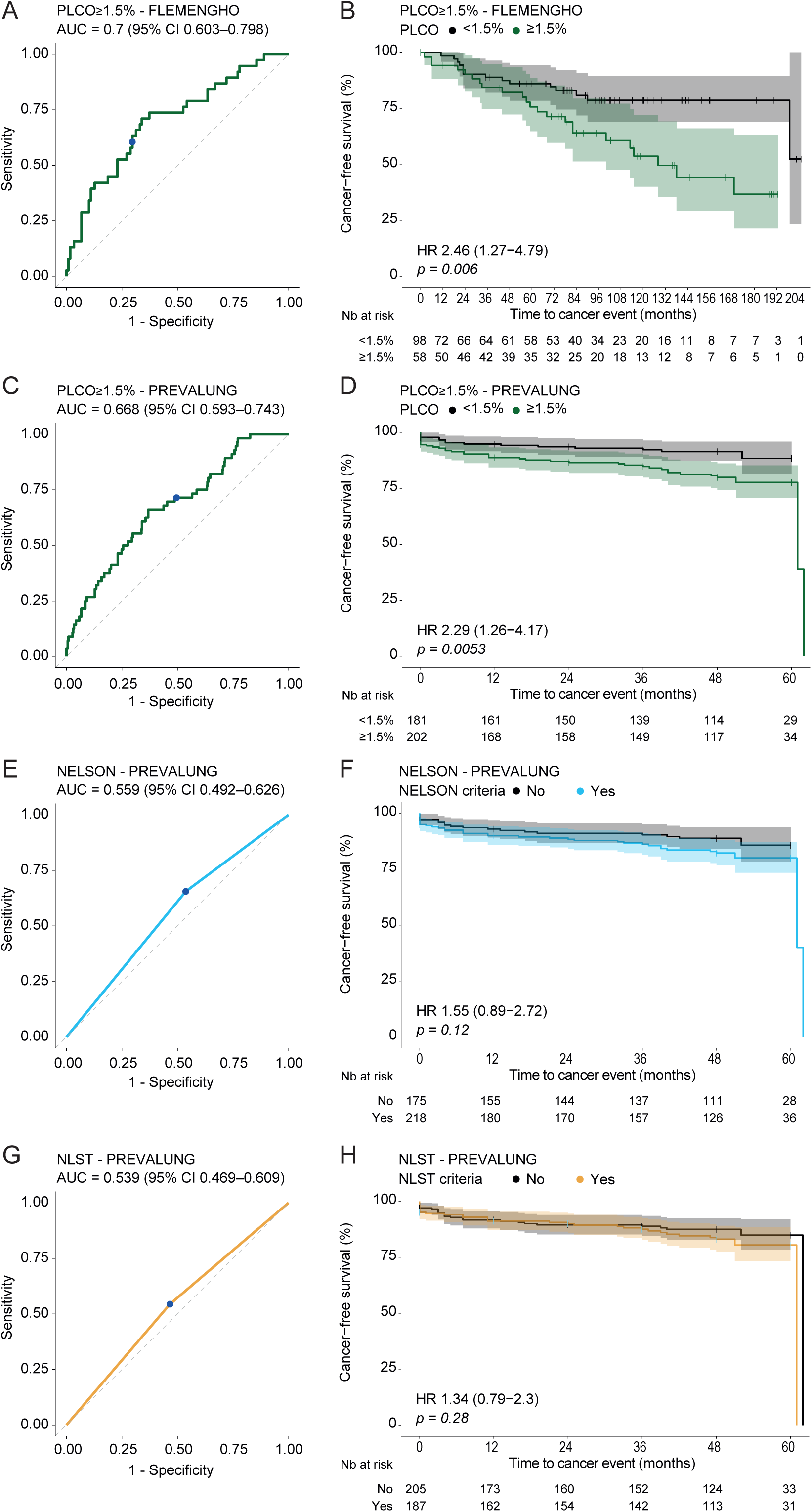
Performance evaluation of the PLCO risk score and eligibility criteria from lung cancer screening trials (NELSON and NLST) **a-g**, Receiver operating characteristic (ROC) curve analyses evaluating the discrimination of cancer status (**a, c, e and g**) and Kaplan-Meier analyses of cancer-free survival (**b, d, g, and h**) in the FLEMENGHO (**a-b**) and PREVALUNG (**c-h**) cohorts. Performance of the PLCO score using the predefined threshold of 1.5% in the FLEMENGHO cohort (**a**) and in the PREVALUNG cohort (**c**) and corresponding Kaplan-Meier analyses of cancer-free in the FLEMENGHO cohort (N = 156) (**b**) and in the PREVALUNG cohort (N = 383) (**d**). Performance of the NELSON (**e**) and NLST (**g**) criteria classification and corresponding Kaplan-Meier analyses of cancer-free survival according to the NELSON (**f**) and NLST (**h**) criteria classification in the PREVALUNG cohort (N = 392 and N = 393, respectively). ROC curves display sensitivity versus 1-specificity, with the corresponding area under the curve (AUC) and 95% confidence interval (CI) indicated. The blue dot represents the predefined score threshold or category. Kaplan-Meier curves represent cancer-free survival probabilities over follow-up time, with the number of participants at risk displayed below each plot. Global log-rank test *P* values and hazard ratios (HRs) and 95% CIs estimated using univariable Cox proportional hazards regression models are shown.

**Extended Data Fig. 9.**
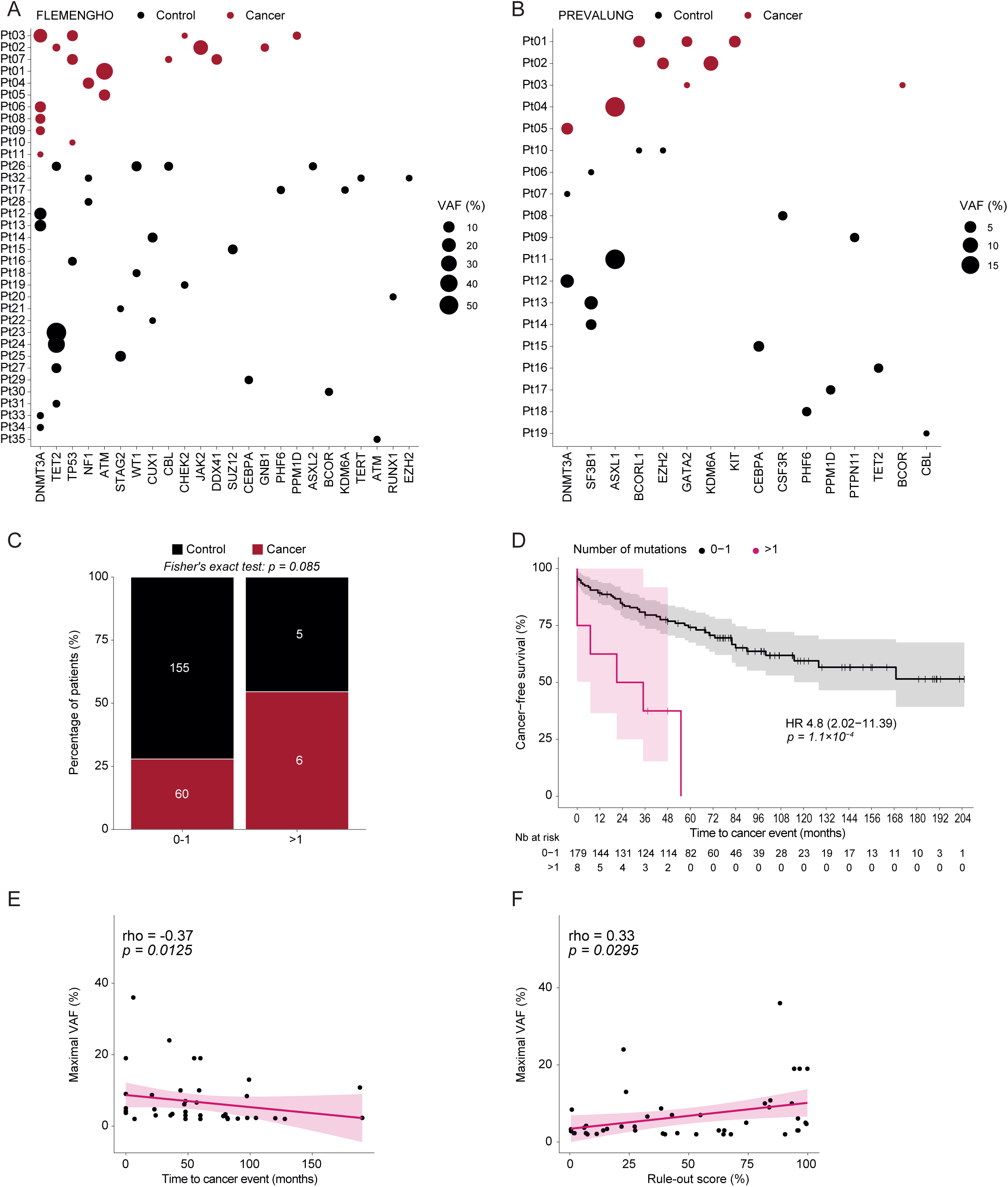
Characterization of clonal hematopoiesis of indeterminate potential in the FLEMENGHO and PREVALUNG cohorts. **a-b**, Distribution of detected clonal hematopoiesis of indeterminate potential (CHIP) mutations in the FLEMENGHO cohort (**a**, N = 35 patients with at least one mutation among 141 participants) and the PREVALUNG cohort (**b**, N = 19 patients with at least one mutation among 85 participants). Each dot represents a mutation detected in an individual patient. Patients are displayed on the y-axis, with cancer patients shown first and control participants below, and ordered within each group according to decreasing number of mutations. Genes are displayed on the x-axis. Dot size represents Variant Allele Frequency (VAF), and dot color indicates participant group (Cancer, red; Control, black). **c**, Distribution of participants with or without detected CHIP mutations in the combined FLEMENGHO and PREVALUNG cohorts (N = 226). Participants were classified into two categories according to the number of detected CHIP mutations: 0-1 or >1 mutation, and according to cancer status (Cancer vs. Control). Bars show the percentage of participants within each mutation category, with the number of participants indicated within each bar. Statistical comparison between Cancer and Control groups was performed using Fisher’s exact test, given the limited sample size. **d**, Kaplan-Meier analysis of cancer-free survival according to the number of detected CHIP mutations (0-1 versus >1 mutation) in the combined FLEMENGHO and PREVALUNG cohorts (N = 187). Shaded areas represent 95% confidence intervals, and tick marks indicate censored observations. The numbers of participants at risk at each time point are shown below. The hazard ratio (HR) with 95% confidence interval (CI) was estimated using a Cox proportional hazards model, with the 0-1 mutation group as the reference. Cancer-free survival was compared between groups using the two-sided log-rank test. **e-f**, Correlations between time to cancer diagnosis (**e**) or the rule-out score (**f**) and the maximal variant allele frequency (VAF) among participants carrying at least one CHIP mutation. Associations were assessed using Spearman rank correlation. Each dot represents an individual participant; the fitted line represents a linear regression with the corresponding 95% confidence interval. Spearman’s correlation coefficient (rho) and two-sided *P* value are indicated on each plot.

## Supplementary information

**Table S1.** All patients (N=156) from FLEMENGHO cohort for serum explorations.

|  | Cancers (n=38) | CVD controls (n=65) | Healthy volunteers (n=53) | p-value* | p-value** |
| --- | --- | --- | --- | --- | --- |
| **Sociodemographic** |  |  |  |  |  |
| **Age**, median (interquartile range) | 65.7 (60.8 ; 74.4) | 67.3 (61.5 ; 75.4) | 63.0 (59.1 ; 66.8) | 0.520 | **0.007** |
| **Sex**, Male, n (%) | 26 (68.4%) | 46 (70.8%) | 31 (58.5%) | 0.827 | 0.179 |
| **Comorbidities** |  |  |  |  |  |
| **Body-mass index (kg/m<sup>2</sup>)**, median (interquartile range) | 26.0 (24.3 ; 29.2) | 27.6 (25.3 ; 28.9) | 25.6 (23.4 ; 29.2) | 0.240 | 0.170 |
| **Tobacco habits** |  |  |  |  |  |
| **Smoking**, Current, n (%) | 20 (52.6%) | 17 (26.2%) | 0 (0.0%) | **0.010** | **<0.001** |
| **Pack-years**, median (interquartile range)<br>(67 NA) | 26.7 (17.3 ; 54.5) | 21.9 (12.1 ; 29.4) | - | 0.196 | - |
| **Smoking duration in years**, median (interquartile range) | 43.2 (33.0 ; 52.3) | 31.0 (21.0 ; 41.9) | - | **0.001** | - |
| **Quit-years**, median (interquartile range) | 0 (0 ; 7.8) | 17.5 (0 ; 30.5) | - | **<0.001** | - |
| **Cancer occurrence** |  |  |  |  |  |
| **Personal history of cancer other than lung cancer**,<br>Yes, n (%) | 3 (7.9%) | 0 (0.0%) | 0 (0.0%) | **0.048** | >0.999 |
| **History of Cardiovascular Disease before examination**,<br>Yes, n (%) | 19 (50.0%) | 53 (81.5%) | 0 (0.0%) | **0.002** | **<0.001** |
| **Time to cancer diagnosis or follow-up (years)**,<br>median (interquartile range) | 4.3 (1.9 ; 6.9) | 7.9 (5.9 ; 11.8) | 4.7 (0.0 ; 8.2) | **0.002** | **<0.001** |

\*Fisher's test: Cancers versus CVD controls
\*\*Fisher's test: Healthy volunteers versus CVD controls

**Table S2.** All patients (N=397) from PREVALUNG cohort for serum explorations.

|  | Cancers (n = 58) | Controls (n = 339) | p-value* | Tobacco-induced Cancers (n = 40) | Other cancers (n = 18) | p-value** |
| --- | --- | --- | --- | --- | --- | --- |
| **Sociodemographic** |  |  |  |  |  |  |
| Age, median (interquartile range) | 69.0 (65.0 ; 72.0) | 64.0 (56.0 ; 69.0) | <0.0001 | 69.0 (65.0 ; 72.8) | 69.0 (60.0 ; 72.0) | 0.4337 |
| Sex, Male, n (%) | 47 (81%) | 281 (82.9%) | 0.730 | 30 (75%) | 17 (94.4%) | 0.146 |
| **Comorbidities** |  |  |  |  |  |  |
| Body-mass index (kg/m<sup>2</sup>), median (interquartile range) | 25.7 (23.1 ; 29.4) | 26.9 (24.2 ; 29.7) | 0.194 | 24.6 (22.1 ; 28.7) | 26.9 (24.1 ; 29.8) | 0.268 |
| Coronary artery disease, n (%) | 36 (62.1%) | 228 (67.3%) | 0.439 | 23 (57.5%) | 13 (72.2%) | 0.285 |
| Peripheral artery disease (PAD), n (%) | 29 (50%) | 120 (35.4%) | 0.034 | 20 (50%) | 9 (50%) | >0.999 |
| Ischemic stroke, n (%) | 9 (15.5%) | 60 (17.7%) | 0.685 | 7 (17.5%) | 2 (11.1%) | 0.706 |
| Aortic aneurysm, n (%) | 9 (15.5%) | 33 (9.7%) | 0.186 | 7 (17.5%) | 2 (11.1%) | 0.706 |
| Upper extremity and visceral artery disease, n (%) | 1 (1.7%) | 7 (2.1%) | >0.999 | 1 (2.5%) | 0 (0.0%) | >0.999 |
| COPD, Yes, n (%) | 13 (22.4%) | 55 (16.2%) | 0.248 | 8 (20%) | 5 (27.8%) | 0.516 |
| Delay last ACVD before blood sampling, median (interquartile range) (months) | 12.5 (4.8 ; 51.3) | 12.0 (4.0 ; 41.0) | 0.379 | 13.0 (4.0 ; 52.5) | 11.0 (7.0 ; 30.0) | 0.894 |
| **Cancer occurrence** |  |  |  |  |  |  |
| **Diagnosed cancer localization** |  |  |  |  |  |  |
| Lung, n (%) | 26 (44.8%) | - | - | 26 (44.8%) | - | - |
| Bladder, n (%) | 10 (17.2%) | - | - | 10 (17.2%) | - | - |
| Head and neck, n (%) | 4 (6.9%) | - | - | 4 (6.9%) | - | - |
| Prostate, n (%) | 5 (8.6%) | - | - | - | 5 (8.6%) | - |
| Colon, n (%) | 4 (6.9%) | - | - | - | 4 (6.9%) | - |
| Hematologic, n (%) | 3 (5.2%) | - | - | - | 3 (5.2%) | - |
| Pancreas, n (%) | 2 (3.4%) | - | - | - | 2 (3.4%) | - |
| Skin, n (%) | 2 (3.4%) | - | - | - | 2 (3.4%) | - |
| Breast, n (%) | 1 (1.7%) | - | - | - | 1 (1.7%) | - |
| Kidney, n (%) | 1 (1.7%) | - | - | - | 1 (1.7%) | - |
| Personal history of cancer, Yes, n (%) | 15 (25.9%) | 15 (4.4%) | <0.001 | 10 (25%) | 5 (27.8%) | >0.999 |
| Time to cancer diagnosis (years), median (interquartile range) | 7 (0.0 ; 33.8) | - | - | 4 (0.0 ; 26.8) | 11 (6.5 ; 39.3) | 0.140 |
| **Tobacco habits** |  |  |  |  |  |  |
| Smoking, Current, n (%) | 22 (37.9%) | 131 (38.6%) | 0.918 | 16 (40.0%) | 6 (33.3%) | 0.628 |
| Smoking duration in years, median (interquartile range) | 44 (36.25 ; 49.75) | 39 (31.5 ; 46) | 0.007 | 45 (38 ; 51) | 39.5 (35 ; 45.5) | 0.110 |
| Quit-years, median (interquartile range) | 13.5 (4.0 ; 21.3) | 8.0 (2.0 ; 19.0) | 0.099 | 11.5 (4.0 ; 20.3) | 16.0 (6.3 ; 25.0) | 0.491 |
| Pack-years, median (interquartile range) | 40.5 (26.0 ; 51.5) | 35.0 (23.0 ; 46.0) | 0.040 | 40.0 (25.0 ; 50.0) | 42.5 (30.0 ; 52.0) | 0.501 |
| NELSON criteria, Yes, n (%) | 38 (65.5%) | 181 (53.7%) | 0.095 | 28 (70.0%) | 10 (55.6%) | 0.284 |
| NLST criteria, Yes, n (%) | 31 (54.4%) | 157 (46.6%) | 0.276 | 21 (53.8%) | 10 (55.6%) | 0.904 |
| PLCOm2012 score, n (%) | 3.0 (1.2 ; 6.2) | 1.5 (0.8 ; 3.2) | <0.001 | 3.0 (1.2 ; 7.1) | 3.0 (1.2 ; 3.7) | 0.395 |
| **Death** |  |  |  |  |  |  |
| Death, Yes, n (%) | 17 (29.3%) | 19 (5.6%) | <0.001 | 12 (30%) | 5 (27.8%) | 0.863 |
| Delay after sampling (years), median (interquartile range) | 3 (1 ; 4) | 2 (1 ; 3) | 0.303 | 3 (1 ; 4) | 3 (2 ; 4) | 0.628 |
| Time to follow-up (years), median (interquartile range) | 4 (3 ; 4.8) | 4 (3 ; 4) | 0.873 | 4 (2 ; 5) | 4 (3.25 ; 4) | 0.773 |

FU: follow-up
\*Wilcoxon rank-sum; Fisher's exact or Chi-square test: Cancer versus Control
\*\*Wilcoxon rank-sum; Fisher's exact or Chi-square test: Tobacco-induced Cancer versus Other cancer

**Table S3.** Exact binomial intervals for each operating characteristic and performance in FLEMENGHO and PREVALUNG cohorts.

| Cohort | Score | Threshold | AUC (95% CI) | Sensitivity (95% CI) | Specificity (95% CI) | PPV (95% CI) | NPV (95% CI) |
| --- | --- | --- | --- | --- | --- | --- | --- |
| FLEMENGHO | Rule-out | ≥26.13% | 0.924 (0.883–0.965) | 0.921 (0.786–0.983) | 0.780 (0.694–0.851) | 0.574 (0.441–0.700) | 0.968 (0.910–0.993) |
| FLEMENGHO | Rule-in | ≥68.73% | 0.789 (0.677–0.901) | 0.429 (0.263–0.606) | 0.885 (0.698–0.976) | 0.833 (0.586–0.964) | 0.535 (0.377–0.688) |
| FLEMENGHO | Combined | ≥26.13% & ≥68.73% | 0.909 (0.852–0.966) | 0.395 (0.240–0.566) | 0.975 (0.927–0.995) | 0.833 (0.586–0.964) | 0.833 (0.760–0.891) |
| PREVALUNG | Rule-out | ≥26.13% | 0.672 (0.600–0.744) | 0.983 (0.908–1.000) | 0.242 (0.197–0.291) | 0.182 (0.140–0.229) | 0.988 (0.935–1.000) |
| PREVALUNG | Rule-in | ≥68.73% | 0.722 (0.647–0.797) | 0.263 (0.155–0.397) | 0.949 (0.915–0.973) | 0.536 (0.339–0.725) | 0.853 (0.807–0.892) |
| PREVALUNG | Combined | ≥26.13% & ≥68.73% | 0.778 (0.716–0.839) | 0.259 (0.153–0.390) | 0.962 (0.935–0.979) | 0.536 (0.339–0.725) | 0.883 (0.846–0.914) |
| PREVALUNG LUNGONLY | Rule-out | ≥26.13% | 0.645 (0.531–0.76) | 0.962 (0.804–0.999) | 0.242 (0.197–0.291) | 0.089 (0.058–0.128) | 0.988 (0.935–1.000) |
| PREVALUNG LUNGONLY | Rule-in | ≥68.73% | 0.805 (0.716–0.895) | 0.320 (0.149–0.535) | 0.949 (0.915–0.973) | 0.381 (0.181–0.616) | 0.935 (0.898–0.962) |
| PREVALUNG LUNGONLY | Combined | ≥26.13% & ≥68.73% | 0.824 (0.739–0.91) | 0.308 (0.143–0.518) | 0.962 (0.935–0.979) | 0.381 (0.181–0.616) | 0.948 (0.919–0.969) |

**Table S4.**
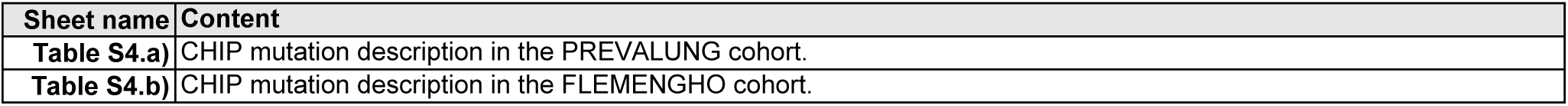
CHIP mutation characteristics in the FLEMENGHO and PREVALUNG cohorts (Fig. 6).

| Sheet name | Content |
| --- | --- |
| Table S4.a) | CHIP mutation description in the PREVALUNG cohort. |
| Table S4.b) | CHIP mutation description in the FLEMENGHO cohort. |

**Table S4.** a) CHIP mutation characteristics in the PREVALUNG cohort.

VAF: variant allele frequency
| Cohort | # | Group | Gene | Reference sequence | Nucleotide change | Amino acid change | VAF (%) |
| --- | --- | --- | --- | --- | --- | --- | --- |
| PREVALUNG | Pt01 | Cancer | *KIT* | NM_000222.2 | c.2866C>T | p.Arg956Trp | 5 |
|  |  |  | *BCORL1* | NM_021946.4 | c.3475A>G | p.Thr1159Ala | 5 |
|  |  |  | *GATA2* | NM_032638.5 | c.988C>T | p.Arg330* | 4 |
| PREVALUNG | Pt02 | Cancer | *KDM6A* | NM_021140.3 | c.3860A>G | p.Lys1315Arg | 9 |
|  |  |  | *EZH2* | NM_004456.5 | c.180G>A | p.Trp60* | 5 |
| PREVALUNG | Pt03 | Cancer | *GATA2* | NM_032638.5 | c.1000A>G | p.K334E | 2 |
|  |  |  | *BCOR* | NM_017745.6 | c.5057A>G | p.N1686S | 2 |
| PREVALUNG | Pt04 | Cancer | *ASXL1* | NM_015338.6 | c.1934dupG | p.G646fsX12 | 19 |
| PREVALUNG | Pt05 | Cancer | *DNMT3A* | NM_022552.5 | c.2645G>A | p.R882H | 5 |
| PREVALUNG | Pt06 | Control | *SF3B1* | NM_012433.3 | c.1997A>G | p.K666R | 2 |
| PREVALUNG | Pt07 | Control | *DNMT3A* | NM_022552.5 | c.2645G>A | p.R882H | 2 |
| PREVALUNG | Pt08 | Control | *CSF3R* | NM_156039.3 | c.2062C>T | p.Q688* | 3 |
| PREVALUNG | Pt09 | Control | *PTPN11* | NM_002834.4 | c.824A>G | p.N275S | 3 |
| PREVALUNG | Pt10 | Control | *EZH2* | NM_004456.5 | c.2015T>C | p.Phe672Ser | 2 |
|  |  |  | *BCORL1* | NM_021946.4 | c.130C>T | p.Gln44* | 2 |
| PREVALUNG | Pt11 | Control | *ASXL1* | NM_015338.6 | c.1537G>T | p.Glu513* | 19 |
| PREVALUNG | Pt12 | Control | *DNMT3A* | NM_022552.5 | c.2332G>A | p.V778M | 7 |
| PREVALUNG | Pt13 | Control | *SF3B1* | NM_012433.3 | c.1986C>G | p.H662Q | 7 |
| PREVALUNG | Pt14 | Control | *SF3B1* | NM_012433.3 | c.2816G>A | p.R939H | 4 |
| PREVALUNG | Pt15 | Control | *CEBPA* | NM_004364.4 | c.383delC | P128Rfs*32 | 4 |
| PREVALUNG | Pt16 | Control | *TET2* | NM_001127208.3 | c.748dupA | p.Thr250fs | 3 |
| PREVALUNG | Pt17 | Control | *PPM1D* | NM_003620.4 | c.1563delG | p.Met521fs | 3 |
| PREVALUNG | Pt18 | Control | *PHF6* | NM_032458.3 | c.433A>C | p.S145R | 3 |
| PREVALUNG | Pt19 | Control | *CBL* | NM_005188.4 | c.1103A>G | p.Tyr368Cys | 2 |

**Table S4.** b) CHIP mutation characteristics in the FLEMENGHO cohort.

HV: Healthy volunteers; VAF: variant allele frequency
| Cohort | # | Group | Gene | Reference sequence | Nucleotide change | Amino acid change | VAF (%) |
| --- | --- | --- | --- | --- | --- | --- | --- |
| FLEMENGHO | Pt01 | Cancer | ATM | NM_000051.4 | c.1229T>C | p.Val410Ala | 36 |
| FLEMENGHO | Pt02 | Cancer | JAK2 | NM_004972.4 | c.1849G>T | p.Val617Phe | 24 |
|  |  |  | TET2 | NM_001127208.3 | c.4893_4905del | Tyr1631fs | 3 |
|  |  |  | GNB1 | NM_001282539 | c.171G>T | p.Lys57Asn | 3.6 |
| FLEMENGHO | Pt03 | Cancer | DNMT3A | NM_022552.5 | c.1243C>T | p.Gln415* | 19 |
|  |  |  | TP53 | NM_000546.6 | c.814G>A | p.Val272Met | 9 |
|  |  |  | CHEK2 | NM_001005735 | c.615dupT | p.His206fs | 2 |
|  |  |  | PPM1D | NM_003620.4 | c.1520_1521insGTGT | p.Met508fs | 3.4 |
| FLEMENGHO | Pt04 | Cancer | NF1 | NM_001042492.3 | c.6943C>G | p.Leu2315Val | 10 |
| FLEMENGHO | Pt05 | Cancer | ATM | NM_000051.4 | c.8188C>G | p.Gln2730Glu | 10 |
| FLEMENGHO | Pt06 | Cancer | DNMT3A | NM_022552.5 | c.1743G>A | p.Trp581* | 9.4 |
| FLEMENGHO | Pt07 | Cancer | CBL | NM_005188.4 | c.1193A>C | p.His398Pro | 2.3 |
|  |  |  | DDX41 | NM_016222 | c.1264G>C | p.Val422Leu | 8.2 |
|  |  |  | TP53 | NM_000546.6 | c.358A>G | p.Lys120Glu | 8.7 |
| FLEMENGHO | Pt08 | Cancer | DNMT3A | NM_022552.5 | c.2141C>G | p.Ser714Cys | 6.6 |
| FLEMENGHO | Pt09 | Cancer | DNMT3A | NM_022552.5 | c.780delC | p.Thr261fs | 4.6 |
| FLEMENGHO | Pt10 | Cancer | TP53 | NM_000546.6 | c.584T>C | p.Ile195Thr | 2 |
| FLEMENGHO | Pt11 | Cancer | DNMT3A | NM_022552.5 | c.2339T>C | p.Ile780Thr | 2 |
| FLEMENGHO | Pt12 | Control | DNMT3A | NM_022552.5 | c.2396C>T | p.Pro799Leu | 13 |
| FLEMENGHO | Pt13 | Control | DNMT3A | NM_022552.5 | c.2408+5G>A | splice donor site | 10.8 |
| FLEMENGHO | Pt14 | Control | CUX1 | NM_001202543 | c.3703G>T | p.Glu1235* | 6.4 |
| FLEMENGHO | Pt15 | Control | SUZ12 | NM_015355 | c.1210G>T | p.Glu404* | 6.1 |
| FLEMENGHO | Pt16 | Control | TP53 | NM_000546.6 | c.473G>A | p.Arg158His | 4.2 |
| FLEMENGHO | Pt17 | Control | PHF6 | NM_001015877 | c.715C>A | p.His239Asn | 3.4 |
|  |  |  | KDM6A | NM_021140 | c.1804G>T | p.Gly602* | 2.6 |
| FLEMENGHO | Pt18 | Control | WT1 | NM_024424 | c.1300C>A | p.Arg434Ser | 3.1 |
| FLEMENGHO | Pt19 | Control | CHEK2 | NM_001005735 | c.493G>T | p.Glu165* | 2.6 |
| FLEMENGHO | Pt20 | Control | RUNX1 | NM_001754 | c.262G>T | p.Glu88* | 2.3 |
| FLEMENGHO | Pt21 | Control | STAG2 | NM_001042749 | c.124-1G>T | splice acceptor site | 2.2 |
| FLEMENGHO | Pt22 | Control | CUX1 | NM_001202543 | c.481G>T | p.Glu161* | 2.1 |
| FLEMENGHO | Pt23 | Control/HV | TET2 | NM_001127208.3 | c.4546C>T | p.Arg1516* | 56 |
| FLEMENGHO | Pt24 | Control/HV | TET2 | NM_001127208.3 | c.3965delT | p.Leu1322fs | 37.3 |
| FLEMENGHO | Pt25 | Control/HV | STAG2 | NM_001042749 | c.2066C>A | p.Ser689* | 8.4 |
| FLEMENGHO | Pt26 | Control/HV | TET2 | NM_001127208.3 | c.5721G>T | p.Met1907Ile | 4.9 |
|  |  |  | ASXL2 | NM_018263 | c.2326G>T | p.Glu776* | 3.3 |
|  |  |  | CBL | NM_005188.4 | c.1868C>A | p.Ser623* | 4.4 |
|  |  |  | WT1 | NM_024424 | c.772G>T | p.Glu258* | 6.7 |
| FLEMENGHO | Pt27 | Control/HV | TET2 | NM_001127208.3 | c.1905dupA | p.Val636fs | 6.3 |
| FLEMENGHO | Pt28 | Control/HV | NF1 | NM_001042492 | c.5164G>T | p.Glu1722* | 2.3 |
|  |  |  | NF1 | NM_001042492 | c.1312G>T | p.Glu438* | 2.8 |
| FLEMENGHO | Pt29 | Control/HV | CEBPA | NM_004364.4 | c.8C>A | p.Ser3* | 3.7 |
| FLEMENGHO | Pt30 | Control/HV | BCOR | NM_001123385.1 | c.4831G>T | p.Glu1611* | 3.3 |
| FLEMENGHO | Pt31 | Control/HV | TET2 | NM_001127208.3 | c.4138C>A | p.His1380Asn | 2.8 |
| FLEMENGHO | Pt32 | Control/HV | EZH2 | ENST00000483012 | c.434G>T | p.Ter145Leuext* | 2.1 |
|  |  |  | NF1 | NM_001042492 | c.7732G>T | p.Glu2578* | 2.4 |
|  |  |  | TERT | NM_198253 | c.268G>T | p.Glu90* | 2.4 |
| FLEMENGHO | Pt33 | Control/HV | DNMT3A | NM_022552.5 | c.2104G>T | p.Asp702Tyr | 2.3 |
| FLEMENGHO | Pt34 | Control/HV | DNMT3A | NM_022552.5 | c.1417G>T | p.Glu473* | 2.3 |
| FLEMENGHO | Pt35 | Control/HV | ATM | NM_000051.4 | c.7708G>T | p.Glu2570* | 2.3 |

**Table S5.**
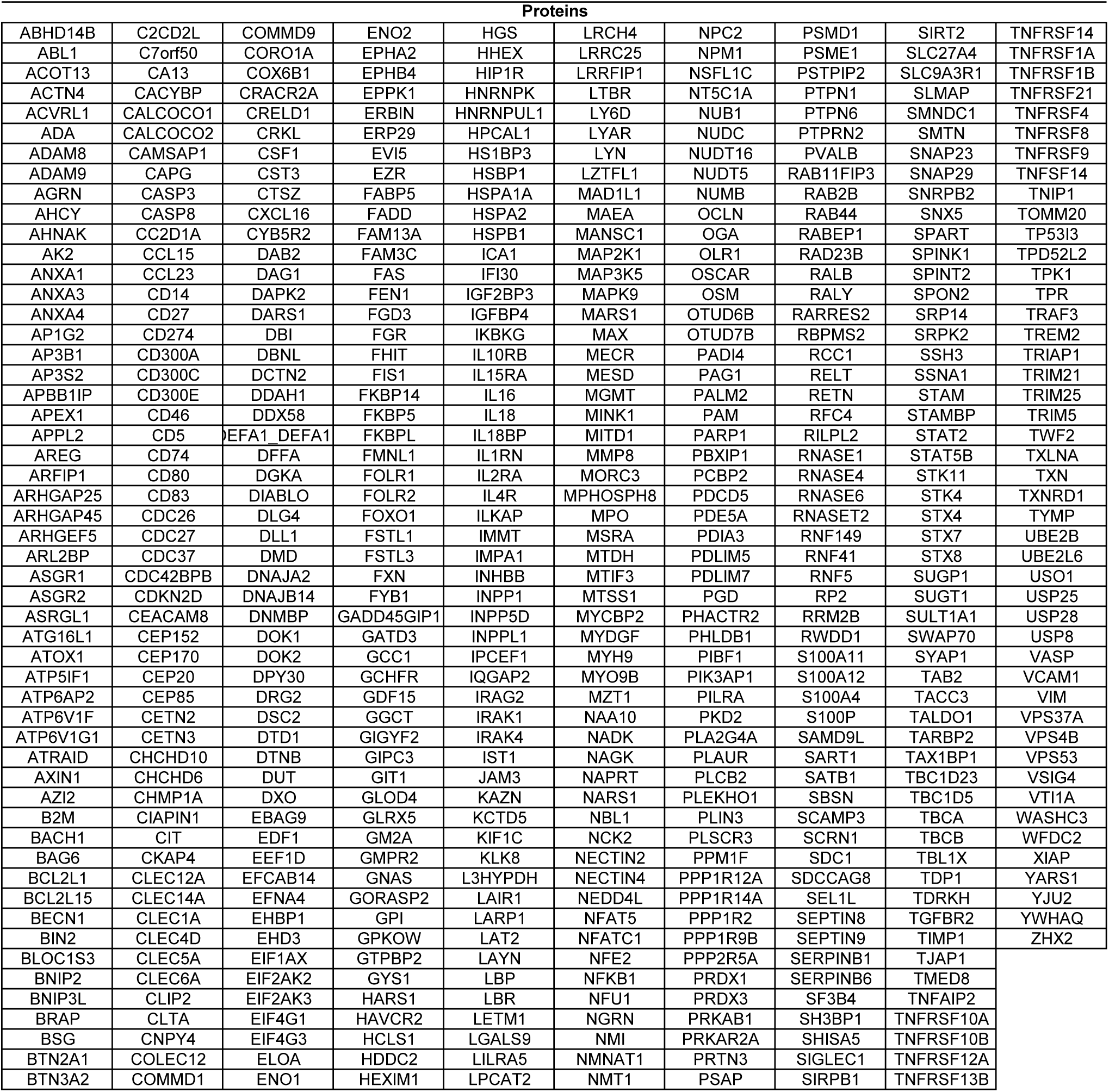
Overlap between FLEMENGHO and PREVALUNG contributive protein lists (n = 523; refer to Fig 4a-b)

**Supplementary Fig. 1.**
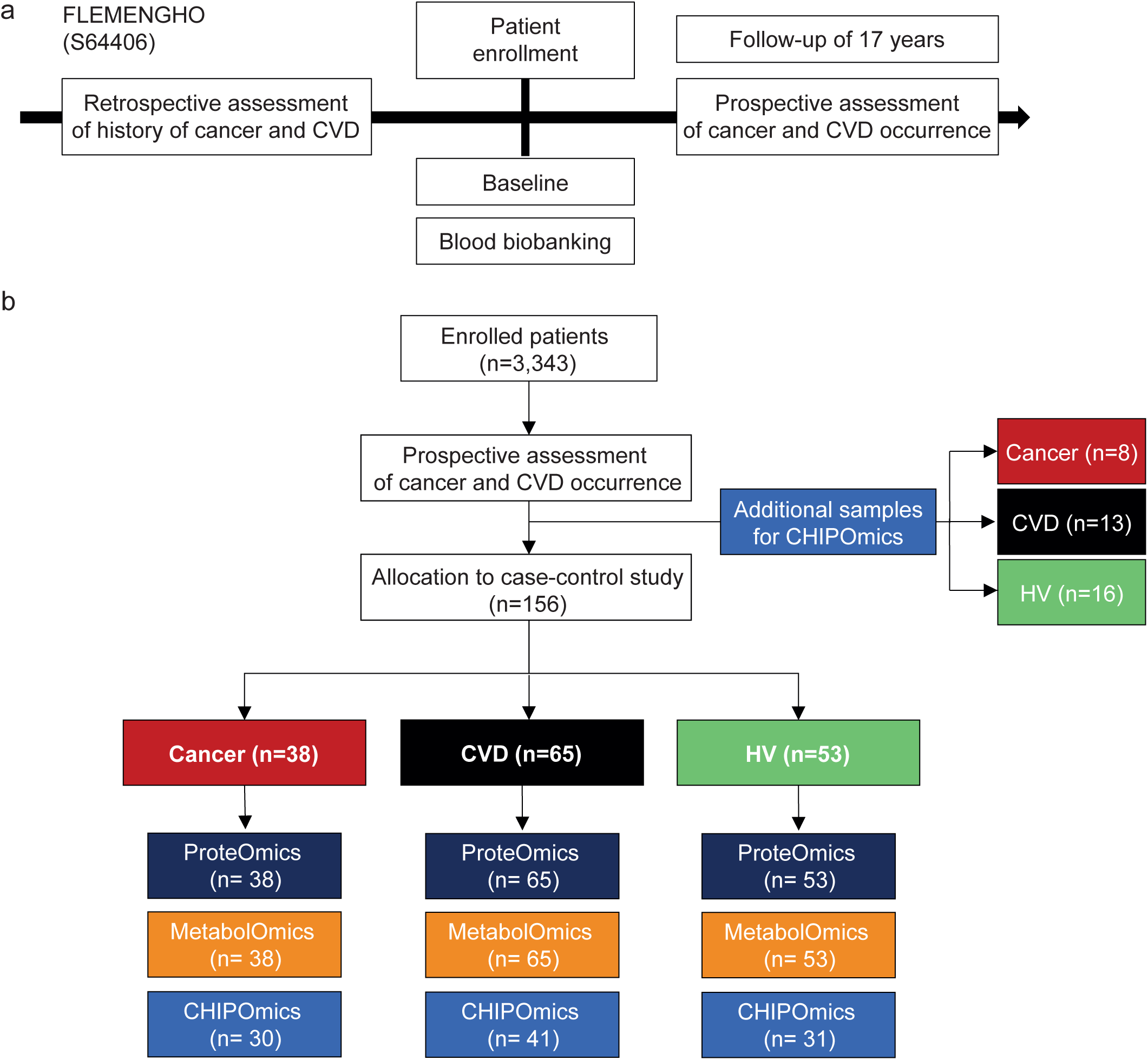
FLEMENGHO cohort consort diagram. **a.** ACVD Flemish patients aged between 60 and 72 years, recorded as current or former smokers were enrolled for a systematic lung cancer and cardiovascular diseases (CVD) screening, and followed up for 1 to 17 years (N=38 CVD cancer, N=65 cancer-free CVD). In parallel, 53 non-smoking healthy volunteers (HV) were paired for age and gender. Plasma and blood were collected 1 to 16 years prior to the diagnosis of lung cancer. **b.** Consort diagram summarizing patient selection, classification, and sampling for ML analysis for each omics. Additional samples of whole blood non selected for the case-control study were used for CHIPOmics.

**Supplementary Fig. 2.**
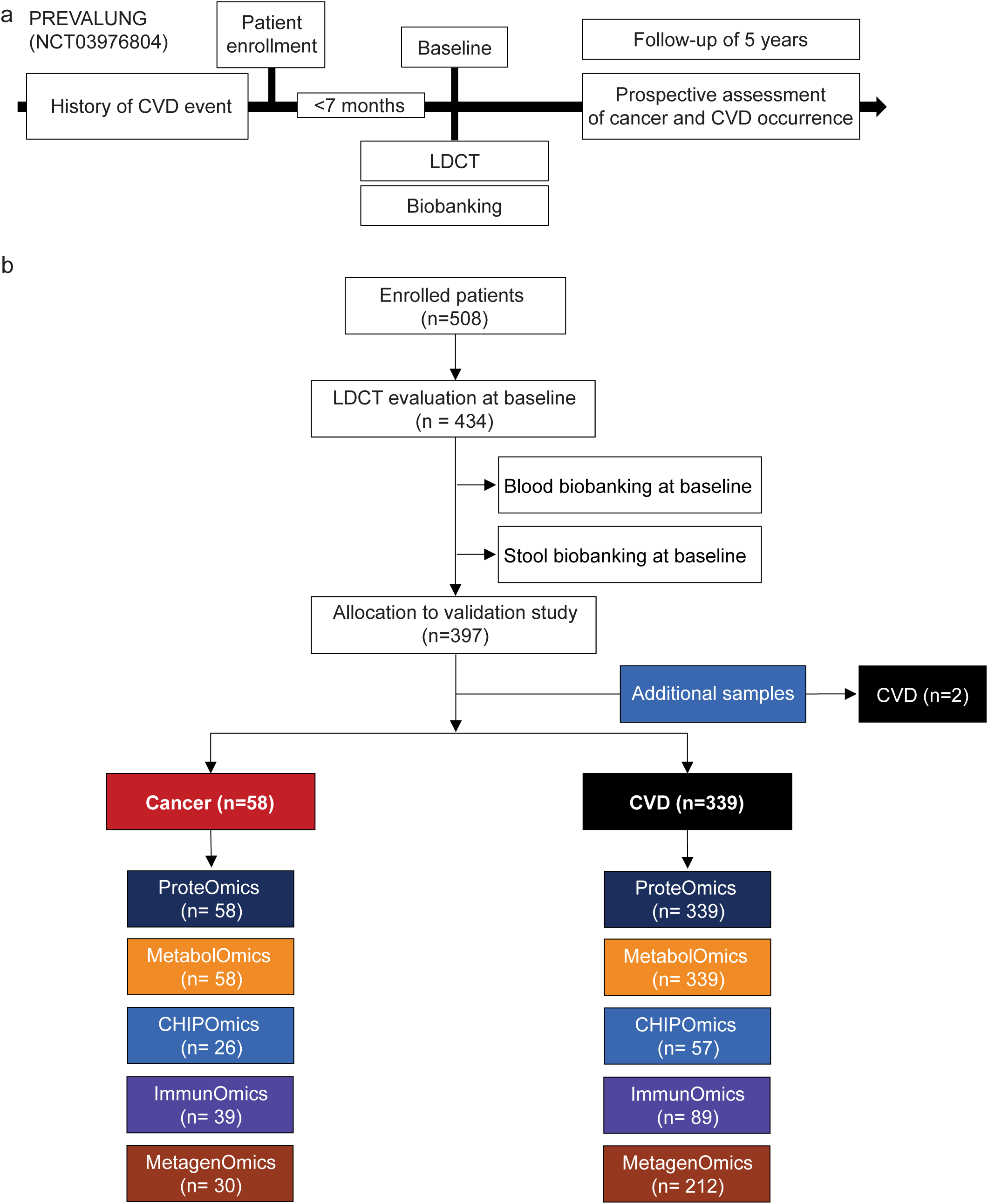
PREVALUNG cohort consort diagram. **a.** Individuals aged between 45 and 75 years, recorded as current or former smokers within the past 10 years and with a diagnosis of ACVD, were enrolled for a systematic Lung Cancer (LC) screening with Low-Dose Computed Tomography (LDCT). Serum, plasma, peripheral blood mononuclear cells (PBMC), and stool samples were collected at day of the LDCT conducted within 7 months after patient enrolment. Patients have been followed up to 5 years to assess asymptomatic cancer diagnosis or CVD events. **b.** Consort diagram summarizing patient selection, classification, and sampling for ML analysis for each omics. Additional samples of PBMCs and non-selected for the validation study were used for CHIPOmics and ImmunOmics.

**Supplementary Fig. 3.**
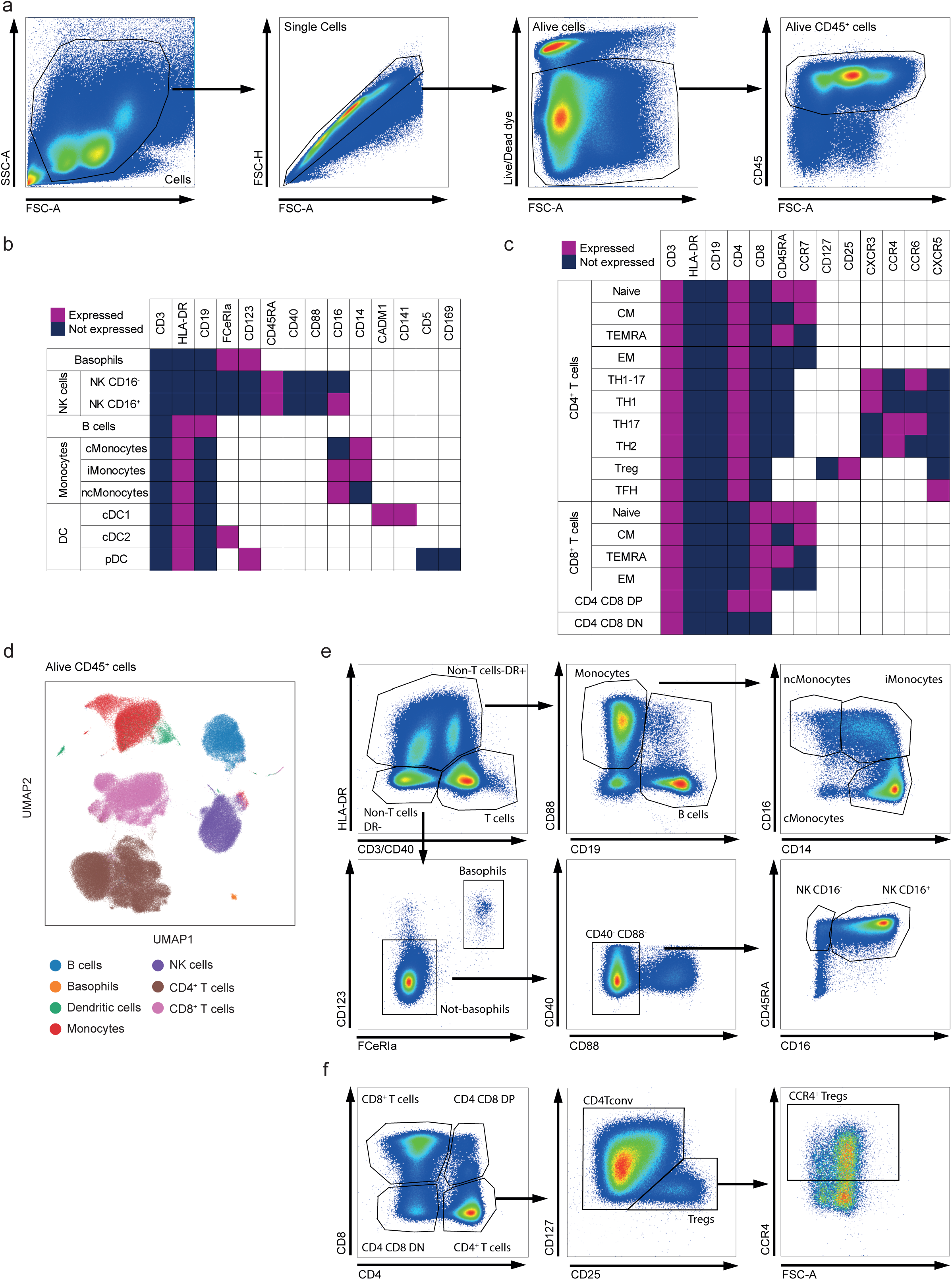
Spectral flow cytometry gating strategies and dimension reduction principles in PREVALUNG cohort. **a.** Flow cytometric analysis of markers in both cancer and control samples was conducted by filtering out debris, doublets and dead cells, and selecting live CD45^+^ cells. **b-d.** Non-linear dimension reduction on cell populations across all patients was applied and displayed through UMAP at different level of analysis (main populations, subpopulations) using automatic annotation with Scyan Python module, based-on expression matrix of characteristic markers for main populations (known to be expressed (purple) or not (blue)) (**b-c**). **e-f.** Manual gating depicts the main populations described in **d**.

**Doc. 1. REMARK reporting checklist.**

**Doc. 2. TRIPOD+AI reporting checklist.**

