## Supplementary material for "Glycometabolic, inflammatory and exocrine plasma proteins predict cancer in individuals with cardiovascular disease": Doc. 1. REMARK reporting checklist.

### INTRODUCTION

1. State the marker examined, the study objectives, and any prespecified hypotheses.

>> The most useful next-generation strategy would combine a parsimonious clinical baseline with a small, mechanistically diverse biomarker set. Cardiovascular phenotype defines an ascertainable risk stratum among smokers. Prospective studies show that cardiovascular disease (CVD) is associated with increased lung-cancer incidence, including among current smokers; severe coronary artery calcification and progression to dyslipidemia are associated with incident lung cancer. **These observations indicate that smokers with cardiovascular risk factors, and particularly those with previous myocardial infarction, stroke, peripheral arterial stenosis or cardiovascular disease requiring revascularization or surgery, are at higher risk than otherwise comparable smokers without such cardiovascular characteristics. We therefore focused on this high-risk population, seeking to establish a convenient and affordable predictive score based on no more than four circulating proteins.** Across the exploration and validation cohorts, we identified two circulating proteins that defined individuals with no or very low risk of future lung-cancer development, as well as two complementary proteins that identified individuals at very high risk. This four-analyte, bidirectional rule-out/rule-in architecture may provide an inexpensive blood-first triage for risk-adapted LDCT.

### MATERIALS AND METHODS

#### Patients

2. Describe the characteristics (e.g., disease stage or comorbidities) of the study patients, including their source and inclusion and exclusion criteria.

>> The "Materials and Methods" section includes a "Cohort Description" subsection, which describes the study population and exclusion and inclusion criteria. References regarding ethics committee approval and clinical trial identification when available (NCT number) are provided for each cohort.

3. Describe treatments received and how chosen (e.g., randomized or rule-based).

>> Not concerned.

>> "Materials and Methods" section includes a "Cohort Description" subsection:

FLEMENGHO is an epidemiological retrospective cohort and represents a random population sample stratified by sex and age from a geographically defined area in northern Belgium.

PREVALUNG is a monocentric and prospective study enrolling adult patients in their follow up post-acute cardiovascular events. Blood and feces samples were collected concomitantly with the CT-scan.

#### Specimen characteristics

4. Describe type of biological material used (including control samples) and methods of preservation and storage.

>> For FLEMENGHO, biobanking is under the supervision of KU Leuven in Belgium. For PREVALUNG, the biobanking is under the supervision of the “Biological Resource Center” from Marie Lannelongue Hospital in France (Research and Innovation Unit, Hôpital Marie Lannelongue, GHPSJ, Le Plessis-Robinson, France).

The "Materials and Methods" section includes a " Method details" subsection, which describes type of biological material used (including control samples) and methods of preservation and storage:

>> For all cohorts, serum and/or plasma, as well as whole blood, when available, were stored at -80°C or lower for long-term storage (FLEMENGHO). Only samples with a good quality for subsequent analysis were selected.

>> For PREVALUNG, peripheral blood mononuclear cells (PBMCs) were isolated using a density gradient medium and stored in liquid nitrogen (-196°C) in a medium composed of 90% fetal bovine serum (FBS) and 10% DMSO.

>> For PREVALUNG, stool aliquots were collected and preserved at -80°C in DNA/RNA Shield Buffer (Zymo Research) until processing.

**Assay methods**

5. Specify the assay method used and provide (or reference) a detailed protocol, including specific reagents or kits used, quality control procedures, reproducibility assessments, quantitation methods, and scoring and reporting protocols. Specify whether and how assays were performed blinded to the study endpoint.

The "Materials and Methods" section includes a " Method details" subsection, which describes the assay method used:

>> For Proteomics: in “Proteomics analysis on serum” subsection
>> For Metabolomics: in “Metabolomics analysis on plasma or serum.” subsection
>> For OGA orthogonal validation: “Orthogonal validation of OGA protein levels in circulating immune cells (PBMCs)” subsection
>> For Immunomics: in “Immunophenotyping by spectral flow cytometry” subsection
>> For CHIP analysis: in “Targeted deep sequencing and CHIP-related mutation analysis” subsection
>> For Metagenomic: in “Collection of patient stools, DNA extraction and sequencing (PREVALUNG).” subsection

**Study design**

6. State the method of case selection, including whether prospective or retrospective and whether stratification or matching (e.g., by stage of disease or age) was used. Specify the time period from which cases were taken, the end of the follow-up period, and the median follow-up time.
>> "Materials and Methods" section includes a "Cohort Description" subsection which states the method of case selection:

>> For FLEMENGHO:
Participants eligible for inclusion in this retrospective validation study were selected from the FLEMENGHO cohort. For 38 selected lung cancer cases the blood samples were of a good quality for proteomics/metabolomics. These were samples obtained from consequently examined participants starting from 1996 till 2015. Median follow-up time between examination/ blood sampling and incidence of lung cancer is 4.3 years (interquartile range, 1.9 to 6.9 years). We matched these cases by age, sex, body mass index (BMI), with 53 healthy non-smoker controls from the same population. We also included 65 subjects with ACVD (atherosclerosis events) who did not develop lung cancer (or any other forms of cancer) during follow-up period (4.9 years). These ACVD controls have been paired on age, gender, BMI, and by tobacco use (current or ex-smokers) with lung cancer cases

>> For PREVALUNG:
PREVALUNG study is a monocentric and prospective study enrolling adult patients in their follow up post-acute cardiovascular events. The inclusion criteria were (1) age 45-75 years old, (2) history of atheromatous cardiovascular disease (ACVD) and (3) history of daily tobacco consumption for 10 years prior to onset of ACVD. Exclusion criteria were symptoms of lung cancer (LC), existing follow-up for pulmonary nodule, fibrosis, pulmonary hypertension, resting dyspnea and active pulmonary infectious disease. 508 patients were included in this study from November 18th, 2019 up to May 18th, 2021 at Marie Lannelongue hospital (Le Plessis-Robinson, France).
Proteomics data were available for 397 patients and clinical follow-up for cancer diagnosis and ACVD events was up to 5 years after inclusion with a median follow-up time of 4 years.

7. Precisely define all clinical endpoints examined.

>> "Materials and Methods" section includes a "Cohort Description" subsection which defines all clinical endpoints examined:

In brief: Outcome = any incident cancer during follow-up (Control vs Cancer). FLEMENGHO development outcome was incident lung cancer; PREVALUNG validation outcome was all incident cancers (26 lung, 14 other tobacco-related, 18 non-tobacco-related). Also in Methods: Data curation.

>> For FLEMENGHO:
Participants eligible for inclusion in this retrospective validation study were selected from the FLEMENGHO cohort. For 38 selected lung cancer cases the blood samples were of a good quality for proteomics/metabolomics. These were samples obtained from consequently examined participants starting from 1996 till 2015. Median follow-up time between examination/ blood sampling and incidence of lung cancer is 4.3 years (interquartile range, 1.9 to 6.9 years). We matched these cases by age, sex, body mass index (BMI), with 53 healthy non-smoker controls from the same population. We also included 65 subjects with ACVD (atherosclerosis events) who did not develop lung cancer (or any other forms of cancer) during follow-up period (4.9 years). These ACVD

controls have been paired on age, gender, BMI, and by tobacco use (current or ex-smokers) with lung cancer cases.

>> For PREVALUNG:
Blood and feces samples were collected concomitantly with the CT-scan. Each patient was scheduled for a follow-up by phone visits at month 3, month 6 and month 12 after the CT-scan. Lung cancer prevalence was the primary endpoint. National Lung Screening Trial and Dutch-Belgian Randomized Lung Cancer Screening Trial (NELSON) trial eligibility criteria, radiation, positive screening, false positivity, rate of localized LC diagnosis, quality of life with the Short Form 12 (SF-12) and anxiety with the Spielberger State-Trait Anxiety Inventory A and B (STAI-YA and STAI-YB, respectively), smoking cessation and onset of cardiovascular and oncological events within 1 year of follow-up were recorded. Proteomics data were available for 397 patients and clinical follow-up for cancer diagnosis and ACVD events was up to 5 years after inclusion with a median follow-up time of 4 years.

8. List all candidate variables initially examined or considered for inclusion in models.
>> "Materials and Methods" section includes a " Statistical analysis" subsection which lists all candidate variables initially examined or considered for inclusion in models (“Data curation and standardization” subsection)

9. Give rationale for sample size; if the study was designed to detect a specified effect size, give the target power and effect size.

>> "Materials and Methods" section includes a "Cohort Description" subsection which gives rationale for sample size:

>> For FLEMENGHO:
The participants (n=3,343) were repeatedly examined at a local examination center. During regular re-examination, standardized and validated questionnaires were administered to collect detailed information about each participant’s medical history, use of medication, socio-economic class and lifestyle including information on smoking habit. Phenotyping at the examination center included clinical and anthropometric measurements. Blood samples (i.e. serum) are available. Every year, the vital status of all registered persons and any changes in their place of residence are requested from the National Register in Brussels. The competent service of the Flemish Community then provides the immediate and underlying causes of death, as registered on the death form. To confirm the diagnoses on the questionnaires (non-fatal endpoints) or on the death certificate, the study team contacts the GP or the treating specialist, provided that the participants gave their prior consent. Participants eligible for inclusion in this retrospective validation study were selected from the FLEMENGHO cohort. **For 38 selected lung cancer cases the blood samples were of a good quality for proteomics/metabolomics.** These were samples obtained from consequently examined participants starting from 1996 till 2015. Median follow-up time between examination/ blood sampling and incidence of lung cancer is 4.3 years (interquartile range, 1.9 to 6.9 years). **We matched these cases by age, sex, body**

**mass index (BMI), with 53 healthy non-smoker controls from the same population. We also included 65 subjects with ACVD (atherosclerosis events) who did not develop lung cancer (or any other forms of cancer) during follow-up period (4.9 years). These ACVD controls have been paired on age, gender, BMI, and by tobacco use (current or ex-smokers) with lung cancer cases.**

>> For PREVALUNG:
**508 patients were included in this study from** November 18th, 2019 up to May 18th, 2021 at Marie Lannelongue hospital (Le Plessis-Robinson, France). **Proteomics data were available for 397 patients only** and clinical follow-up for cancer diagnosis and ACVD events was up to 5 years after inclusion with a median follow-up time of 4 years.

**Statistical analysis methods**
10. Specify all statistical methods, including details of any variable selection procedures and other model-building issues, how model assumptions were verified, and how missing data were handled.

>> "Materials and Methods" section includes a " Statistical analysis" subsection which specifies all statistical methods.

>> In addition, the study is reported per TRIPOD+AI; the completed checklist is supplied as a Supplementary file.

11. Clarify how marker values were handled in the analyses; if relevant, describe methods used for cutpoint determination.

>> "Materials and Methods" section includes a " Statistical analysis" subsection which clarifies how marker values were handled, especially in the “Machine-learning signature” subsection.

>> In addition, the study is reported per TRIPOD+AI; the completed checklist is supplied as a Supplementary file.

**RESULTS**
**Data**
12. Describe the flow of patients through the study, including the number of patients included in each stage of the analysis (a diagram may be helpful) and reasons for dropout. Specifically, both overall and for each subgroup extensively examined report the numbers of patients and the number of events.

>> For FLEMENGHO: flow chart in a **Supplementary figure 1**.
>> For PREVALUNG: flow chart in a **Supplementary figure 2**.

13. Report distributions of basic demographic characteristics (at least age and sex), standard (disease-specific) prognostic variables, and tumor marker, including numbers of missing values.

>> basic demographic characteristics:
For FLEMENGHO: in **Supplementary Table 1**
For PREVALUNG: in **Supplementary Table 2**

>> CHIP mutations analysis:
For FLEMENGHO: in **Supplementary table S4b**
For PREVALUNG: in **Supplementary table S4a**

**Analysis and presentation**
14. Show the relation of the marker to standard prognostic variables.

>> In the **Extended Data Figure 8**, AUC and Kaplan-Meier curves are presented for previously published scores (PLCO) or lung cancer screening inclusion criteria (NELSON and NLST studies); these can be compared with the AUC and KM curves associated with the rule-out and rule-in scores **-** as shown in **Figure 6 -** and with the exact binomial intervals for each operating and performance characteristic in the FLEMENGHO and PREVALUNG cohorts described in **Table S3**.

>> In addition, in the **Extended Data Figure 5 and 7**, volcano plots display proteins associated with various previously published cancer risk signatures, categorized by the rule-out and the rule-in classification in individuals from FLEMENGHO and PREVALUNG cohorts.

15. Present univariate analyses showing the relation between the marker and outcome, with the estimated effect (e.g., hazard ratio and survival probability). Preferably provide similar analyses for all other variables being analyzed. For the effect of a tumor marker on a time-to-event outcome, a Kaplan – Meier plot is recommended.

>> The rule-out and rule-in scores and their combination were studied in each cohort with ROC curves generated using the pROC (v1.19.0.1) package and Kaplan-Meier curves and Cox proportional hazards regression analyses were performed using the survival (v3.8-6) and survminer (v0.5.2) packages.

>> Results are depicted in Figure 6 and exact binomial intervals for each operating and performance characteristic in the FLEMENGHO and PREVALUNG cohorts are described in **Table S3**.

16. For key multivariable analyses, report estimated effects (e.g., hazard ratio) with confidence intervals for the marker and, at least for the final model, all other variables in the model.
>> Not concerned.

17. Among reported results, provide estimated effects with confidence intervals from an analysis in which the marker and standard prognostic variables are included, regardless of their statistical significance.

>> In the Extended Data Figure 8, AUC and Kaplan-Meier curves are presented for previously published scores (PLCO) or lung cancer screening inclusion criteria (NELSON and NLST studies); these can be compared with the AUC and KM curves associated with the rule-out and rule-in scores - as shown in Figure 6 - and with the exact binomial intervals for each operating and performance characteristic in the FLEMENGHO and PREVALUNG cohorts described in Table S3.

>> In addition, in the Extended Data Figure 5 and 7, volcano plots display proteins associated with various previously published cancer risk signatures, categorized by the rule-out and the rule-in classification in individuals from FLEMENGHO and PREVALUNG cohorts.

18. If done, report results of further investigations, such as checking assumptions, sensitivity analyses, and internal validation.
>> The study is reported per TRIPOD+AI; the completed checklist is supplied as a Supplementary file.

**DISCUSSION**

19. Interpret the results in the context of the prespecified hypotheses and other relevant studies; include a discussion of limitations of the study.
>> done.

20. Discuss implications for future research and clinical value.

>> done.
