## Supplementary material for "Glycometabolic, inflammatory and exocrine plasma proteins predict cancer in individuals with cardiovascular disease": Doc. 2. TRIPOD+AI reporting checklist.

### Methods

| Item | Section | Checklist item | Where reported / how addressed |
|---|---|---|---|
| **4a** | Source of data | Describe the study design and key dates (accrual, prediction horizon). | FLEMENGHO: population-based Belgian cohort, samples 1996–2015, median follow-up 4.3 y. PREVALUNG: prospective French cohort, enrolled 18 Nov 2019–18 May 2021, LDCT within 7 months, follow-up to 5 y. Methods: Cohort description. |
| **4b** | Source of data | Specify whether development and validation data came from the same or different sources. | Different sources: development on FLEMENGHO (Leuven), transfer to PREVALUNG (Le Plessis-Robinson). Different countries, eras, storage and assay batches; stated in Discussion (limitations). |
| **5a** | Participants | Specify eligibility criteria and setting. | PREVALUNG inclusion: age 45–75, atheromatous CVD, ≥10 y daily tobacco before ACVD; exclusions listed (existing nodule follow-up, fibrosis, PH, resting dyspnoea, active infection). FLEMENGHO: nested case–control (38 lung-cancer cases, 53 healthy non-smoker controls, 65 ACVD controls). Methods: Cohort description. |
| **5b** | Participants | Describe treatments/interventions received, if relevant. | Not an intervention study. PREVALUNG participants received a scheduled LDCT within 7 months of sampling; no treatment allocation. Confounding by CVD medications discussed in the Discussion (limitations). |
| **6a** | Outcome | Define the outcome that is predicted, including how and when assessed. | Outcome = any incident cancer during follow-up (Control vs Cancer). FLEMENGHO development outcome was incident lung cancer; PREVALUNG validation outcome was all incident cancers (26 lung, 14 other tobacco-related, 18 non-tobacco-related). Methods: Data curation. |
| **6b** | Outcome | Report any actions to blind assessment of the outcome. | Model preprocessing constants, classifiers and both thresholds were frozen on FLEMENGHO before PREVALUNG outcomes were examined; outcome ascertainment used registry/clinical follow-up independent of the score. Methods: Stage 3 / validation. |
| **7a** | Predictors | Define predictors, including how and when measured. | Predictors: OGA (OGA_Oncology@II), IL-6 (IL6_Oncology), CTRC (CTRC_Inflammation), MSMB (MSMB_Cardiometabolic) — Olink Explore 3072 NPX, single serum draw at inclusion — plus smoking status and personal cancer history at sampling. Methods: Proteomics analysis; Data curation. |

| Item | Section | Checklist item | Where reported / how addressed |
|---|---|---|---|
| **7b** | Predictors | Report any actions to blind predictor assessment. | Olink NPX generated by the assay provider under standard normalisation/QC, blind to outcome; feature selection and thresholds derived before PREVALUNG outcomes were unblinded. Methods: Proteomics; Stage 3. |
| **8** | Sample size | Explain how the study size was arrived at. | Sample sizes were fixed by the available biobank cases with adequate sample quality (FLEMENGHO 38 cases; PREVALUNG 397 with proteomics). The modest size and its consequences (wide intervals, especially the 28-participant high-risk stratum) are acknowledged in the Discussion. |
| **9** | Missing data | Describe how missing data were handled. | Missing NPX values imputed with FLEMENGHO training-set medians; the same constants applied to PREVALUNG. Methods: Data curation and standardisation. |
| **10a** | Analysis — development | Describe how predictors were handled in the analysis. | Continuous NPX standardised with a FLEMENGHO-fit StandardScaler; smoking status mapped to 0/1/2; cancer history binary. 2,910 QC-passing proteins entered feature selection. Methods: Data curation; Machine-learning signature. |
| **10b** | Analysis — development | Specify type of model, model-building procedure, and validation (internal). | Three-stage discovery pipeline: (1) consensus multivariate ranking over 1,000 bootstraps (LASSO stability, RF permutation, TreeSHAP, ridge coefficients, LightGBM gain); (2) mRMR (MID) shortlisting; (3) classifier-coupled backward elimination over repeated stratified 80/20 splits, selecting a soft-voting GaussianNB + L2 logistic ensemble. Run twice to build the rule-out and rule-in arms. Internal evaluation by 5-fold CV. Scripts 01–04. |
| **10c** | Analysis — AI specifics | Report the AI/ML learning algorithm details sufficient for reproduction (architecture, hyperparameters, software). | Locked ensemble: GaussianNB (class-balanced sample weights) + LogisticRegression (C=1.0, L2, class_weight='balanced', lbfgs, max_iter=2000), soft-voting mean probability. Preprocessing, coefficients and thresholds deposited as locked_classifier.json. Implemented in Python/scikit-learn; versions pinned in requirements.txt. Scripts fully deposited. |
| **10d** | Analysis | Specify how risk groups / thresholds were created, if done. | Two thresholds: rule-out = lowest score at sensitivity ≥ 0.925 (FLEMENGHO); rule-in = lowest score at sensitivity ≥ 0.40 among non-cleared. Manuscript values 26.13 %/68.73 % are means of 100 bootstrap-derived thresholds. Three strata: rule-out, medium, high. Methods: Stage 3; README note on thresholds. |

| Item | Section | Checklist item | Where reported / how addressed |
| --- | --- | --- | --- |
| **10e** | Analysis — validation | Describe how the model was applied to validation data. | The locked pipeline (medians, scaler, classifiers, stage assignments, both thresholds) was applied to PREVALUNG without refitting or recalibration. Script 04; Methods: Signature validation on PREVALUNG. |
| **11** | Risk groups | Provide details on how risk groups were defined, if done. | Rule-out (very low risk), medium risk, high risk, defined by the two thresholds as above. Labels name classifier arms, not clinical decisions (stated explicitly in Results). |
| **12** | Development vs validation | For validation, identify any differences from development (setting, eligibility, outcome, predictors). | Development outcome was incident lung cancer; validation outcome was all incident cancers. Cohorts differ in country, era, storage and assay batch; preprocessing constants transferred without a bridging study. Discussion (limitations). |
